# Metabolic disease-relevant stimuli unmask context-dependent genetic regulation of cardiometabolic loci in human adipocytes

**DOI:** 10.64898/2026.08.12.744187

**Authors:** Yi Huang, Joaquin Perez-Schindler, Sanchari Datta, Yijia Christiana Liu, Bellis Min, Mayank Murali, Renee M. Gibson, Adam G. Maynard, Nivedita Nambrath, Phil Kubitz, Sunita V. Singh Poma, Wei-Lin Qiu, Kyra Henriques, Thiago M. Batista, Bandana Sharma, Thouis R. Jones, Robin Andersson, Hesam Dashti, Cornelia Griggs, Ben Neale, Wei Zhou, Melina Claussnitzer

**Author notes:** These authors contributed equally to this work (Yi Huang, Joaquin Perez-Schindler).

## Abstract

Genome-wide association studies (GWAS) have identified thousands of loci associated with cardiometabolic disease, yet translating these associations into regulatory mechanisms, effector genes, and cellular programs remains a major challenge. A key limitation is that genetic effects are often highly context dependent, varying across cell states and environmental conditions that are difficult to model at scale. Here, we leverage *CellGenBank*, a population-scale biobank of primary human adipose-derived mesenchymal stem cells (AMSCs), to implement a multi-donor cell village *in vitro* system to map cardiometabolic disease genetic variation across adipocyte differentiation and metabolic stress conditions.

We pooled AMSCs from 118 donors into multiplexed villages, differentiated them toward adipocytes, and profiled chromatin accessibility and gene expression using single-nucleus multiome sequencing under four disease-relevant conditions: basal, elevated free fatty acids, low glucose, and hypoxia. By combining with genetic demultiplexing, we quantify how regulatory element activity, gene expression, and higher-order cellular programs are modulated by both genotype and environmental context.

Across conditions, we identify widespread context-specific cis-regulatory effects, including expression and chromatin accessibility quantitative trait loci that are masked in baseline states. Genetic effects frequently converge on coordinated transcriptional programs linked to lipid metabolism, insulin responsiveness, and stress adaptation, enabling the identification of cellular program QTLs that bridge variants, genes, and disease-relevant phenotypes. Integration with cardiometabolic GWAS reveals enhanced colocalization in condition- and state-resolved analyses, highlighting the importance of modeling environmental context to resolve disease mechanisms.

Together, our study establishes large-scale adipocyte cell villages as a powerful and generalizable framework to map the context-dependent regulatory architecture of cardiometabolic disease and provides a resource linking human genetic variation to adipocyte cellular programs.

## Introduction

Large-scale genome-wide association studies (GWAS) have transformed our understanding of the genetic architecture of cardiometabolic diseases such as type 2 diabetes, obesity, and dyslipidemia^1–3^. The majority of associated variants reside in non-coding regions of the genome^4–6^, implicating gene regulation as a central driver of polygenic disease risk^7–14^. However, connecting these variants to causal regulatory elements, effector genes, and pathogenic cellular mechanisms at scale remains a fundamental challenge.

The Variant-to-Function (V2F) framework is a multidisciplinary strategy for characterizing how GWAS-identified genetic variants functionally impact regulatory elements, target genes, cell states, and cellular programs to inform disease mechanisms^7–13^. A major obstacle in V2F efforts is context dependence. Genetic effects on gene expression and chromatin accessibility can vary dramatically across cell types, differentiation stages, and environmental conditions, including nutrient availability and metabolic stress^15–18^. Metabolic cell types including adipocytes, hepatocytes, and myocytes are instrumental in maintaining whole-body metabolic homeostasis^15–17^. Among these, adipocytes play a particularly pivotal role, and their dysfunction is increasingly recognized as a key pathogenic driver of metabolic disease. Impaired adipocyte function contributes to systemic insulin resistance, chronic low-grade inflammation, and dysregulated lipid metabolism^18,19^. Furthermore, the functional effects of genetic variants often manifest through altered adipose tissue mass and distribution^20,21^. Recent *in vivo* and *in vitro* studies have also revealed substantial cellular heterogeneity across adipose tissue and adipocyte differentiation^22–25^. As a result, cell state- and context-specific pathogenic mechanisms critical for metabolic disease development may be diluted or entirely missed in bulk or single context data, highlighting the relevance of readouts with single cell resolution^26^.

Recent advances in single-cell genomics and pooled experimental designs have begun to address this challenge by enabling scalable molecular QTL (molQTL) mapping in defined cellular contexts^27–29^. However, scaling these approaches across donors, stimuli, and data modalities in primary human adipocytes has remained difficult. The development of the cell village framework provides a solution by enabling large-scale molQTL mapping in pooled cellular systems. In this approach, cells from genetically distinct donors are pooled, cultured and differentiated together, profiled jointly, and computationally demultiplexed to assign them to their donor of origin, allowing genetic effects to be assessed at single-cell resolution while minimizing technical variability. Cell villages have been successfully applied to induced pluripotent stem cell-derived systems, establishing a foundation for “GWAS in a dish” approaches^30–32^. Whether this framework can be extended to metabolic cells such as primary human adipocytes and leveraged to interrogate stimuli-responsive disease-relevant mechanisms has remained an open question.

Here, we leverage *CellGenBank*, a population-scale biobank of primary human adipose-derived mesenchymal stem cells (AMSCs), to implement cell villages coupled to multimodal single-nucleus genomic readouts. Multi-donor AMSCs provide a genetically diverse, physiologically relevant entry point to study adipocyte biology and cardiometabolic disease mechanisms^33,34^, while enabling controlled differentiation and perturbations *in vitro*. We pooled AMSCs from 118 donors into multiplexed villages, differentiated them toward adipocytes, and exposed them to metabolic disease-relevant stimulatory conditions. Using single-nucleus multiome profiling, we simultaneously measured gene expression and chromatin accessibility across donors, cell states, and stimuli. This design allows us to systematically map how cardiometabolic disease-associated genetic variation influences regulatory elements, genes, and coordinated cellular programs in a context-dependent manner. By integrating *cis*-regulatory QTL mapping with cellular program-level analyses and GWAS colocalization/association, we show that environmental context is a key factor of genetic effect linked to metabolic disease mechanisms in human adipocytes. Our results establish human metabolic primary cell villages as a scalable V2F platform for dissecting the regulatory mechanisms underlying complex metabolic disease.

## Results

### Establishing a human adipocyte single-nucleus Multiome (snMultiome) atlas under metabolic disease-relevant stimuli

To enable large-scale, context-specific genetic mapping in primary human adipocytes, we implemented a cell village framework leveraging *CellGenBank*, a cellular biobank of primary human AMSCs derived from 1,275 donors (see Methods, **Fig. 1a**). Donors were first matched by age, sex, body mass index (BMI), and polygenic risk scores for cardiometabolic traits, then randomly selected to generate an evenly distributed set that ensures broad representation of relevant demographic and genetic factors **(Fig. 1c, Extended Data Fig. 1a-c, Supplementary Table 1)**. The majority of selected donors are of European (EUR) ancestry (90.7%), with genetic principal components (PCs) clustering closely with the EUR superpopulation from the 1000 Genomes reference panel **(Extended Data Fig. 1d)**. AMSCs from 118 genetically distinct donors were pooled into four multiplexed villages, each comprising 20-40 donors, differentiated toward different adipose trajectories, and profiled using snMultiome (snRNA-seq + snATAC-seq) (**Fig. 1a**). Villages were differentiated under four metabolic disease-relevant conditions, namely basal (high glucose normoxia, hereafter referred to as “basal”), elevated free fatty acids (FFA), low glucose, and hypoxia, with nuclei collected at day 0 before induction and day 14 of differentiation with stimuli (**Fig. 1a**). These conditions were selected to model key environmental factors implicated in cardiometabolic disease, including nutrient overload, healthy physiological glucose level, and hypoxic stress^35–40^. To facilitate robust integration across villages and conditions, each village included an internal immortalized AMSC control line derived from a single donor, enabling quantitative assessment of batch effects and integration performance **(Supplementary Notes)**. We constructed a unified adipocyte differentiation and stimulation atlas by integrating snMultiome profiles across villages and conditions (**Fig. 1b**). After genetic demultiplexing using donor genotype information (Dropulation, average doublet/empty droplet rate = 31.64%) and stringent quality control (**Methods**), we obtained an atlas of 337,666 high-quality nuclei for the RNA modality, of which 312,088 nuclei also passed quality thresholds for chromatin accessibility (**Supplementary Table 2**). Nuclei were assigned to their donor of origin and analyzed in a cell state- and stimuli-specific manner to capture context-specific gene regulatory mechanisms. This framework allowed us to systematically interrogate how genetic variation influences chromatin accessibility, gene expression, and coordinated cellular programs across healthy and disease-like states relevant to metabolic diseases.

**Fig. 1.**
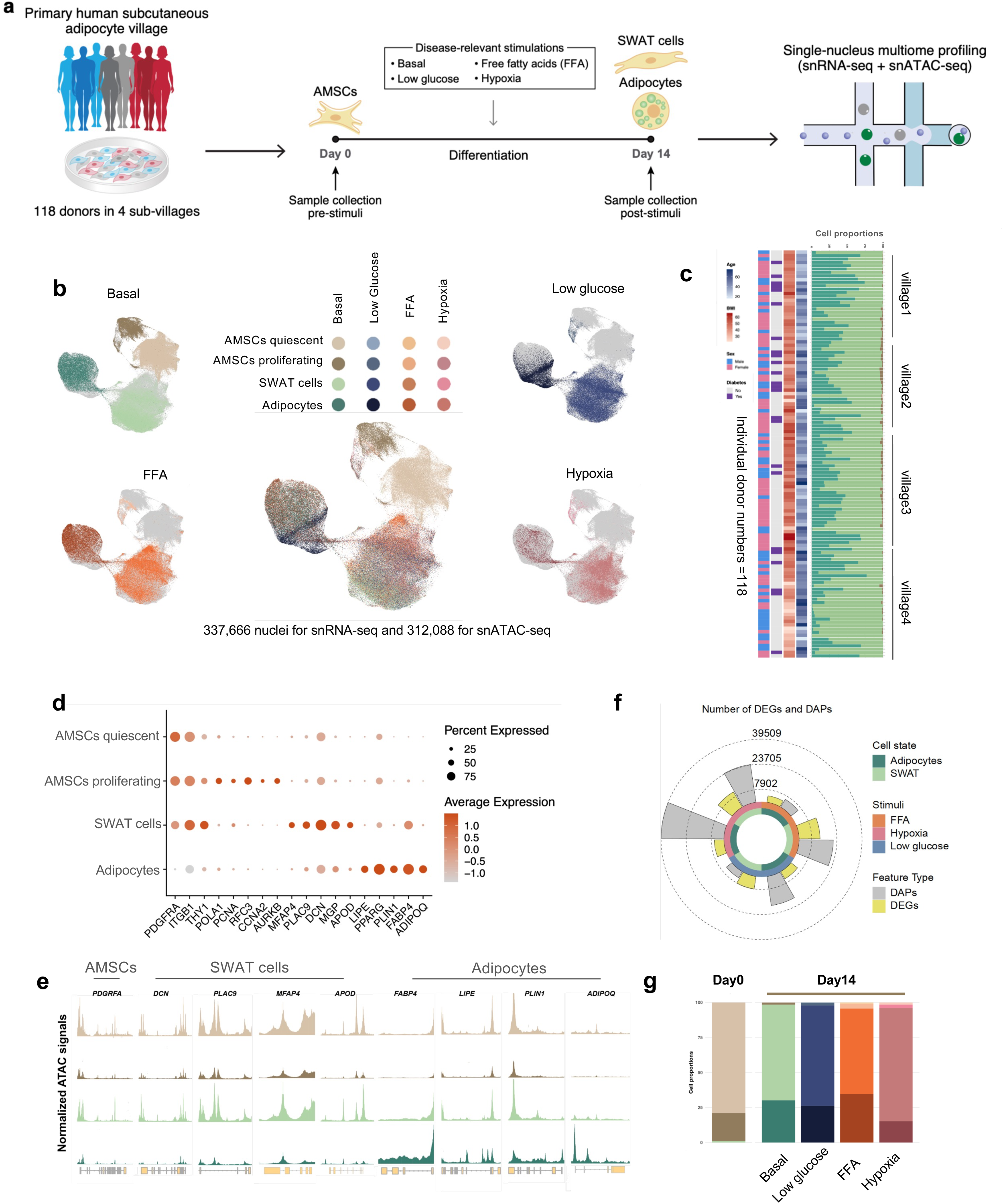
A human adipocyte single-nucleus Multiome map under metabolic disease-relevant stimuli. **a,** Schematic of the study design. **b,** UMAP embedding of the single-nucleus multiome dataset (337,666 nuclei for snRNA-seq and 312,088 for snATAC-seq), showing four annotated cell states across four stimulus conditions. Colors represent cell states and stimuli. Surrounding UMAPs show cells split by stimulus condition (basal, low glucose, FFA, hypoxia), with grey background representing cells not assigned to the given condition. **c,** Per-donor overview across the 118 individual donors, grouped by village (1-4). From left to right, donor metadata (sex, diabetes status, BMI, age) and relative cell-state proportions per donor. **d,** Dot plot of cell-state marker gene expression for basal condition. Dot size indicates the percentage of nuclei expressing each gene and color indicates scaled average expression. **e,** Normalized snATAC-seq accessibility tracks at cell-state marker gene loci, shown for each cell state at basal condition. **f,** Number of differentially expressed genes (DEGs) and differentially accessible peaks (DAPs) per cell state (adipocytes, SWAT) and stimulus (FFA, hypoxia, low glucose) relative to the basal condition, shown as a polar bar plot. **g,** Cell-state proportions at day 0 and at differentiation day 14 across stimulus conditions, using the same cell-state color scheme as in (b).

Integration across villages using reciprocal PCA (RPCA) removed batch effects while preserving biological structure (**Fig. 1b, Extended Data Fig. 1e-j, Methods, Supplementary Notes**). Low-dimensional embedding of the integrated atlas revealed four major cell states: quiescent AMSCs (marked by *PDGFRA*), proliferating AMSCs (*PDGFRA* with cell cycling markers including *POLA1*, *PCNA, RFC3, CCNA2, AURKB*), structural Wnt-regulated adipose tissue-resident (SWAT) cells (*MFAP4, PLAC9, DCN, MGP, APOD*), and adipocytes (*LIPE, PPARG, PLIN1, FABP4*) (**Fig. 1b,d, Extended Data Fig 1k, 2a,** canonical markers were collected from literature ^22,24,25^. Independent from canonical marker gene-based annotations, we applied consensus non-negative matrix factorization (cNMF) to infer active gene expression programs across our dataset **(Extended Data Fig. 2b-d)**. We identified 6 distinct transcriptional programs **(Extended Data Fig. 2c, Supplementary table 3)**, which are highly concordant with the major cell states defined by marker genes, including quiescent AMSCs (cNMF cluster 1), proliferating AMSCs (cluster 6) and adipocytes (cluster 3). Notably, cNMF further resolved three distinct programs within SWAT cells (clusters 2,4,5), revealing a higher level of granularity and heterogeneity within this multipotent cell state **(Extended Data Fig. 2b,c)**. Pathway analysis of the top 100 cNMF cluster-specific genes of the SWAT cell specific programs showed that each program mapped to a distinct functional state. Cluster 2 was enriched for extracellular-matrix organization, elastic-fibre and complement-cascade pathways (e.g. Collagen Fibril Organization, Molecules Associated with Elastic Fibres, Complement and Coagulation Cascades, led by *C1S*/*C3*/*CFD* and *MMP2*/*FBLN5*), suggesting a matrix-remodeling, complement-secreting fibroblast-like state^41,42^. Cluster 4 was enriched for smooth-muscle/actomyosin contraction and TGFβ-SMAD signaling pathways (Vascular Smooth Muscle Contraction, Actomyosin Structure Organization, TGF Smad Signaling, led by *ACTA2*, *TAGLN*, *POSTN*, *CCN2*), indicating a myofibroblast-like profibrotic state^41,43^. Cluster 5 was dominated by cytoplasmic-translation/ribosome and glycolytic pathways (Cytoplasmic Translation, Ribosome, Aerobic Glycolysis, Myc Targets) but lacked cell-cycle genes, identifying a high-biosynthetic, metabolically active state distinct from the cell-cycle-driven proliferating AMSC program (cluster 6)^44^ (**Supplementary table 4**). Within these states, cells did not segregate into discrete sub-clusters but instead formed continuous transcriptional trajectories whose occupancy shifted as a function of metabolic disease stimulation. PCs captured major axes of variation corresponding to adipogenic (PC1) and AMSC/SWAT axis (PC2) branches **(Extended Data Fig. 2e)**. Pseudotime analysis independently recapitulated progressive differentiation along both PC-derived branches, supporting a continuum of cell state trajectories **(Extended Data Fig. 2f-i)**. The mature adipocyte marker gene *ADIPOQ* increased along pseudotime **(Extended Data Fig. 2i)**, while the progenitor marker gene *PDGFRA* declined **(Extended Data Fig. 2g)**. SWAT cell marker gene *APOD* exhibited a transient enrichment at intermediate pseudotime, consistent with their intermediate differentiation status and phenotypic plasticity^24^ **(Extended Data Fig. 2h).** Cell-state labels were then transferred to the snATAC-seq modality, where chromatin accessibility at marker gene loci revealed cell state-specific peak sets recapitulating the same cluster structure (**Figure 1e**).

Differential expressed gene (DEG) and accessible peaks (DAP) analyses between cell states under basal conditions identified 1,452 cell-state specific marker genes and 29,066 cell state-specific chromatin accessible regions (**Supplementary Table 5, 6, Fig1f**). Transcription factor (TF) motif enrichment within cell state-specific peaks highlighted key adipogenic TFs, including CEBPA, CEBPB and PPARG motifs in adipocytes^45^, whereas motifs enriched in quiescent AMSCs included TFs previously implicated to repress adipogenic differentiation^46^ (e.g. TWIST2) (**Extended Data Fig. 1l, Supplementary Table 7**). Pathway enrichment of cell state-specific marker genes further supported canonical functions, including enrichment of PPAR signaling, insulin signaling, fatty acid beta-oxidation, and fat cell differentiation in adipocytes, extracellular matrix assembly, collagen formation in SWAT cells, and cell cycle processes in proliferating AMSCs^24,25,47^ (**Supplementary Table 8**). To further validate the biological relevance of the identified cell states, we compared our atlas with published single-nucleus human adipose tissue and *in vitro* adipocyte differentiation reference datasets^22,24^. We co-embedded our data with these reference datasets using the Symphony reference mapping framework^48^ and performed label transfer to assess correspondence between annotations. Transferred labels demonstrated that cell-state annotations in our dataset robustly recapitulate both *in vivo* adipose tissue cell states **(Extended Data Fig. 2j-l)** and *in vitro* adipocyte differentiation stages **(Extended Data Fig. 2m-p)**. In the joint embedding space, our adipocytes clustered tightly with *in vivo* adipocyte populations, while SWAT cells and AMSCs align closely with the *in vivo* human adipose progenitor cells (hASPCs). Notably, SWAT subsets co-clustered with distinct hASPC subpopulations (**Extended Data Fig. 2k-l**), reflecting transcriptional heterogeneity within this intermediate state. In the *in vitro* space, SWAT cells and adipocytes mapped specifically to the T5 maturation stage^24^ (differentiation day 6) **(Extended Data Fig. 1n, o; Fig. 1a** from ^24^), without mapping to the earlier proliferation or differentiation stages (T1-T4). Consistently, AMSCs mapped to their progenitor T1 stage **(Extended Data Fig. 1n, o)**. These results confirm that the cell states identified in our atlas align with established *in vivo* adipose biology and occupy appropriate positions along the adipogenic differentiation continuum.

To define if human adipocyte villages are responsive to metabolic disease-relevant stimulatory conditions, we defined DEGs (FDR < 0.05; |log2 fold change [FC]| > 0.2) within each cell state comparing stimulated conditions to basal controls. Exposure to FFA, low glucose, or hypoxia induced comparable numbers of DEGs, ranging from 4,074-6,027 in adipocytes and 8,006-13,133 in SWAT cells (**Fig. 1f and Supplementary Table 9**). Similarly, we observed extensive remodeling of the chromatin accessibility landscape in response to stimulation, with the magnitude of these changes varying across cell states and stimuli (**Fig. 1f and Supplementary Table 10**). Adipocytes and SWAT cells exhibited 5,649-39,509 and 4,471-24,038 stimulus-responsive DAPs (FDR < 0.05 and |log2FC| > 0.2), respectively (**Fig. 1f**). Altogether, these results underscore the extensive cell state- and context-specific responses at both the transcriptional and chromatin accessibility levels, highlighting the comprehensive mapping of metabolic disease-relevant gene regulatory signatures enabled by adipocyte cell villages across diverse stimulatory conditions.

We next quantified cell state composition across stimuli **(Fig. 1g)**, showing that SWAT cells constituted a substantial fraction at day 14 across conditions (median=59%-80.2%, IQR=15.8%- 37.9%), whereas the cell state proportions varied markedly upon stimulation. Specifically, adipocytes accounted for 31.4% of cells under basal conditions and were significantly decreased to 14.9% under hypoxia (-16.5%; Wilcoxon test, BH-adjusted *P* = 4.3E-09). In contrast, adipocytes represented 26.7% of cells under low glucose and were significantly increased to 34.3% under elevated FFA exposure (+7.6%; Wilcoxon test, BH-adjusted *P* = 8.8E-04), (**Figure 1g**). These shifts are consistent with the known pro-adipogenic effects of lipid exposure and suppression of terminal adipogenesis under oxygen deprivation^49–51^. Notably, The cNMF-defined adipocyte program (cluster 3) closely matched the canonical marker-based adipocyte annotation across conditions (median donor-level difference <1% in basal, low-glucose, and hypoxia), with a modest excess under FFA (+4.1%), which might be due to early activation of an adipogenic program in lipid-loaded SWAT cells captured by cNMF before switching to full adipocyte identity.

At the donor-level, cell state proportions exhibited substantial inter-individual variability even under basal conditions, where the fraction of adipocytes per donor ranged from 0.9% to 77.3% (mean=32.3%, sd = 19.4%, coeff of var = 60.3%), with comparable variability observed for SWAT cells **(Fig. 1c, Supplementary Notes).** Donor-level cell-state compositions differed across villages (Kruskal-Wallis, BH-adjusted P < 0.01 for all states), with median adipocyte fraction decreasing from village 1 to 4 (49.2, 31.1, 32.7 and 21.5%). However, this inter-individual variability was reproduced within every village (per-village adipocyte ranges 6.6-69.3, 6.3-57.2, 3.6-59.8 and 1-68.2% for villages 1-4), indicating a reproducible biological characteristic rather than a batch effect. Sequencing depth co-varied with village (median UMI 4,018-7,599; Kruskal-Wallis P = 2.5×10⁻¹⁵), but this reflects the lower RNA content of adipocyte nuclei, which showed the lowest sequencing depth and detected genes among all cell states **(Supplementary Notes)**, donors with more adipocytes therefore have lower median depth (Spearman ρ = −0.56). The depth-composition association is thus a consequence of adipocyte composition, not a technical confound of the village differences, which were not associated with nuclei recovery or demultiplexing efficiency **(Supplementary Notes**). Village was included as a covariate in downstream analyses. Donor identity explained the majority of variance in adipocyte fraction (65.9%), far exceeding stimulation (12.4%) and village batch (14.2%) (similar for SWAT cells **Supplementary Notes)**. This donor-dominated variance indicates that inter-individual differences in adipogenic differentiation capacity are largely donor-intrinsic, potentially shaped by underlying genetic variation. Collectively, our comprehensive snMultiome atlas provides strong evidence that it captures core aspects of adipose-related biology and heterogeneity across differentiation under metabolic disease-relevant stimuli.

### Stimuli-responsive genes regulate cellular programs underlying metabolic disease development

To assess whether human adipocyte villages reflect transcriptional responses relevant to metabolic disease, we performed functional enrichment analysis of stimuli-responsive genes (adjusted P < 0.05; |normalized enrichment score [NES]| > 1; **Supplementary Table 11**). We found an overall decrease in processes linked to oxidative metabolism across stimuli, including oxidative phosphorylation, lipid metabolism and thermogenesis in both adipocytes and SWAT cells (**Fig. 2a, Extended Data Fig. 3a and b**). FFA exposure induced DNA damage- and senescence-related processes while repressing oxidative stress detoxification pathways in adipocytes, collectively reflecting the development of a disease-like state (**Fig. 2a, Extended Data Fig. 3a and b**). SWAT cells showed a similar disease-like response to FFAs, which additionally exhibited the up-regulation of genes linked to Maturity-Onset Diabetes of the Young and the repression of the mTOR signaling pathway (**Fig. 2a, Extended Data Fig. 3a**). In contrast, adipocytes exposed to physiological glucose levels exhibited activation of insulin signaling-related processes and repression of genes linked to the senescence-associated secretory phenotype, consistent with the notion that low-glucose stimulation represents a healthy cellular state (**Fig. 2a**). While the insulin signaling pathway was not responsive to low glucose in SWAT cells, glucose metabolism- and adipogenesis-related processes were modulated under this stimulatory condition (**Fig. 2a, Extended Data Fig. 3a and b**). Moreover, comparisons between stimuli FFA, low glucose, hypoxia) (**Supplementary Table 11**) revealed that adipocytes exposed to FFA exhibited a stronger enrichment of metabolic processes (e.g., respiratory electron transport and fatty acid metabolism) compared to hypoxia or low glucose. In SWAT cells, FFA exposure led to positive enrichment of pro-adipogenic programs relative to hypoxia, whereas a negative effect was observed when compared to low glucose.

**Fig. 2.**
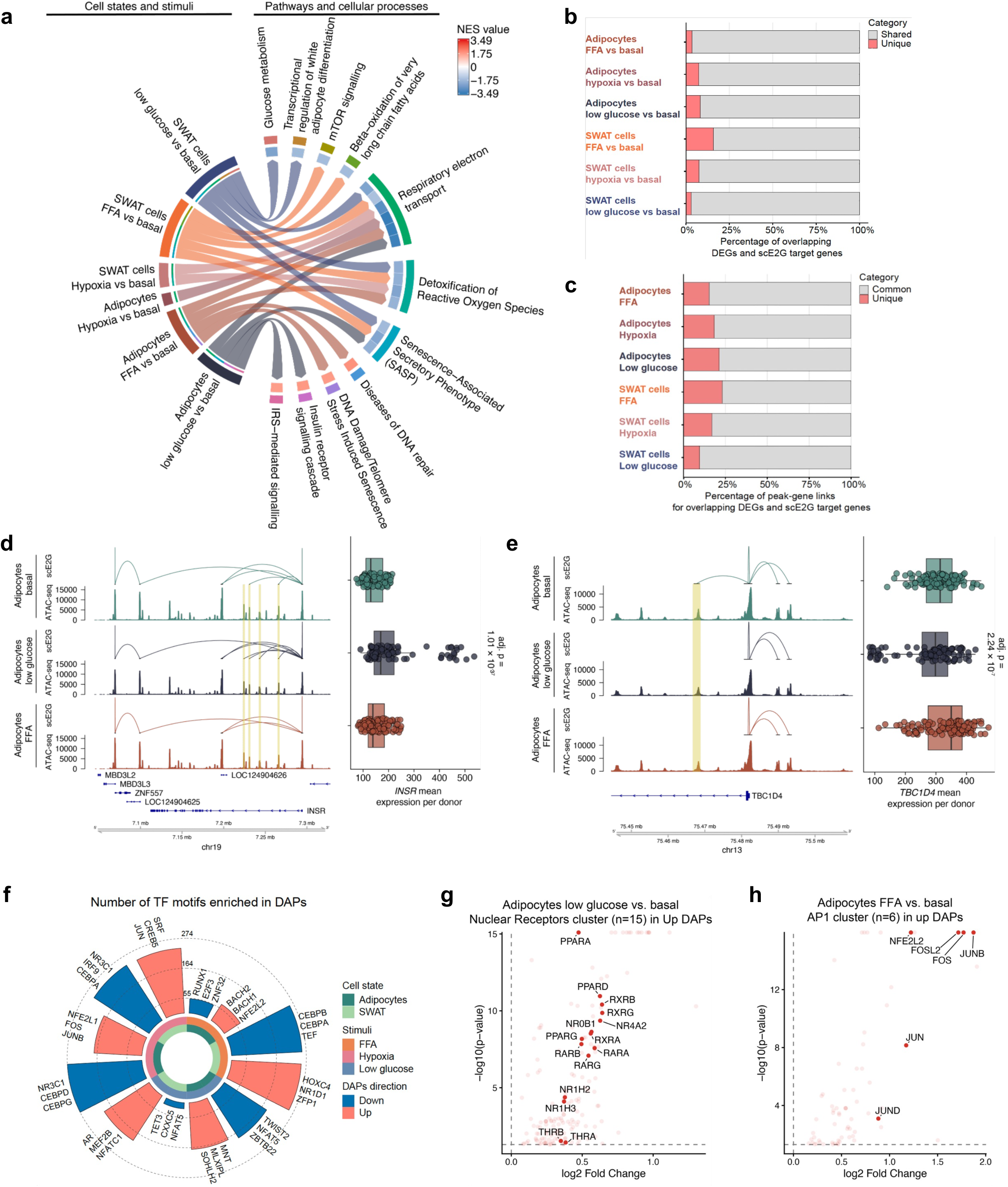
Stimuli regulate gene programs underlying metabolic disease. **a**, Representative metabolic disease-associated cellular programs from functional enrichment analysis (Reactome database; NES: normalized enrichment score) of DEGs after 14 days of exposure to FFA, low glucose (5 mN), or hypoxia compared to the basal state (17.5 mM glucose). **b,** Number of common or unique stimulus-responsive genes (DEGs compared to basal) also predicted as target genes by scE2G analysis across cell state and stimuli. **c,** Number of common or unique peak-gene links for overlapping DEGs and scE2G predicted target genes across cell state and stimuli. **d,e**, Genomic tracks with peak-gene links predictions (scE2G), chromatin accessibility (normalized snATAC-seq signal) and box plots with gene expression (snRNA-seq) at the (**d**) *INSR* and (**e**) *TBC1D4* loci in adipocytes under basal condition and following low glucose or FFA exposure (context-specific peak-gene links are highlighted with a yellow rectangle). **f**, Number of TF motif enrichments in DAPs across cell states and stimuli, with the three most strongly enriched motifs shown. **g,h**, TF motif enrichment in chromatin regions gaining accessibility (Up DAPs) in adipocytes, with (**g**) the nuclear receptor cluster highlighted under low glucose and (**h**) the AP1 cluster highlighter under FFA.

We next leveraged paired snRNA-seq and snATAC-seq data to predict peak-gene regulatory interactions using the single-cell enhancer-to-gene (scE2G) multiome model^52^. We identified an average of 46,878 peak-gene links per cell state and stimulus (**Supplementary Table 12**), with 2,974-4,404 and 4,297-7,945 scE2G target genes also showing significant differential expression following stimulation in adipocytes and SWAT cells, respectively (**Extended Data Fig. 3c**). The majority of these genes were responsive across cell states and stimuli, while context-specific genes represented an average of 6.3% in adipocytes and 8.8% in SWAT cells (**Fig. 2b and Extended Data Fig. 3c**). Among DEGs-scE2G predicted target genes, we identified a total of 16,842-23,477 peak-gene links in adipocytes (18.3% context-specific) and 23,263-41,432 in SWAT cells (16.5% context-specific; **Fig. 2c and Extended Data Fig. 3d**). Notably, consistent with its beneficial effects on insulin signaling, low glucose up-regulated *INSR* (log2 FC=0.5, adjusted P=1.01 × 10^-37^) and down-regulated *TBC1D4* (log2 FC= −0.27, adjusted P= 2.79 × 10^-8^) in adipocytes **(Fig. 2d and e**). The insulin receptor (*INSR*) is a central mediator of insulin signal transduction^53^, and its increased expression was accompanied by the induction of two predicted enhancer-gene interactions following low-glucose exposure in adipocytes (**Fig. 2d, Supplementary Table 12**). In contrast, *TBC1D4* inhibits the translocation of the glucose transporter GLUT4 to the plasma membrane^53^, and its down-regulation under low-glucose stimulation was accompanied by the loss of a predicted enhancer-gene interaction in adipocytes (**Fig. 2e, Supplementary Table 12**).

To further investigate the regulatory basis of these transcriptional changes, we performed TF motif enrichment analysis of DAPs (adjusted P < 0.05, |log2 FC| > 0, and predicted TF mRNA CPM > 1), identifying 123-450 enriched motifs in adipocytes and 218-515 motifs in SWAT cells (**Fig. 2f and Supplementary Table 13**). Among these TF motifs, we found several key regulators of adipocyte differentiation and function^54^ across stimuli, for example as observed for NR3C1, CEBPA and CEBPB in chromatin regions losing accessibility in response to hypoxia in adipocytes and SWAT cells (**Fig. 2f and Supplementary Table 13**). Protein-protein interaction analysis of TFs linked to DAP-enriched motifs further revealed prominent clusters comprising well-established regulators of adipocyte biology (**Supplementary Table 14**). Notably, the healthy state induced by low glucose in adipocytes was associated with enrichment of the nuclear receptor cluster at chromatin regions that gained accessibility (**Fig. 2g**). This cluster comprises several TFs with well-established roles in improving adipocyte metabolic fitness and maintaining glucose homeostasis, including PPARG, PPARD, and NR1H3^54,55^. Interestingly, the nuclear receptor cluster was not observed among upregulated DAPs in SWAT cells exposed to low glucose. Instead, the largest TF cluster corresponded to the SMAD protein complex, which includes TFs previously reported to be expressed in SWAT cells (e.g., SMAD3, TGIF1, and FOXO3)^24^, suggesting a potential role for TGF-β signaling in regulating adaptive responses to low glucose (**Extended Data Fig. 3e**). In contrast, chromatin regions that gained accessibility under the disease-like state induced by FFAs in adipocytes were enriched for the stress-responsive AP1 cluster (**Fig. 2h**). TF members of the AP1 cluster are activated under disease states, where they have been implicated in pro-inflammatory responses and the pathogenesis of insulin resistance^56,57^. In this disease-like context, SWAT cells also exhibited distinct TF clusters, with HOX proteins representing the largest cluster among DAPs gaining accessibility (**Extended Data Fig. 3f**), with these TFs having established roles in adipose tissue development^58^. Altogether, these results demonstrate that adipocyte villages exhibit context-specific transcriptional and chromatin accessibility responses relevant to metabolic disease pathogenesis, characterized by coordinated modulation of gene regulatory mechanisms governing cellular programs associated with both healthy and disease states.

### The enrichment of disease heritability is cell state-, trait- and context-dependent

To assess whether metabolic disease risk preferentially maps to specific cell states and metabolic healthy or disease-like contexts, we applied single cell disease relevance score analysis^59^ (scDRS) using GWAS-derived gene sets for 17 metabolic disease-relevant traits (**Methods**). scDRS quantifies the coordinated expression of disease-associated genes at single-cell resolution relative to matched control gene sets, enabling systematic comparison of heritability enrichment across cell states and stimulation conditions. Disease heritability was not uniformly distributed across cell states, but instead showed distinct cell state-, and trait-specific patterns. Mature adipocytes were strongly enriched for traits related to fat distribution, glucose homeostasis, and lipid metabolism. Such cell state-specificity was further observed by the proportion of significant cells (FDR < 0.2), with adipocytes showing enrichments in 48% for HbA1c, 45.8% for triglycerides, 45.5% for HDL, 42.5% for LDL, 32.% for GFATadjBMI, 21.9% for WHRadjBMI and 9.3% for FI (P < 0.001 for all; **Fig. 3a**, **Extended Data Fig. 4a, and Supplementary Table 15**), which is consistent with their central role in whole body metabolic health. In contrast, SWAT cells and AMSCs showed minimal enrichment for these traits (<3% of cells for most). BMI heritability enrichment was restricted to proliferative and quiescent AMSCs (P = 0.001; 2.8-3.3% of cells with FDR < 0.2), with no significant enrichment in adipocytes (0.1%) (**Fig 3a**, **Extended Data Fig. 4a, and Supplementary Table 15**), consistent with prior evidence linking the strongest genetic risk locus for obesity at the *FTO* locus to preadipocytes rather than mature adipocytes^9^. Complex disease endpoints including T2D, T1D, and CAD showed a negligible enrichment in AMSCs proliferating or adipocytes, with almost no cells reaching significance after multi-testing correction (FDR <0.2) (**Fig 3a, Extended Data Fig. 4a, Supplementary Table 15**), consistent with their heterogeneous pathophysiology involving contributions from multiple tissues and cell types beyond adipocytes^59–64^. Remarkably, a subset of the SWAT cell population showed significant enrichment for HDL heritability (P = 0.002, 9% of cells with FDR < 0.2) **(Fig 3a, Extended Data Fig. 4b)**, suggesting that this progenitor-like population may contribute to lipid homeostasis through mechanisms distinct from mature adipocytes. To explore the molecular basis of HDL heritability enrichment in SWAT cells, we stratified this population by scDRS score and identified differentially expressed genes between high- and low-HDL-enriched subpopulations. High-HDL SWAT cells showed significant enrichment for pathways related to cellular senescence (including telomere maintenance, DNA replication pre-initiation, and mitochondrial ATP synthesis; adjusted P < 0.05) and, notably, negative regulation of canonical Wnt signaling (**Extended Data Fig. 4c, Supplementary Table 16**). Given that SWAT cells are defined by active Wnt signaling and multipotent properties^24,25^, this enrichment may reflect a shift in differentiation potential in response to polygenic HDL-associated regulatory perturbations.

**Fig. 3.**
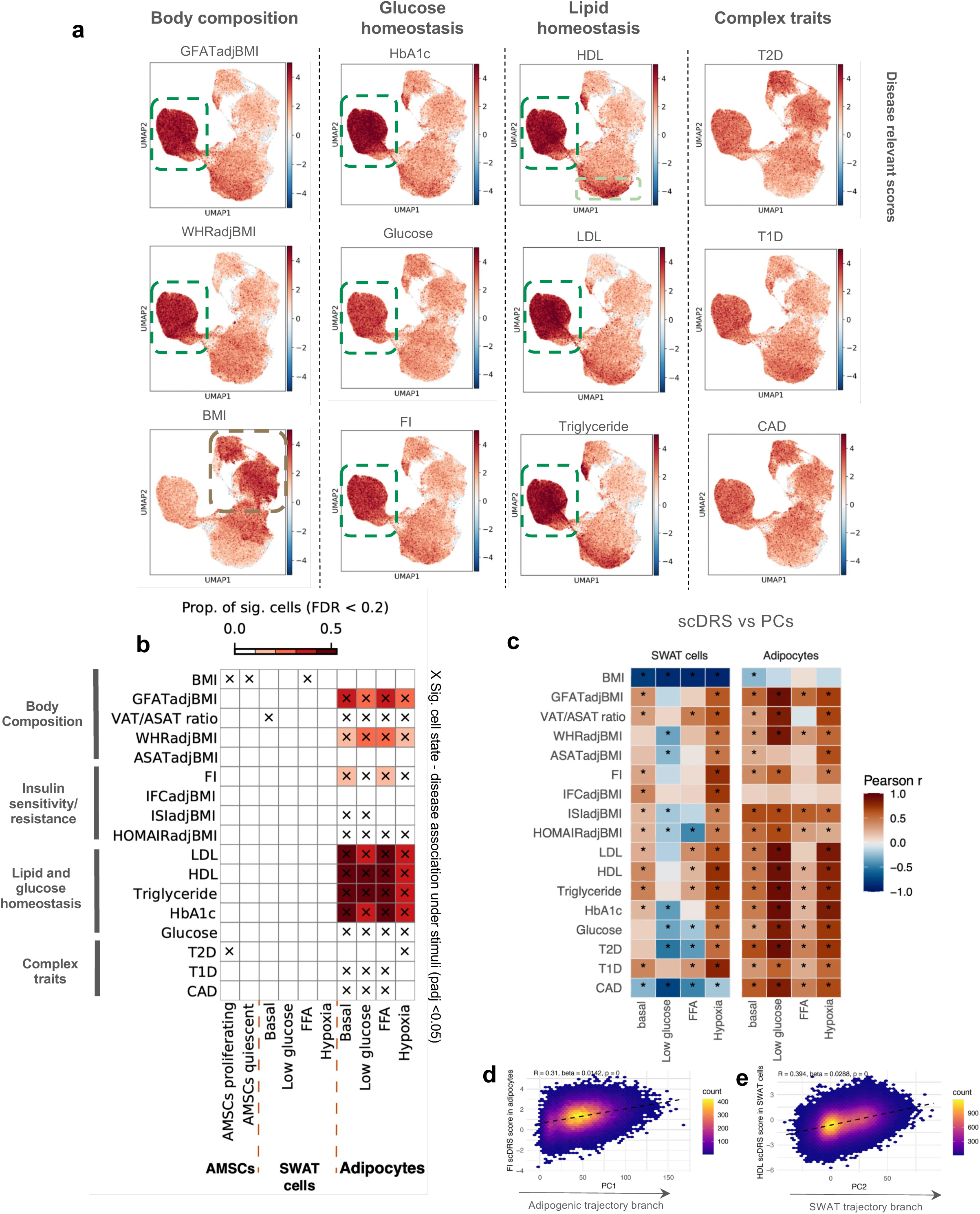
Disease heritability mapping. **a,** UMAP embeddings of metabolic disease heritability mapping across body composition, glucose and lipid homeostasis traits, and complex traits. Colors (blue to red) indicate normalized disease relevant scores of each cell. **b**, Heatmap showing heritability enrichment between cell states under different stimuli (columns) and GWAS traits (rows). Heatmap color represents the proportion of significantly associated nuclei (FDR < 0.2 across all nuclei for a given trait). Cross indicates significant cell-state-disease associations (BH adjusted FDR < 0.05 across all pairs of cell states and traits; MC test). **c,** heatmap showing associations between PC1 (adipogenic trajectory)/PC2 (SWAT trajectory) and single cell disease scores in adipocytes and SWAT cells. **d,e**, Hexbin density plots of single-cell scDRS scores against differentiation-trajectory principal components, with (**d)** FI scDRS score in adipocytes along PC1 (adipogenic trajectory branch), and (**e)** HDL scDRS score in SWAT cells along PC2 (SWAT trajectory branch). Color indicates the number of nuclei per bin, dashed line shows the linear fit.

To evaluate the impact of metabolic contexts on heritability enrichment, we stratified the dataset by both cell state and stimulatory condition (**Fig. 3b, Supplementary Table 17**). Strikingly, nutrient overload exposure under both basal and FFA conditions (both in a high glucose state) consistently induced a greater heritability enrichment than low glucose or hypoxia across multiple trait classes in adipocytes. For lipid traits, basal conditions yielded the strongest enrichment (LDL: 50.5%, triglycerides: 51.7%, HDL: 48.7% of cells with FDR <0.2), representing a 1.3-1.5-fold increase in enriched cells compared to hypoxia (LDL: 35.1%, triglycerides: 36.5%, HDL: 38.2%). Glucose homeostasis traits showed a similar pattern: HbA1c enrichment reached 51-52% of cells under basal and FFA conditions versus 40-45% under low glucose and hypoxia. The effect was most pronounced for fasting insulin, a direct measure of insulin resistance, where FFA exposure yielded 1.5-fold more enriched cells than hypoxia (10.6% vs. 6.9%), and basal conditions showed a 1.4-fold increase (9.7% vs. 6.9%). Fat distribution traits (GFATadjBMI, WHRadjBMI) followed overall comparable trends, with up to 1.4-fold greater enrichment under nutrient overload conditions (basal or FFA vs low glucose or hypoxia). Notably, hypoxia represents a metabolic stress condition relevant to adipose dysfunction, yet nutrient excess, rather than oxygen deprivation, emerged as the context in which genetic risk for cardiometabolic traits is most strongly expressed. On average, nutrient overload conditions captured approximately 9% more heritability-enriched cells than hypoxia across the highly enriched traits examined (mean fold-change: 1.3x), demonstrating that modeling disease-relevant metabolic environments substantially enhances the resolution of genetic risk in adipocytes.

We further applied stratified LD-score regression (s-LDSC) analysis to assess heritability enrichment in cell state-specific pseudobulk snATAC-seq peaks (**Extended Data Fig. 4d**). Due to the limited number of donors available for pseudobulking under the hypoxia condition in adipocytes, this condition was excluded from s-LDSC analysis. After Bonferroni correction, we observed significant enrichment of chromatin accessible regions for traits including FI, GFATadjBMI, ASATadjBMI, and the VAT/ASAT ratio in adipocytes, consistent with the heritability patterns observed in scDRS analyses based on RNA modality. Notably, adipocytes exposed to FFA were not enriched for ASATadjBMI; given that higher ASATadjBMI is associated with favorable metabolic health, this suggests that FFA exposure shifts adipocytes towards a less healthy state. WHRadjBMI heritability showed a distinct pattern: heritability was enriched across all cell states at the chromatin level, but restricted to adipocytes in the RNA modality, indicating broader regulatory priming that may not yet be transcriptionally active in other cell types. Lipid traits (LDL, HDL), despite strong enrichment by scDRS, showed no significant chromatin enrichment, a discordance likely reflecting the limited sensitivity of pseudobulk s-LDSC compared to single-cell approaches. More broadly, while scDRS detected clear stimulation-dependent effects at single-cell resolution, s-LDSC has limited power for partitioning context-specific signals within a given cell state, and chromatin accessibility alone does not reflect the transcriptional output.

We next asked whether disease heritability varies along cell differentiation trajectories by correlating donor-level scDRS scores with principal components capturing differentiation axes: PC1 (adipogenic trajectory) and PC2 (AMSC axis and SWAT trajectory) (**Extended Data Fig. 2e**) under each stimulus. This approach revealed that most fat distribution, lipid, glycemic, and complex disease traits exhibited strong significant positive correlations with the adipogenic differentiation axis across stimulatory conditions, whereas associations with the SWAT axis showed a higher level of context dependency and the effects were comparatively weaker or opposite **(Fig. 3c, Supplementary Table 18)**. This is consistent with cardiometabolic disease-associated transcriptional programs being preferentially expressed in metabolically active, mature adipocytes rather than in progenitor-like populations^22,65^. Interestingly, fat distribution traits as well as lipid and glycemic traits showed stronger correlations with adipocyte maturation under low-glucose exposure compared with other stimuli (**Fig. 3c, Supplementary Table 18**), suggesting that a metabolically favorable environment may sharpen the alignment between cardiometabolic genetic programs and the adipocyte differentiation axis. Notably, BMI showed a strong negative association with the SWAT trajectory across stimuli (r = −0.78 to −0.89, BH-adjusted P < 0.001 for all, **Fig. 3c, Supplementary Table 18**), consistent with previous observations that BMI heritability is enriched in adipocyte precursor states. Importantly, under low-glucose conditions, several fat distribution, glycemic and complex traits (WHRadjBMI, ASATadjBMI, ISIadjBMI, HOMAIRadjBMI, HbA1c and CAD) showed negative correlations with the SWAT differentiation axis, suggesting that a metabolically healthier environment may suppress cardiometabolic disease programs in SWAT cells. Under the disease-like state induced by nutrient overload basal or FFA exposure, lipid traits (LDL, HDL, triglycerides) as well as the VAT/ASAT ratio became positively associated with the SWAT differentiation trajectory **(Fig. 3c, Supplementary Table 18)**, potentially reflecting increased lipid metabolic activity in SWAT cells under lipid-overload condition. In contrast, insulin- and glucose-related traits were negatively associated along the SWAT trajectory **(Fig. 3c, Supplementary Table 18)**, indicating differential sensitivity of metabolic pathways to lipid overload in SWAT versus adipocyte states. To illustrate scDRS and PCs correlation along cell state trajectories at single-cell resolution, we examined two representative traits, FI along the adipogenic trajectory (**Fig. 3d**, r=0.31) and HDL along the SWAT trajectory (**Fig. 3e**, r=0.39), indicating that polygenic risk accumulates progressively across individual cells.

Together, these results indicate that cardiometabolic genetic programs are progressively enriched along the adipocyte differentiation trajectory, serving as the primary repository of genetic risk for fat distribution, glucose homeostasis, and lipid metabolism traits, while SWAT progenitor-like states exhibit greater transcriptional plasticity in response to metabolic stimulations. These analyses demonstrate that cardiometabolic disease heritability is strongly dependent on cell state, metabolic context, and differentiation trajectory.

### Molecular cis-QTL mapping identifies thousands of e/caQTL signals colocalizing with key metabolic disease traits

To investigate how genetic variants influence gene expression and chromatin accessibility across cell states and metabolic disease-relevant stimulatory conditions, we performed genome-wide cis-expression (e) and cis-chromatin accessibility (ca) QTL mapping. Significant QTLs were then tested for colocalization with key metabolic GWAS traits to quantify the overlap with genome-wide significant GWAS loci (PP.H4 >0.7, **Methods; Fig. 4a)**. Pseudobulk QTL discovery stratified by cell states and stimulations using TensorQTL identified 3,374 eGenes of 7,103 eQTLs (eGenes = unique genes that associated with at least one eQTL with q < 0.05) and 25,657 caPeaks of 59,550 caQTLs (caPeaks = unique peaks that associated with at least one caQTL with FDR < 0.05) across cell states and stimuli **(Extended Data Fig. 5a,b, Supplementary Table 19,20)**. To improve detection power and capture shared versus context-specific effects, we applied a multivariate adaptive shrinkage framework^66^ (mash), yielding 6,835 eGenes and 31,941 caPeaks at lfsr < 5% **(Fig. 4b, Extended Data Fig. 5c)**, representing a 2- and 1.2-fold increase over the univariate approach.

**Fig. 4.**
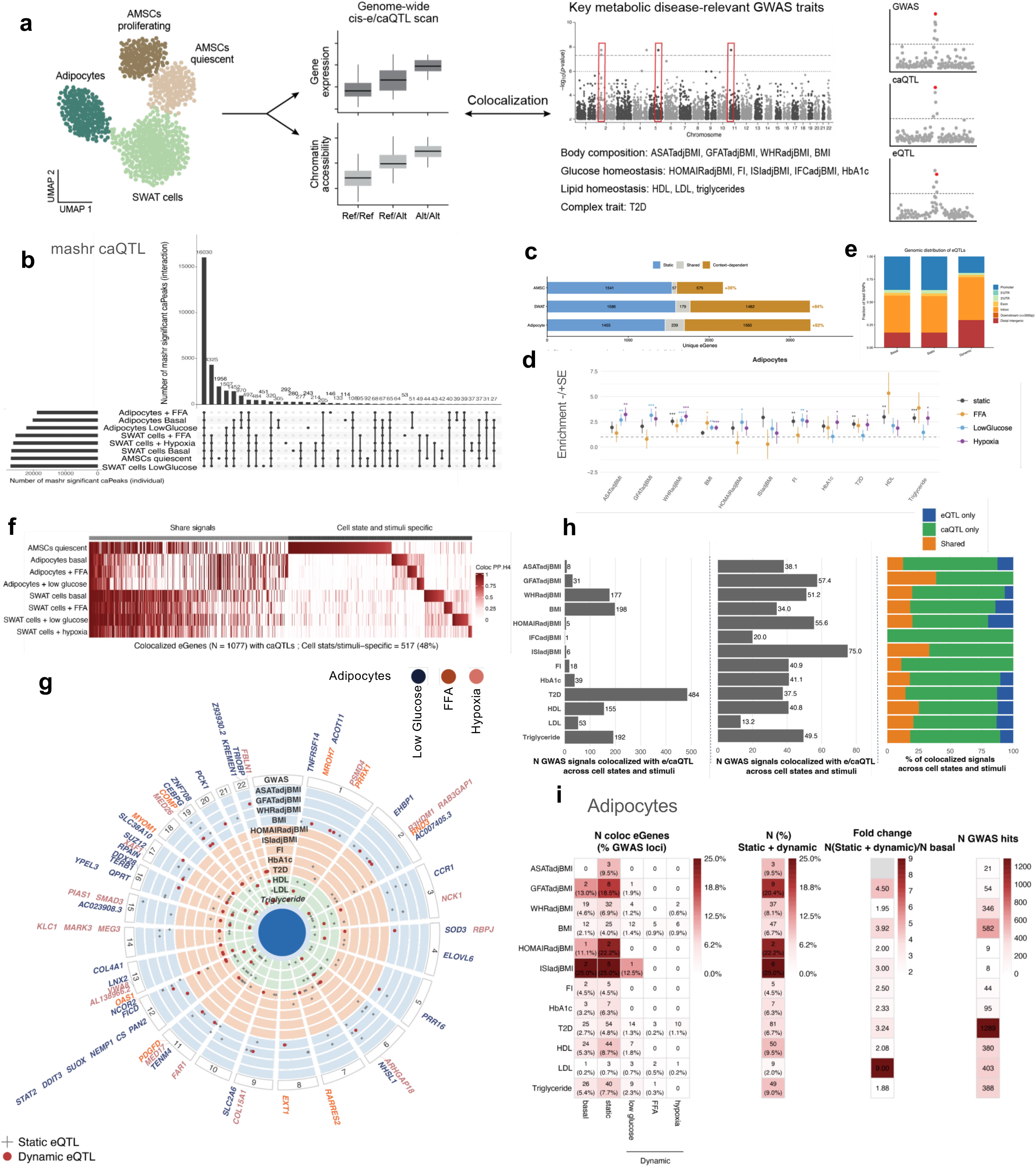
Molecular cis-QTL mapping identifies thousands of e/caQTL signals colocalize with key metabolic disease traits. **a,** Schematic overview of cis-eQTL and cis-caQTL mapping strategy. **b,** UpSet plot showing the number of caPeaks identified across cell states under stimuli by pseudobulk QTL mapping with multivariate adaptive shrinkage (mashr, lfsr <0.05). **c,** Number of unique eGenes in each cell state (AMSCs quiescent, SWAT cells, adipocytes), classified as static, shared or context-dependent. Percentages denote the relative gain in eGene discovery from context-dependent eQTLs (context-dependent as a fraction of static + shared eGenes). **d,** Stratified LD score regression (sLDSC) heritability enrichment (mean ± SE) for adipocyte eQTLs across cardiometabolic GWAS traits, partitioned by context: static, FFA, low glucose and hypoxia. Dashed line marks no enrichment (=1); asterisks denote significance (∗ *P* < *0*.*05*, ∗∗ *P* < *0*.*01*, ∗∗∗ *P* < *0*.*001*). **e,** Genomic distribution of eQTL lead variants from basal condition, static and dynamic effects (promoter, 3’ and 5’ UTR, exon, intron, downstream, distal intergenic). **f,** Heatmap of caQTL-eQTL colocalization, showing eGenes whose expression colocalizes with caPeaks (N of eGenes = 1,077). Columns are ordered into signals shared across cell states/stimuli versus cell-state/stimulus-specific signals; color indicates the posterior probability of a shared variant (PP.H4). **g**, Circular plot of eQTL-GWAS colocalizations in adipocytes across stimuli. Genes are positioned by genomic location and colored by stimulus, concentric tracks correspond to GWAS traits, and points mark significant colocalizations (red dot = dynamic eGenes, grey cross = static eGenes). **h,** Number (left), percentage (middle), and colocalization pattern (right) of GWAS signals that colocalize (PP.H4>0.7) with eQTL and/or caQTL signals across cell states, stimuli and traits. The right panel shows the proportion of colocalized GWAS signals explained by caQTL only, eQTL only or both. **i,** Per-trait eQTL colocalization summary in adipocytes. From left to right: number of colocalized eGenes at basal condition, under static and dynamic effects (and % of GWAS loci explained); number (%) of GWAS signals with static + dynamic eQTL explain; fold change in colocalized signals from static + dynamic versus basal-only models; and number of total GWAS hits. Color intensity scales with each metric.

While pseudobulk eQTLs are informative, they may mask regulatory effects that vary across metabolic contexts. To address this, we applied CASTIE (Context-Aware Single-cell eQTL Tool), a Poisson mixed model framework that jointly models two components at single-cell resolution: (1) main genotype effects (βG), referred to hereafter as “static” effects, representing baseline genetic effects on gene expression at the reference metabolic condition while adjusting for cell states under metabolic contexts and covariates; and (2) genotype-by-context interaction effects (βG×C), referred to hereafter as “dynamic” effects, representing context-dependent changes in genetic effects across cell states and stimuli. Crucially, static effect estimation pools cells across all conditions, providing substantially greater statistical power than any single-condition analysis. This approach identified 5,403 unique eGenes of 8,235 eQTLs, a 1.6-fold increase over pseudobulk eGenes. Critically, 2,758 eGenes and up to 92% eGene per cell states were detected exclusively under specific contexts and not under main genotype effects, demonstrating that a substantial fraction of regulatory variation is masked in unstimulated cells **(Fig. 4c, Extended Data Fig. 5d,e, Supplementary Table 21)**. We found that both static and context-dependent adipocyte eQTLs were significantly enriched for the heritability of cardiometabolic traits using s-LDSC. Context-dependent eQTLs, particularly under low glucose and hypoxic conditions, showed enrichment comparable to or exceeding that of static eQTLs, with the most pronounced context-specific enrichment observed for fat distribution traits (WHRadjBMI, ASATadjBMI, GFATadjBMI) **(Fig. 4d)**. To test whether constitutive and context-dependent regulatory variation occupy distinct genomic locations, we annotated each eQTL’s lead variant relative to gene structure. Static basal cis-eQTLs were strongly promoter-proximal (36.6% of lead variants within ±3 kb of a TSS), as were static effect eQTLs (36.8%). In contrast, dynamic eQTLs were redistributed away from promoters, with only 17.9% promoter-localized, a relative reduction of ∼51% (OR 0.38, P = 3.3×10⁻⁵³) and corresponding enrichment in introns (40.6% to 46.9%; P = 3.3×10⁻⁶) and distal intergenic regions (16.4% to 30.1%; OR 2.19, P = 3.3×10⁻³²). The near-identical promoter fractions of basal and static effect eQTLs indicate this shift reflects the interaction term itself rather than the dynamic modelling framework. Promoter depletion was reproducible across every stimulus in both cell states (37-38% at baseline vs 11-22% under stimulus; all P < 10⁻⁴), and strongest in SWAT cells under fatty-acid exposure (16.8%; OR 0.32, P = 1.0×10⁻³⁶) **(Fig 4e, Supplementary Table 22)**. No existing methods are available for context-specific caQTL mapping at single cell resolution; thus, we retained the pseudobulk strategy for caQTLs in this study.

We next tested if eQTL and caQTL signals were colocalized across cell states and stimuli to nominate gene regulatory mechanisms. We identified colocalized caQTL signals with PP.H4 >0.7 for 1077 eGenes, 48% of these signals (517 eGenes) are cell states and stimuli specific (**Fig. 4f, Supplementary Table 23**). Notably, while we observed colocalized signals shared broadly across cell states, exposure to metabolic disease-relevant stimuli reveal a large fraction of context-specific signals with potential regulatory function that remain undetectable under basal conditions. We then explored whether molQTL signals exhibit colocalization with 13 metabolic disease-related GWAS traits using coloc (PP.H4 > 0.7) **(Methods)**. Using both single-cell context eQTLs and pseudobulk caQTLs, we identified 794 colocalized GWAS-eQTL signals and 4,372 colocalized GWAS-caQTL signals across cell states, stimulatory conditions, and GWAS traits (**Extended Data Fig. 5f,g, Supplementary Table 24-25**). We observed 60 stimuli-specific dynamic eGenes (**Fig 4g, Extended Data Fig. 5f**) colocalized with GWAS traits in adipocytes and 40 in SWAT cells (**Extended Data Fig. 5f, Extended Data Fig. 6a**). Remarkably, several of these dynamic eGenes are well-established regulators of adipocyte metabolism. *PCK1*, encoding the cytosolic phosphoenolpyruvate carboxykinase that drives adipocyte glyceroneogenesis and the re-esterification of free fatty acids into triglycerides^67,68^, harbored a low-glucose–responsive eQTL that colocalized with triglyceride, HDL cholesterol and waist-to-hip-ratio signals (PP.H4 = 0.89, 0.78 and 0.75, respectively). *RARRES2*, encoding the adipokine chemerin, a key regulator of adipogenesis and insulin sensitivity^69,70^, colocalized with type 2 diabetes specifically under free-fatty-acid stimulation (PP.H4 = 0.9). And *SMAD3*, a TGF-β signalling effector whose loss protects mice from diet-induced obesity and insulin resistance^71,72^, colocalized with type 2 diabetes specifically under hypoxia (PP.H4 = 0.91).

Overall, 13.2-75% of GWAS loci were explained by at least one molQTL **(Fig 4h)**. The majority of colocalizations were driven by caQTLs, particularly for T2D and BMI, which is consistent with previous studies showing that caQTLs capture regulatory variation not detected at the expression level^73^. Notably, the cell states in which we detected the most QTLs were not the most disease-relevant. Quiescent AMSCs and SWAT cells also yielded a large number of eGenes (**Fig. 4c**, **Extended Data Fig. 5d**), likely reflecting greater statistical power due to cell abundance and transcriptional homogeneity. However, GWAS colocalization was concentrated in adipocytes rather than AMSCs and SWAT cells (**Extended Data Fig. 6b-d**), indicating that regulatory variation in progenitor cells, while readily detectable, is less relevant to cardiometabolic disease mechanisms. This pattern is consistent with our findings from heritability enrichment analysis, where disease risk was mostly enriched in mature adipocytes rather than AMSCs, underscoring the importance of the importance of mapping QTLs in disease-relevant cell states and contexts for connecting genetic variation to pathophysiological mechanisms.

Colocalization rates aligned well with known contributions of adipose tissue to metabolic disease-related traits. Fat distribution traits, whose genetic architecture is closely linked to adipose tissue function^74^, showed strong colocalization (ASATadjBMI: 36.1%, GFATadjBMI: 57.4%, WHRadjBMI: 51.2%) (**Fig. 4h**). Fasting insulin and HbA1c, both of which adipose dysfunction is a major contributor to systemic dysregulation, also showed substantial colocalization (FI: 40.9%, HbA1c: 41.1%) (**Fig. 4h**). HDL and triglycerides, which are regulated across liver, adipose, and intestine, also showed high colocalization (HDL: 40.8%, triglycerides: 49.5%), consistent with a meaningful adipose functional contribution. In contrast, LDL showed the lowest colocalization rate (13.2%), consistent with its main regulation in the liver^75^ (**Fig. 4h**). For traits related to insulin sensitivity (FI, HOMAIRadjBMI, IFCadjBMI, ISIadjBMI), which have relatively few genome-wide significant GWAS loci, we observed up to 75% colocalization across cell states and stimuli (**Fig. 4h**). Notably, these signals were captured primarily through caQTLs, and no colocalized signals were detected for IFCadjBMI in adipocytes (**Extended Data Fig. 6b-e**). Despite T2D heritability not being enriched in adipocytes, 37.5% of T2D GWAS signals are colocalized with at least one molQTL (**Fig. 4h**), likely reflecting the contribution of adipose tissue dysfunction-driven insulin resistance to T2D pathophysiology^18^.

A key question is whether metabolic context influences the discovery of disease-relevant regulatory variants. Stimulation context-aware modeling yielded substantially greater colocalization discovery than conventional basal eQTL analysis (**Fig. 4i, Extended Data Fig. 6f, Supplementary Table 24**). In adipocytes, context-aware modeling identified a total of 2.6-fold more GWAS colocalizations than basal-only eQTLs (305 vs 117) across traits. The magnitude of improvement was trait-dependent: LDL (9-fold) showed the largest gain, followed by ASATadjBMI and GFATadjBMI (3 hits vs 0 hits, 4.5-fold), BMI (3.9-fold), T2D (3.2-fold), ISIadjBMI (3-fold), fasting insulin (2.5-fold), and HbA1c (2.3-fold) (**Fig. 4i)**. This improvement derives from two complementary factors. First, static effect joint estimation of baseline genetic effects provides greater statistical power than single-condition analysis, yielding 2.5-fold more adipocyte eGenes (1,694 vs 676 from basal-only single-cell QTL mapping). Static effect colocalizations in adipocytes accounted for the majority of signals (65%, 341 of 523 across traits). Second, genotype-by-context interactions capture an additional 35% of colocalizations (182 of 523) distributed across FFA (26 colocalizations), low glucose (102), and hypoxia (54), representing regulatory variants entirely undetectable without metabolic stimuli **(Supplementary Table 24)**. This proportion was highest for ISIadjBMI (25%) and GFATadjBMI (20.4%) in differentiated adipocytes. Across cell states and stimuli, FFA, a nutrient overload condition modeling the disease state, contributed the most stimuli-specific colocalizations overall, predominantly for traits associated with metabolic disease risk including HbA1c and WHRadjBMI in SWAT cells. In contrast, in adipocytes, low glucose was the dominant context, contributing 56% (102 of 182) of stimuli-specific signals. Of note, the high glucose concentration in our basal medium (17.5 mM) is widely used *in vitro* to recapitulate features of hyperglycemic stress, whereas low glucose (5 mM) more closely approximates normoglycemic conditions. Strikingly, this normoglycemic condition was the only one that captured colocalizations with protective cardiometabolic traits in adipocytes, including the favorable gluteofemoral fat mass trait GFATadjBMI^76^ and the proxy for insulin sensitivity ISIadjBMI^77^. This pattern was not observed in SWAT cells. These findings suggest that metabolic context not only increases discovery power but determines which biology is revealed, where disease states expose pathological mechanisms, while the healthy metabolic state uniquely unmasks protective ones.

Collectively, these results demonstrate that mapping molQTLs in adipocytes across metabolically distinct conditions representing both health and disease states substantially improves the resolution of GWAS mechanisms. The 2.6-fold increase in colocalized signals in adipocytes achieved through context-aware single-cell modeling, combined with the 35% of signals detected exclusively through genotype-by-context interactions, underscores that disease-relevant stimuli are essential for revealing the full regulatory architecture of cardiometabolic diseases.

### Distinct genetic regulatory patterns across cell states and metabolic contexts

To better understand how genetic variation influences adipocyte regulatory programs across metabolic stimuli, we classified GWAS-colocalized molQTL signals based on their regulatory architecture. We observed three distinct patterns (**Fig. 4h**): 1) GWAS loci colocalized exclusively with eQTLs, suggesting gene-proximal regulatory mechanisms; 2) GWAS loci colocalized only with caQTLs, indicating that genetic effects may act primarily through chromatin accessibility changes that do not necessarily propagate to gene expression under specific contexts; and 3) loci showing concordant colocalization with both eQTLs and caQTLs, supporting a regulatory cascade in which genetic variants influence chromatin accessibility and downstream gene expression. These distinct patterns illustrate multiple molecular routes through which genetic variation can influence adipocyte biology and cardiometabolic disease risk.

The *NCOR2* locus exemplifies this principle. The NCOR2/HDAC3 co-repressor complex plays a critical role in inhibiting adipocyte differentiation and regulating lipid metabolism, and its disruption affects lipolysis and insulin sensitivity^78,79^. We identified a low glucose-specific eQTL (rs7311233-*NCOR2*, P_eQTL_ = 8.97E-07) in adipocytes **(Fig. 5a,b,c,e,f, Supplementary Table S21)** that colocalizes with disease protective traits ISIadjBMI (GWAS lead variant rs1906937 at the *RFLNA* (also known as *FAM101A*) locus, PP.H4 = 0.71) (**Fig. 5b)** as well as with disease risk trait T2D (**Fig. 5c**, PP.H4 = 0.72-0.73, GWAS lead variants rs11057418, rs34680764, rs2229840). Notably, other eGenes (*CCDC92*, *ZNF664*, *RFLNA*) near this locus colocalize with the same and other GWAS hits (PP.H4 = 0.88-0.99 for colocalization with ISIadjBMI, GFATadjBMI, HDL, FI, T2D and triglycerides) through static eQTL effect rather than low-glucose-specific effects, highlighting that NCOR2 represents a context-specific regulatory contribution at this multi-gene locus (**Supplementary Table S24, Extended Data Fig. 7a-f**). Critically, the NCOR2 eQTL signal was detected only under low glucose, the normoglycemic condition, and was absent under disease-associated conditions (high glucose, FFA, hypoxia). Low glucose-exposed adipocytes also showed the strongest total cell-level allelic effect^80^ (**Methods**) compared to other conditions (total cell-level allelic effect under low glucose=0.187, P_eQTL_=8.97E-07; FFA=0.034, P=0.52; hypoxia=0.067, P=0.33; basal=-0.038, P=0.14).

**Fig. 5.**
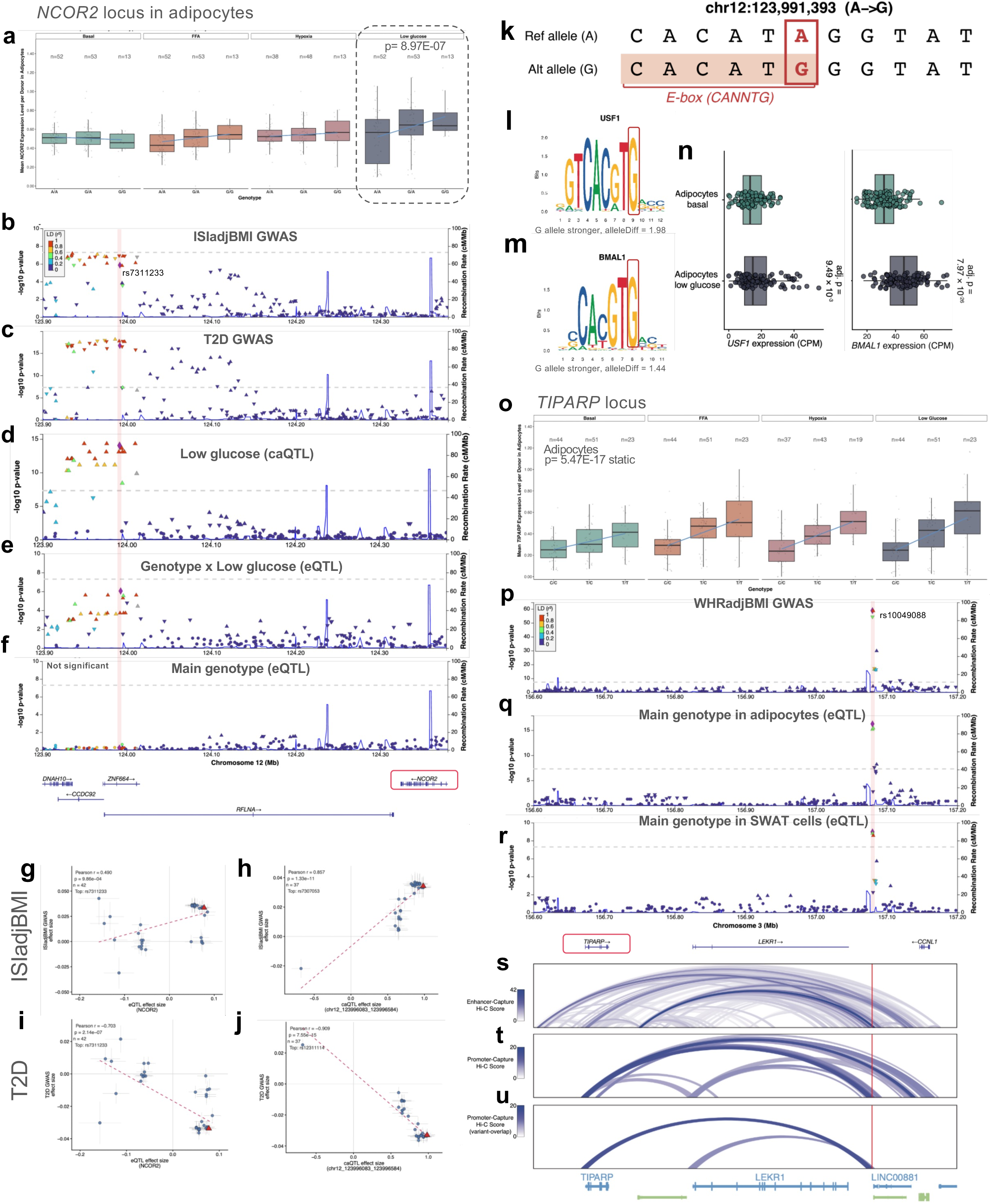
Cell-state- and context-specific and shared e/caQTL examples at the *NCOR2* and *TIPARP* loci. **a,** *NCOR2* expression by genotype across stimuli in adipocytes, showing a low-glucose-specific eQTL (dashed box). **b,c,** Locus plots showing the colocalization of the *NCOR2* eQTL in adipocytes under low-glucose exposure with GWAS signals for **(b)** ISIadjBMI and **(c)** T2D. Each panel shows −log₁₀(P) for the GWAS (left y-axis) and the recombination rate (right y-axis); points are colored by LD (r²) with the eQTL lead variant rs7311233 (vertical line with purple diamonds). **d-f,** Colocalization at the same locus for **(d)** the low-glucose caQTL, **(e)** the genotype × low-glucose (dynamic) eQTL, and **(f)** the main genotype effect (static) eQTL, which is not significant. **g-j**, Scatter plots of variant-level posterior associations versus GWAS for the *NCOR2* eQTL **(g,i)** and caQTL **(h,j)**, against ISIadjBMI **(g,h)** and T2D **(i,k)**. **k,** Reference (A) and alternate (G) allele sequences at chr12:123,991,393. The alt allele (G) creates a CANNTG-family E-box (boxed); the variant nucleotide is highlighted. **l,m,** Position weight matrix logos for **(o)** USF1 and **(p)** BMAL1 (consensus E-box sequence CACGTG). The G allele increases predicted binding (alleleDiff for USF1 and BMAL1 are indicated below the logos). **n,** *USF1* and *BMAL1* expression in adipocytes under basal versus low-glucose conditions, showing higher expression under low glucose (adj. *P* as indicated). **o,** *TIPARP* expression by genotype across cell states and stimuli, showing a static eQTL in adipocytes. **p-r,** Colocalization of the *TIPARP* locus with **(p)** WHRadjBMI GWAS and the main genotype effect (static) *TIPARP* eQTL in **(q)** adipocytes and **(r)** SWAT cells (lead eQTL variant rs10049088). **s-u,** 3D chromatin contact maps at the 3q25 (*TIPARP*) locus, showing distal cis-regulatory loops overlapping the lead WHRadjBMI risk variant (red line). **s,** Enhancer-capture Hi-C in cultured human AMSCs^86^. **t,u,** Promoter-capture Hi-C in human white adipose-derived adipocytes^87^, showing **(t)** all detected promoter-anchored contacts and **(u)** those restricted to contacts overlapping the WHRadjBMI lead variant.

The caQTL signal at the *NCOR2* locus (chr12:123,996,083-123,996,584) was robustly detected in adipocytes across three conditions (basal, low glucose and FFA), tagged by two tightly linked variant pairs in strong LD within each pair (rs4765219/rs11057409 and rs2178663/rs12311114, ∼6 kb apart) (**Supplementary Table S20;** under low glucose: P_caQTL_ = 4.91E-15, FFA: P_caQTL_ = 2.25E-10, Basal: P_caQTL_ = 1.99E-11). All four SNPs showed consistent effect direction and magnitude across stimuli (beta range 0.79-0.98, p < 1 × 10⁻⁹ in every condition), indicating that chromatin accessibility at this regulatory element is broadly maintained across stimuli. These caQTLs are also colocalized with the same GWAS signals of ISIadjBMI, GFATadjBMI, HDL, FI, T2D and Triglycerides) as the eQTLs (PP.H4 = 0.72-0.99, **Fig. 5d, Supplementary Table S25**) across stimuli. *NCOR2* expression and chromatin accessibility at the same locus are positively associated with the metabolically protective trait ISIadjBMI and negatively associated with disease trait T2D (**Fig. 5g-j**). Unlike the eQTL, the caQTL signal at the *NCOR2* locus showed no specificity to low glucose exposure (**Supplementary Table S25**), we asked whether a mechanism at the DNA sequence level could explain the low glucose-specific eQTL effect. TF motif binding analysis at this locus revealed that the alt allele (G) creates a consensus E-box sequence (CACATG) of the CANNTG family^81,82^ that is recognized by bHLH family transcription factors (**Supplementary Table 26, Fig. 5k, Extended Data Fig. 7g**), including the glucose-responsive USF1/USF2^83,84^ and circadian CLOCK/BMAL1^85^ (**Fig. 5l, m)**. Notably, both *USF1* and *BMAL1* (encoded by *ARNTL*) showed significantly higher expression under low glucose compared to basal conditions (**Fig. 5n**). Together, these results support a model in which the alt allele creates a permissive E-box motif whose activity requires context-dependent activation of these bHLH transcription factors, thereby driving the low glucose-specific increase in *NCOR2* expression. Collectively, this locus would not have been identified without modeling metabolic context, highlighting how disease-relevant stimuli can unmask genetic effects on insulin sensitivity.

In contrast to context-specific effects, some regulatory variants show consistent effects across metabolic conditions, providing validation of core adipocyte biology. The *TIPARP* locus, where *TIPARP* regulates maturation of both white and brown adipocytes^82–84^, harbors eQTLs (rs10049090-*TIPARP* and rs10049088-*TIPARP*) with strong static effects in adipocytes and SWAT cells (**Fig. 5o, Extended Data Fig. 7h, Supplementary Table S21**). These eQTLs colocalize with GWAS signals for WHRadjBMI, HDL, T2D, triglyceride across both cell states **(Supplementary Table S24)**. For example, the WHRadjBMI GWAS locus (lead SNP rs10049088) colocalized with *TIPARP* eQTLs (PP.H4 >0.99 for both adipocytes and SWAT cells) (**Fig. 5p-r, Supplementary Table S24**), and colocalized with caQTLs across stimuli in adipocytes (basal PP.H4 = 0.91-0.99; FFA PP.H4 = 0.75, low glucose PP.H4 = 0.93) (**Supplementary Table S25)**. 3D chromatin maps derived from cultured human AMSCs^86^ (Enhancer-capture Hi-C, **Fig. 5s**) and human, white adipose-derived adipocytes^87^ (Promoter-capture Hi-C, **Fig. 5t, u**) reveals many distal cis-regulatory loops overlapping the lead WHRadjBMI risk variant in the 3q25 risk locus. These data nominate several effector-target genes, including *TIPARP* **(Fig. 5s-u)**. Together, molQTL and capture Hi-C approaches results nominate and provide physical interaction evidence for a distal, cis-regulatory mechanism of gene-regulation for *TIPARP*. The consistency of this signal across molecular layers, cell states, and metabolic contexts suggests it represents a fundamental regulatory mechanism in adipose biology.

Finally, some GWAS signals colocalize exclusively with eQTLs without corresponding caQTLs. The *PTGER3* locus associated with HDL levels colocalizes with an eQTL regulating *PTGER3* expression (rs500647-*PTGER3*, this eQTL lies in an intron of *PTGER3* and ∼60kb away from the TSS) in adipocytes with static effects (PP.H4 = 0.997, **Extended Data Fig. 7i-k; Supplementary Tables S21, S24),** without a corresponding colocalized caQTL, suggesting gene-proximal regulation. *PTGER3* encodes a prostaglandin E receptor implicated in the regulation of lipolysis and adipose lipid metabolism^88,89^, and was more recently identified as a marker of a lipid-deposition adipocyte subpopulation^90^.

Together, these examples highlight the distinct genetic regulatory mechanisms operating across cell states and metabolic contexts, and demonstrate that integrating chromatin accessibility, gene expression, and metabolic contexts is essential to resolve the causal paths from variant to cardiometabolic trait.

### Cell state- and context-specific and shared pleiotropic associations pinpoint causal regulations of metabolic diseases

While colocalization indicates whether molQTLs and GWAS signals share a common causal variant, it does not estimate the direction or magnitude of the underlying effect. To quantify the putative pleiotropic links between genetic regulation and metabolic traits, we applied summary-data-based Mendelian randomization (SMR), using our cis-molecular QTLs as instruments to identify variant-eGene/caPeak-trait associations. We further used caQTL as instruments to infer causal relationships between chromatin accessibility and gene expression, allowing us to reconstruct three-way regulatory chains linking accessible chromatin, gene expression, and GWAS traits (caQTL→eQTL→GWAS) (**Fig. 6a**).

**Fig. 6.**
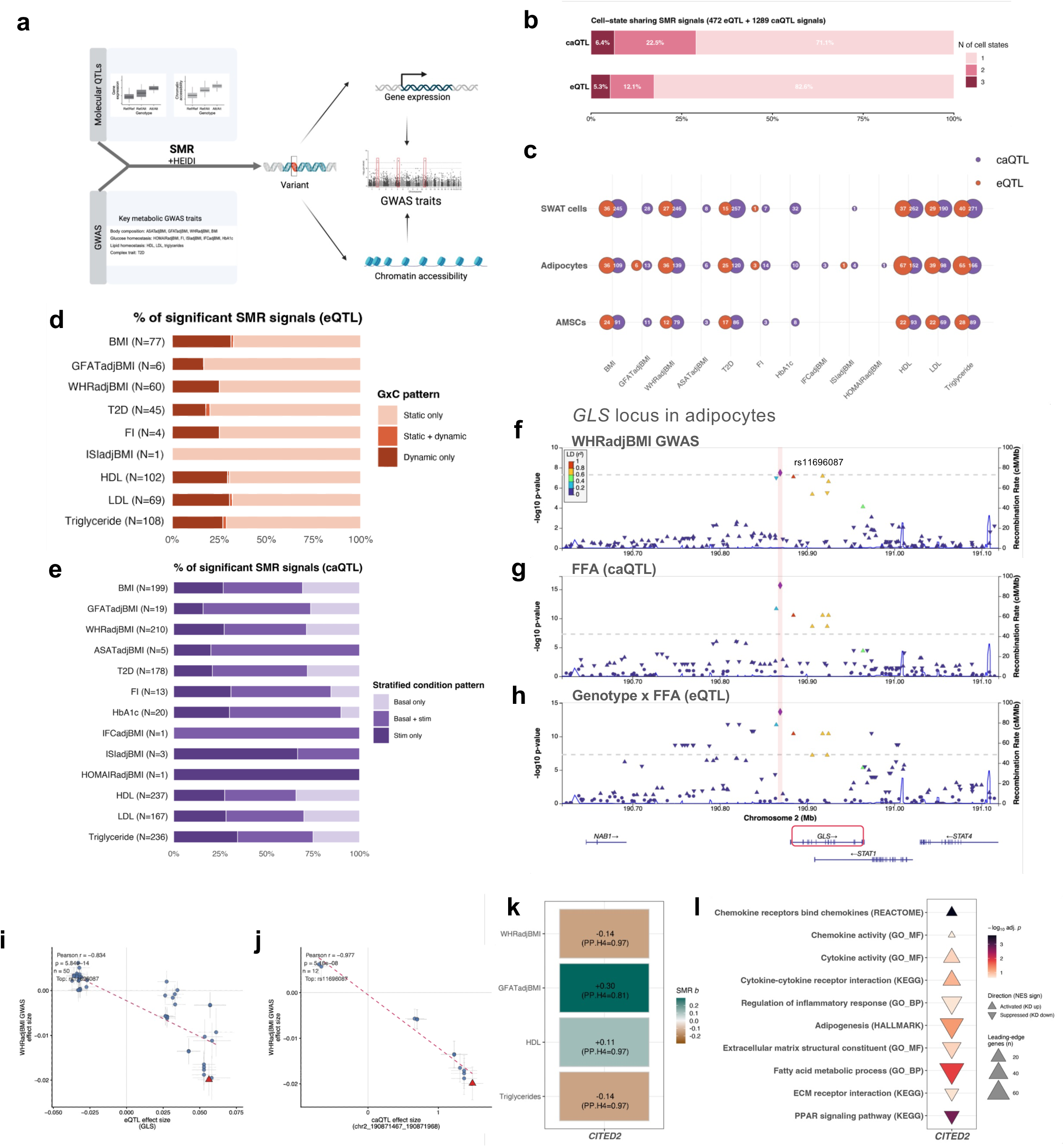
Summary-data-based Mendelian randomization links molecular QTLs to cardiometabolic traits through directional regulatory cascades. **a,** Schematic of the SMR + HEIDI framework. cis-eQTLs and cis-caQTLs are used as instruments to test for pleiotropic associations between molecular phenotypes (gene expression, chromatin accessibility) and cardiometabolic GWAS traits. cis-caQTLs are also used as instruments to test for associations between chromatin accessibility and gene expression. **b,** Cell-state sharing of significant SMR pairs. Bars show the proportion of caQTL (top) and eQTL (bottom) associations that are significant in one, two, or three cell states. **c,** Number of significant SMR pairs per cell state (SWAT, adipocyte, AMSC) and GWAS trait, for eQTL (orange) and caQTL (purple). Dot size scales with the number of pairs, numbers are indicated within each dot. **d,** Distribution of significant eQTL SMR signals by GxE pattern for each trait: static only, static + dynamic, or dynamic only. N indicates the number of significant eGene-trait associations per trait. **e,** Distribution of significant caQTL SMR signals by stratified condition pattern for each trait: basal only, basal + stimulated, or stimulated only. N indicates the number of significant caPeak-trait associations per trait. **f-h,** Locus plots showing colocalization of **(f)** the WHRadjBMI GWAS signal, **(g)** the caQTL under FFA exposure, and **(h)** the genotype × FFA (dynamic) eQTL at the *GLS* locus in adipocytes. Each panel shows −log₁₀(P) (left y-axis) and recombination rate (right y-axis); points are colored by LD (r²) with the lead variant rs11696087 (highlighted). **i,j,** Scatter plots of variant-level effect sizes for WHRadjBMI GWAS versus (i) *GLS* eQTL and **(j)** caQTL (chr2:190,871,467-190,871,968) effect sizes. Pearson r and P as indicated; lead variant rs11696087 (red triangle). **k.** SMR effect estimates (*b*_SMR) for adipocyte *CITED2* expression against four cardiometabolic traits. All associations passed the HEIDI test (*P*_HEIDI > 0.01) and colocalized with the corresponding GWAS signal (PP.H4 = 0.81-0.97); full statistics in Supplementary Table S28. **l.** Gene-set enrichment (GSEA) of the transcriptional response to *CITED2* knockdown by CRISPRi in differentiating human adipocytes, relative to non-targeting controls. Triangle orientation denotes NES sign (up, activated on KD; down, suppressed on KD); fill color, −log₁₀ adjusted *p*; size, number of leading-edge genes (GSEA FDR < 0.25).

We identified 472 eGene-trait and 1,289 caPeak-trait pleiotropic associations across the three cell states and 13 cardiometabolic traits (significance: *P*_SMR FDR < 0.05, HEIDI *P* > 0.01 except for dynamic eQTLs, cis-QTL instrument threshold *P* < 1E-05) (**Fig. 6b,c**). Most associations were specific to a single cell state for both eQTL (82.6%) and caQTL (71.1%) signals, with only 5.3% of eGene-trait and 6.4% of caPeak-trait associations shared across all three cell states (**Fig. 6b**). The predominance of cell state-specific signals indicates that the genetic regulation underlying metabolic traits is largely deployed in a cell state-specific manner, with chromatin-level associations slightly more often shared across cell states than gene expression-level associations. More caPeak-trait pleiotropic associations were observed than eGene-trait associations across nearly all cell states, stimuli, and traits (**Fig. 6c, Extended Data Fig. 8a,b**), consistent with the colocalization results, in which caQTLs colocalized with and explained more GWAS signals than eQTLs. Approximately 32-97% of SMR significant signals were independently supported by colocalization (eQTL-GWAS: 189/588, 32.1%; caQTL-GWAS: 1,154/2,914, 39.6%; caQTL-eQTL: 339/348, 97.4%; PP4 > 0.7), indicating substantial concordance between the two complementary methods (**Supplementary Table S27, 28, 29**).

We next examined how metabolic context shaped these associations by stratifying eQTL SMR signals according to their static (G) and dynamic (G × C) regulatory patterns, and caQTL signals according to the cell states and stimulations. Most eGene-trait associations were driven by static eQTLs (static only); a smaller fraction were attributable to context-interaction effects, comprising signals supported by both static and dynamic models (static + dynamic) and signals detected through dynamic effects alone (dynamic only) (**Fig. 6d**). The contribution of dynamic-only signals was most pronounced for FI, Triglyceride, BMI, and HDL, indicating that for these traits a meaningful proportion of expression-level genetic regulation is revealed only when metabolic context is modeled. CaPeak-trait associations were predominantly detected under stimulated conditions, either in stimulated states alone (stim only) or in both basal and stimulated states (basal + stim), rather than in the basal state only (**Fig. 6e**), indicating that much of the chromatin-level genetic regulation relevant to metabolic traits is unmasked by metabolic stimulation.

To test directional cascades linking chromatin, gene expression and trait, we identified 31 high-confidence three-way chains in which all three associations (caPeak→eGene, caPeak→trait, and eGene→trait) were SMR-significant and directionally concordant. These chains reflect a genetic variant that plausibly alters chromatin accessibility, which in turn alters gene expression and ultimately the metabolic trait. Of these, 8 were further supported by both e/ca QTL-trait colocalization (**Supplemental Table S30**). At the 12q24.31 locus, a well-replicated insulin-resistance locus^91^ that also harbors *NCOR2*, rs7975482 anchors a directionally concordant chain in which accessibility at chr12:123996083-123996584 raises *CCDC92* expression and lowers FI risk (*P*_SMR = 6E-06, *b*_SMR = −0.05, HEDI *P* = 0.97, **Supplemental Table S30**), with support on both e/caQTL-FI colocalization (PP.H4 = 0.99 each, **Supplemental Table S24,25**) under static effect in adipocytes. *CCDC92* influences adipocyte differentiation and lipid storage in human cellular models, and the insulin-raising alleles at this locus lower its expression in subcutaneous adipocytes^91^, consistent with the direction we identified. Notably, the *NCOR2* low glucose specific signal described above has robust colocalization with multiple metabolic traits, reaching nominal SMR significance but not the FDR-corrected cutoff (eQTL-ISIadjBMI *P*_SMR = 5.5E-04, FDR >0.05). This illustrates that context-specific regulatory effects, which are restricted to a single metabolic context and therefore supported by fewer instruments, can be penalized by stringent multiple-testing correction, underscoring the complementary value of colocalization for recovering context-dependent variant-gene-trait links that SMR may miss.

An example of cell state-specific and dynamic cascade regulation was observed at rs11696087. At this locus, the caPeak chr2:190871467-190871968 showed pleiotropic association with WHRadjBMI (caQTL-WHRadjBMI *P*_SMR = 1.37E-06, HEIDI *P*= 0.1) mediated through its effect on *GLS* expression specifically in adipocytes under FFA exposure (eQTL-WHRadjBMI *P*_SMR =7.45E-06; caQTL-eQTL *P*_SMR =2.73E-06, HEIDI *P*=0.4, **Fig 6f-h, Supplemental Table S30**). GLS encodes the rate-limiting enzyme of glutaminolysis, a pathway central to adipocyte amino-acid and TCA-cycle metabolism^92^. The G allele at rs11696087 increases chr2:190871467-190871968 accessibility and GLS expression and is associated with lower WHRadjBMI, suggesting that enhanced glutaminolytic capacity may favour peripheral fat partitioning during fatty-acid stress (**Fig 6i, j, Extended Data Fig.8c**). This three-way regulatory cascade links adipocyte glutamine metabolism to genetically determined body-fat distribution, underscoring how context-specific exposures can unmask genetic variation.

Of particular relevance, SMR-based analysis uncovered that the variant rs9800462 driving higher *CITED2* expression was associated with an overall protective profile. This was characterized by greater GFATadjBMI (*b*_SMR = 0.30, *P*_SMR = 1.0 × 10⁻⁵) and HDL cholesterol (*b*_SMR = 0.11, *P*_SMR = 1.2 × 10⁻⁷), whereas exhibiting lower triglycerides (*b*_SMR = −0.14, *P*_SMR = 5.0 × 10⁻⁸) and WHRadjBMI (*b*_SMR = −0.14, *P*_SMR = 2.1 × 10⁻⁷; all aforementioned traits with HEIDI *P* > 0.01, **Fig. 6k, Supplementary Table S28**). Importantly, each association colocalized with the corresponding GWAS signal (PP.H4 = 0.81-0.97, Supplementary Table S24), altogether implicating genetic mechanisms regulating adipocyte *CITED2* expression as a potential driver of a metabolically favorable phenotype. Next, we experimentally examined the transcriptome-wide effect of *CITED2* suppression by CRISPR interference in differentiated human adipocytes (at day 14) using single-cell transcriptomic profiling. CRISPRi suppressed *CITED2* expression by a log2 fold change of −0.69 (FDR =2.27E-15), confirming effective target repression. Knocking down *CITED2* in adipocytes coordinately suppressed well-established PPARγ-driven programs related to adipogenesis, lipid metabolism and mitochondrial function, comprising leading-edge genes including *PPARG*, *ADIPOQ*, *LPL*, *CD36*, *FABP4* and *SCD*, all of which are canonical markers of mature adipocytes^22,93^ (**Supplementary Table S31**). We additionally observed a reciprocal induction of gene programs related to inflammatory response and extracellular matrix remodeling (**Fig. 6l**, **Supplementary Table S31**). These results are consistent with *CITED2* acting as a CBP/p300 co-activator of PPARγ^94^. Notably, CBP/p300 has been demonstrated to be required for PPARγ-driven adipocyte differentiation^95^, supporting our results providing direct mechanistic basis for the SMR-inferred protective direction, demonstrating that reduced *CITED2* expression dismantles key cellular programs controlling adipocyte function.

Together, these analyses show that the genetic regulation underlying cardiometabolic traits acts largely through cell-state- and context-specific chromatin and expression effects, which integrative Mendelian randomization resolves into directional regulatory cascades.

### Cellular program QTLs link genetic variation to adipocyte function and metabolic disease

Metabolic diseases are regulated by aggregated polygenic effects, which can be probed through *trans*-eQTL mapping. However, individual *trans*-eQTL discovery is often imprecise and statistically demanding due to large sample size requirements, but these challenges can be mitigated by analyzing such polygenic effects at the level of broader cellular programs (PMC11019359). To investigate whether genetic variants influence biological processes central to adipocyte function and metabolic disease, we implemented a cellular program QTL (cpQTL) mapping strategy **(Fig. 7a, Methods)**. To this end, we defined three key categories of adipocyte biology: energy metabolism, adaptive responses, and regulation of differentiation, each encompassing a set of interpretable cellular programs. Importantly, these cellular programs were responsive to FFA, low-glucose, and hypoxia exposure, demonstrating their functional relevance in our experimental system **(Fig. 2a; Extended Data Fig. 3a and b)**. By generating PCA-derived program-specific gene expression signatures, we mapped cpQTLs and linked significant signals to metabolic disease-associated PheWAS traits **(Fig. 7a, Methods)**.

**Fig. 7.**
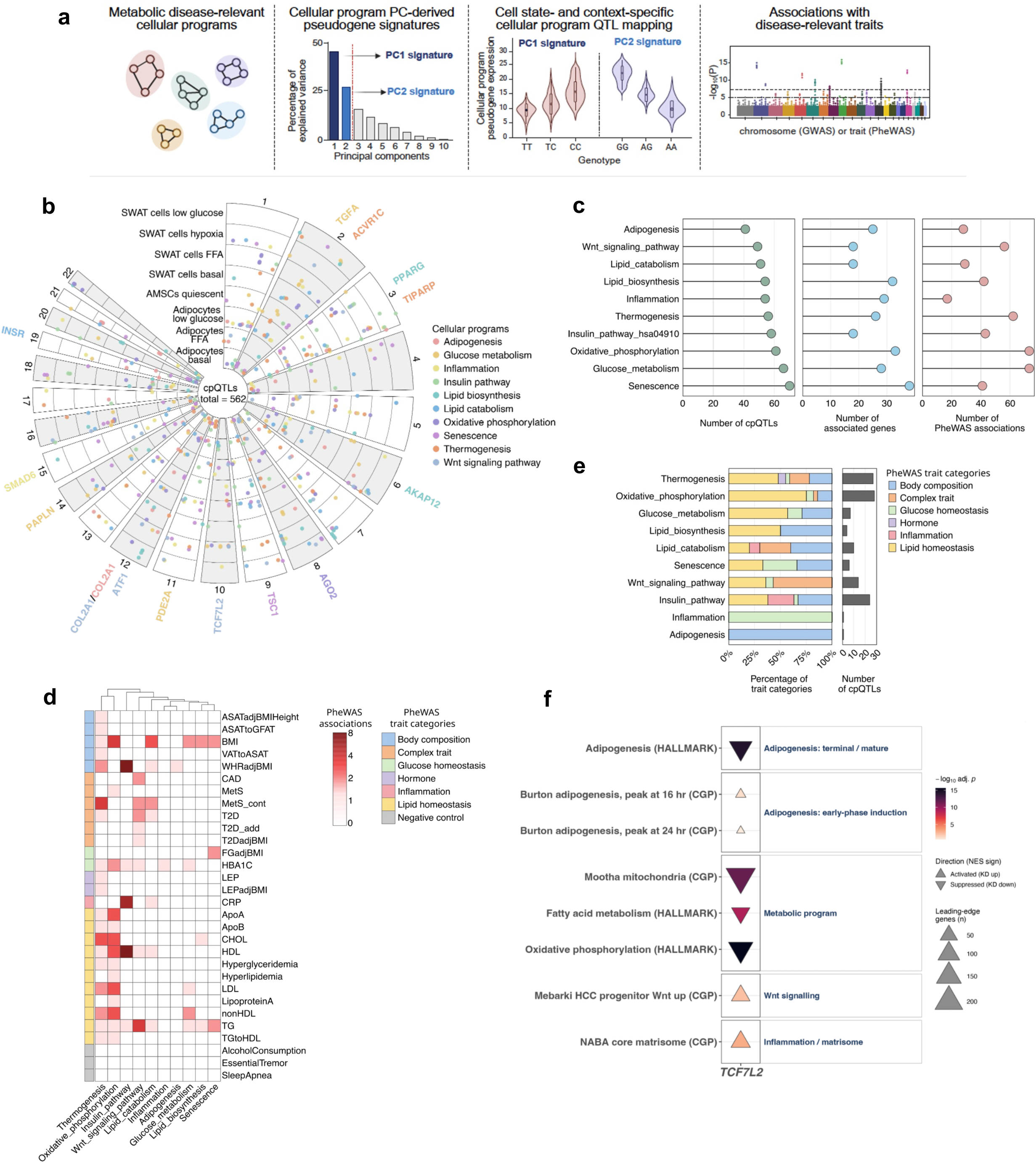
Cellular program QTLs link to metabolic disease mechanisms. **a**, Schematic of cpQTL mapping strategy. **b**, Circular Manhattan plot showing all cpQTLs across cell states and cellular programs after LD clumping, with representative predicted causal genes highlighted. **c**, Summary of LD-clumped cpQTL counts (left panel), predicted causal genes (middle panel), and PheWAS associations (right panel) across cellular programs. **d**, Heatmap of cpQTL associations with cardiometabolic disease-related PheWAS traits (p-value < 5 × 10^-8^) after LD clumping. **e**, Proportion of cardiometabolic disease-related trait categories across cellular programs (left panel), and the total number of associated cpQTL (right panel). **f.** Gene-set enrichment (GSEA) of the genome-wide *trans* effects of *TCF7L2* knockdown by CRISPRi in differentiating human adipocytes, relative to non-targeting controls. Triangle orientation denotes NES sign (up, activated on KD; down, suppressed on KD); fill color, −log₁₀ adjusted *p*; size, number of leading-edge genes (GSEA FDR < 0.25).

We discovered 811 cpQTLs across cellular programs, cell states and stimuli, the majority of which were located within gene introns (54.6%), distal intergenic regions (24.3%) and promoters (17.6%) (p-value < 5 × 10^-6^; **Extended Data Fig. 9a and Supplementary Table 32**). It should be noted that the number of cells per donor for AMSC proliferating and adipocytes under hypoxia was insufficient for robust analysis, and these conditions were therefore excluded. We next applied LD clumping to reduce redundancy among variants, resulting in 522 cpQTLs and revealing a high degree of context specificity, with over 90% of cpQTLs being cell state- and stimulus-specific (**Fig. 7b and Extended Data Fig. 9b**). This systematic catalog of context-specific cpQTL mapping in primary adipocyte villages comprises 41-70 cpQTLs across cellular programs, associated with 18-38 predicted causal genes and 17-73 PheWAS traits (**Fig. 7c**). Remarkably, when focusing on cardiometabolic disease-related PheWAS traits (P < 5 × 10⁻⁸), we identified multiple associations with cpQTLs, with traits such as BMI, WHRadjBMI, HbA1c, HDL, and TG exhibiting several associations across multiple cellular programs (**Fig. 7d**). Among all cellular programs, thermogenesis, oxidative phosphorylation and the insulin pathway showed the highest number of cpQTLs associations to PheWAS traits **(Fig. 7e)**. Interestingly, thermogenesis and oxidative phosphorylation were predominantly associated with lipid homeostasis-related traits, whereas the insulin signaling pathway exhibited a more balanced distribution of associations across traits related to lipid metabolism, inflammation, and body composition (**Fig. 7e**). Therefore, combining cpQTL mapping with a stringent phenome-wide statistical threshold provides robust evidence for variants with potential regulatory effects on disease-relevant phenotypes.

Among the predicted causal genes associated with cpQTLs, we identified several with established roles in regulating cellular programs central to metabolic homeostasis and adipocyte function, including *INSR* (rs3786680), *TIPARP* (rs900399), *TSC1* (rs4962081), *PDE2A* (rs341061), *TCF7L2* (rs11196236) and *PPARG* (rs4256108) (**Fig. 7b**). Of particular interest, we found that, under low-glucose conditions in adipocytes, the risk T/T allele of the cpQTL rs4556933 reduced thermogenesis program expression (*P* = 1.40 × 10^-7^, b = −0.79) and was associated with lower HbA1c, total cholesterol, and LDL levels **(Extended Data Fig. 9c and d)**. This variant, a GWAS signal for total cholesterol, resides within an intron of *ACVR1C* **(Extended Data Fig. 9e and f)**, which was further implicated as the causal gene by cS2G analysis^96^ (**Supplementary Table 32**) and supported by eQTL data from GTEx and Open Targets. Importantly, we also found that the cpQTLs rs3742825 (*P* = 6.47 × 10^-7^, b = −0.89) and rs12587806 (P = 3.23 × 10^-8^, b = −0.96), which regulate the glucose-metabolism program in SWAT cells exposed to hypoxia (**Extended Data Fig. 9g**) are colocalized (PP.H4 = 0.79) with the HbA1c GWAS signal (**Extended Data Fig. 9h**). Causal gene analysis using cS2G predicted PAPLN (rs3742825) and PSEN1 (rs12587806) as candidate effector genes, suggesting previously unrecognized roles in the regulation of glucose metabolism under hypoxic conditions, in addition to their established functions in extracellular matrix organization and Notch signaling^97,98^, respectively.

To test whether cpQTL-nominated genes are functional regulators of their corresponding cellular programs, we examined the effects of CRISPRi-mediated gene silencing on transcriptomic remodeling in differentiated human adipocytes using scRNA-seq readout. We focused on *TCF7L2*, a predicted cpQTL causal genes nominated in low-glucose exposed adipocytes corresponding to *TCF7L2* linked to the Wnt-signalling. *TCF7L2* has a strong link to T2D and adipocyte function as described above. CRISPRi reduced *TCF7L2* expression by log2 fold change of −0.25 (FDR = 0.014), confirming effective target repression. Interestingly, *TCF7L2* knockdown upregulated the transcriptional signature of early adipogenesis, whereas it produced a broad, coordinated suppression of the terminal adipogenesis and metabolic processes, including oxidative phosphorylation and fatty-acid metabolism (**Fig. 7f, Supplementary Tables S34**). Moreover, we observed an induction of Wnt-associated signatures, consistent with *TCF7L2*’s function as a canonical Wnt/β-catenin effector required for adipocyte differentiation^99^ and acting as a positive regulator of the mature adipocyte state. CRISPRi-mediated knockdown of *TCF7L2* showed a diverged regulation of the extracellular-matrix compartments. Together, silencing this cpQTL-nominated gene reshaped core adipocyte programs, providing orthogonal experimental support that cpQTLs pinpoint functional regulators of adipocyte biology (GSEA FDR < 0.25 throughout; **Fig. 7f, Supplementary Tables S33**).

Altogether, our data demonstrate that cpQTL mapping in cell villages is a powerful approach for uncovering how genetic variants modulate key cellular programs underlying metabolic disease in a context-specific manner.

## Discussion

In this study, we present a paired single-nucleus gene expression and chromatin accessibility atlas of human primary adipocyte villages under metabolic disease-relevant stimuli from 118 donors. By linking heritability of metabolic disease relevant traits and genetic variants to specific cell states and contexts, we demonstrate that the adipocyte village platform offers a robust and scalable framework for V2F discovery. The approach implemented in this study explains 13.2-75 % of genome-wide significant GWAS loci across 13 cardiometabolic disease-relevant traits through colocalized molQTLs, and identifies thousands of putative specific and pleiotropic causal variants linking molecular phenotypes to disease, underscoring the importance of modeling disease-relevant cell states under controlled experimental conditions.

Polygenic metabolic traits are shaped by both genetic and environmental factors, making it essential to examine molecular phenotypes within disease-relevant cellular contexts to uncover pathophysiological mechanisms. In addition to mapping trait heritability to the most relevant cell states, we observed context-dependent heritability enrichment along the adipocyte and SWAT cell differentiation axis, highlighting dynamic genetic risk enrichment across these distinct trajectories. The observation that metabolic disease risk heritability is positively associated with the adipogenic and SWAT differentiation trajectories, with the strongest coupling observed under disease-like stimulations such as FFA and hypoxia. Notably, this positive association was attenuated in SWAT cells under low glucose exposure, indicating that healthy glucose concentration shifts cells toward a protective, healthy-like state in which genetic disease risk becomes decoupled from the differentiation trajectory.

Most existing eQTL studies in adipose tissue have been conducted at the bulk-tissue level, lacking the resolution to capture cell type-specific regulatory effects^100–107^. While population-scale single-cell genomics offers a path forward, most single-cell QTL studies still rely on pseudobulk aggregation, which can obscure both cell type- and context-specific signals^108–112^. Additionally, incorporating context into genetic modeling by jointly modeling genotype main effects and genotype-by-context interactions at the single cell resolution enables the detection of interactive and dynamic eQTLs^113^ (CASTIE, Liu *et al*., 2026). Using this approach, we identified 1.6-fold more eQTLs compared to the pseudobulk strategy. It is well known that eQTL can only explain a small proportion of GWAS signals^111,111,114–116^, with our findings revealing that incorporating metabolic disease-relevant contexts yielded 2- to 5-fold more colocalizations than basal-only eQTL mapping. A recurring obstacle in connecting eQTLs to disease is that these two categories have systematically different genomic architectures. Standard cis-eQTLs cluster tightly around transcription start sites (TSS), whereas GWAS variants for complex traits lie farther from the TSS, near genes under stronger selective constraint with more complex regulatory landscapes^115^. Because conventional eQTL mapping is dominated by these promoter-proximal effects, it preferentially misses the distal regulatory variants that underlie most trait associations. Our context-dependent dynamic QTLs depart from this basal architecture in exactly that direction, promoter-proximal representation falls from ∼37% at baseline to ∼18% among dynamic QTLs, with reciprocal enrichment in intronic and distal intergenic regions, these results indicate that stimulation uncovers a more distal, GWAS-like regulatory variant that basal mapping does not capture.

By capturing these additional dynamic, context-dependent signals, our approach explained a greater fraction of GWAS loci, underscoring the value of context-resolved single-cell QTL mapping. Consistent with previous studies in other cell types^73,117,118^, we identified substantially more cis-caQTLs than cis-eQTLs, which may reflect both the larger number of chromatin accessible regions compared to expressed genes and the higher proximity between genetic variants to chromatin accessible regions than genes. Notably, caQTLs explained a larger proportion of GWAS colocalized signals than eQTLs, with a subset of loci supported by both caQTLs and eQTLs. This aligns with prior observations that eQTLs alone are often insufficient to resolve the regulatory architecture of complex traits. In contrast, caQTLs can reveal noncoding regulatory DNA regions enriched for disease-associated variants, offering critical insights into noncoding GWAS signals^118–120^. These loci exhibit the diverse regulatory mechanisms between functional variants and their molecular phenotypes, highlighting the need for cell state- and context-specific QTL discovery. The GWAS signals colocalized with a low glucose-specific eQTL modulating *NCOR2* expression highlights the importance of metabolic context in revealing the associations between regulatory variants and disease risk. In contrast, the *TIPARP* locus illustrates a fundamental regulatory mechanism active across multiple cell states and stimuli, with robust colocalizations observed across both transcriptomic and chromatin accessibility layers. The *PTGER3* region represents a case in which transcriptional regulation occurs in the absence of detectable chromatin accessibility variation: colocalization was observed only at the eQTL level, suggesting a gene-proximal regulatory mechanism independent of chromatin remodeling. Together, these examples demonstrate the value of integrating multiple molecular layers with single-cell and context-specific resolution to disentangle the complex mechanisms by which regulatory variants shape the architecture of cardiometabolic traits. Critically, different metabolic contexts reveal different biology: the healthy state (low glucose) uniquely captures genetic variants affecting insulin sensitivity and favorable fat distribution, as exemplified by the *NCOR2* locus, while disease states (nutrient overload, hypoxia) reveal pathological mechanisms. This approach, profiling differentiated adipocytes rather than progenitors, and modeling multiple metabolic states representing both health and disease, provides a framework for connecting genetic variation to the complete spectrum of disease-relevant cellular mechanisms. By layering SMR onto colocalization, we moved beyond identifying shared causal variants to ordering them into directional chromatin-to-expression-to-trait cascades, illustrating how integrative Mendelian randomization can nominate not only effector genes but the regulatory logic linking them to disease.

Another key advance of our study was the implementation of cpQTL mapping to link genetic variants to interpretable pathophysiological processes. All selected cellular programs were dynamically modulated by stimuli resembling both healthy- and disease-like states, recapitulating, for example, the impairments in oxidative metabolism observed in human obesity^121,122^. This approach demonstrates a high degree of context specificity, revealing an extensive repertoire of cpQTLs and identifying over 500 regulatory associations with metabolic disease-relevant cellular programs. By combining causal gene prediction and molecular cis-eQTL data, our results identify several genes with key regulatory functions, including canonical regulators of pathways central to adipocyte biology, such as insulin and mTORC1 signaling (*INSR* and *TSC1*) and adipogenesis (*PPARG*). Further highlighting the power of this approach for therapeutic target discovery, we identified rs4556933 as a representative cpQTL linked to thermogenesis-associated programs under low-glucose conditions. This cpQTL is associated with a favorable glucose and lipid profile, with *ACVR1C* predicted as the causal gene and implicated in adipogenic transcription, catecholamine responsiveness, and thermogenesis^123–125^. Importantly, *ACVR1C* is currently under investigation as a therapeutic target for obesity and type 2 diabetes, with clinical development being pursued by Arrowhead Pharmaceuticals (clinical trial ID: NCT06937203). Additionally, the colocalization of context-specific cpQTLs with GWAS signals provides preliminary validation of this framework and highlights its potential to reveal genetically anchored pathophysiological mechanisms. In support of this, we identified cpQTLs (lead variants: rs3742825 and rs12587806) associated with reduced expression of a glucose metabolism program in SWAT cells specifically under hypoxia, with their significant colocalization with HbA1c GWAS further suggesting impaired systemic glucose homeostasis. Collectively, these results establish a framework for interpreting genetic variants that converge on cellular programs, in which human adipocyte villages provide a powerful strategy to uncover cell state- and context-specific mechanisms relevant to metabolic disease. The cpQTLs identified in this study therefore provide a foundation for systematically dissecting the mechanisms by which these variants regulate fundamental processes in adipocyte biology, while also providing genetic support for prioritizing therapeutic targets.

CRISPRi in differentiated human adipocytes experimentally validated effector genes from both arms of our framework: *CITED2*, nominated by colocalization and directionally inferred by SMR, and *TCF7L2*, nominated by a context-aware cellular program QTL. In both cases, silencing these targets reshaped core adipocyte differentiation programs in the direction predicted by the genetic nomination, demonstrating that variant-level and program-level discovery pinpoint genes whose perturbation reshapes core adipocyte biology.

Our data encompasses evidence from disease heritability enrichment and multiomic molQTL mapping, emphasizing context-dependency as a key driver of pathophysiological mechanisms. Combining metabolic disease-relevant stimuli in human primary adipocyte villages with snMultiome profiling provides a powerful strategy to link genetic variants to specific contexts and cellular programs that may inform targeted therapies. A limitation of this study is the modest sample size and low representation of certain cell populations, which constrained statistical power for certain analyses. Nonetheless, our work lays the foundation for population-scale profiling of adipocyte villages. Expanding this approach to additional depots is also an important step toward uncovering the genetic mechanisms underlying depot-specific contributions to whole-body metabolic health. In conclusion, using a physiologically relevant cellular model, we implemented a scalable natural genetic variation screening strategy that accelerates the discovery of genetically anchored mechanistic insights with therapeutic potential for metabolic diseases.

## Methods

### Subjects and primary AMSC isolation

Abdominal subcutaneous adipose tissues were collected from 118 donors (47 males and 71 females) undergoing clinically indicated scheduled surgery at the Technical University of Munich (Munich, Germany), the University of Hohenheim (Stuttgart, Germany) and Weight Center at Massachusetts General Hospital (Boston MA, USA) (Supplementary Table 1). Prior to the sample collection each participant was informed, and written consent was obtained. The studies were approved by the ethics committees at the Faculty of Medicine of the Technical University of Munich (study number: 5716/13) and Massachusetts General Hospital (IRB protocol:2022P000669). No patient information was shared outside of MGH, and participants’ identities remained anonymous.

Adipose tissue biopsies were processed on the day they were received to isolate adipose mesenchymal stem cells (AMSCs). AMSC isolation and quality control was performed as previously described^34,126,127^. Briefly, biopsies were dissected, minced, and treated with Krebs-ringer phosphate buffer (KRP-pH 7.4) containing 4 % bovine serum albumin (Sigma A7906) and 200 u/ml collagenase (Cell Signaling 44204S) for one hour at 37°C under strong agitation in a water bath. After digestion, the sample was centrifuged at 200g for 10min. The supernatant containing mostly ruptured lipids from mature adipocytes was removed and the pellet was resuspended in the Isolation media (DMEM/F12 Gibco 31330-038, 1% penicillin-streptomycin Sigma Aldrich P0781 and 10% FBS Sigma Aldrich F2442) and filtered through 70uM cell strainer. Subsequently, cells were seeded into appropriate plates or flasks (e.g. cells isolated from 1g fat seeded in 2 wells of 12 well plate). AMSCs were grown in an incubator maintained at 37°C temperature and at 5% CO2. On the next day, cells were washed three times with phosphate-buffered saline (PBS pH-7.4) and the medium was replaced to proliferation medium (DMEM/F12 Gibco 31330-038, 1% penicillin-streptomycin Sigma Aldrich P0781, 2.5% FBS Sigma Aldrich F2442, 17μM D-Pantothenic acid hemicalcium Sigma Aldrich P5155, 33μM (+) D-Biotin Sigma Aldrich 2031, 1ng/ml rhFGF Fisher Scientific 233FB025, 10ng/ml rhEGF Fisher Scientific 236EG200, 0.13μM insulin Sigma Aldrich I9278 and 0.1mg/ml Gentamicin Scientific NC0753440). Following, the medium was changed every three days, and cells were grown to the 90-95% confluency before passaging and cryopreserved and stored in liquid nitrogen for further use.

### Genotyping

Genotyping and data harmonization was performed for 1,275 donors from CellGenBank in longitudinal batches through 2019 to 2024 using the Illumina Infinium Global Screening Array v3.0 (GSA) (Illumina, Inc. Infinium Global Screening Array Data Sheet. San Diego, CA) for the 2019, 2021, and 2022 cohorts, and the Illumina Diversity Array (GDA) for the 2024 cohort. Each cohort was individually imputed using the TOPMed r3 reference panel [<u>REF</u>, <u>REF</u>, <u>REF</u>]. To integrate these different array data, raw genotyping data underwent harmonization using the BCFtools suite (v1.9). Strand orientation was performed to the hg38 reference genome (bcftools +fixref plugin), which diagnosed and corrected strand flips to ensure alignment consistency across batches. Subsequently, variants were normalized (bcftools norm) to left-align insertions or deletion and spit multiallelic sites to biallelic records. The harmonized data was aggregated (bcftools merge) and concatenated across chromosomes (bcftools concat). Finally, quality control filtering (bcftools filter) was applied to exclude low-quality markers, removing variants with a call rate < 95%, MAF < 1%, and Hardy-Weinberg equilibrium (P < 1 x 10e-6). The final dataset comprised 8,292,147 SNPs.

### Cell culture and generation of AMSC villages

Primary AMSCs were cultured and differentiated as previously described^126^, with the following modifications. Cells were maintained in proliferation media (PM: DMEM/F-12 supplemented with 1% Penicillin-Streptomycin, 33 μM Biotin, 17 μM Pantothenate, 0.13 μM Insulin, 10 ng/ml EGF, 1 ng/ml FGF and 2.5% FBS) and differentiation was initiated in fully confluent cells by changing PM to induction media (IM: DMEM/F-12 supplemented with 1% Penicillin-Streptomycin, 33 μM Biotin, 17 μM Pantothenate, 0.86 μM Insulin, 1 nM T3, 0.1 μM Hydrocortisone, 0.01 mg/ml Transferrin, 0.17 % fatty acid free BSA, 2% FBS, 1 μM Rosiglitazone, 25 nM Dexamethasone and 0.25 mM IBMX). Following 3 days, IM was changed to differentiation media (DM: DMEM/F-12 supplemented with 1% Penicillin-Streptomycin, 33 μM Biotin, 17 μM Pantothenate, 0.86 μM Insulin, 1 nM T3, 0.1 μM Hydrocortisone, 0.01 mg/ml Transferrin, 0.17 % fatty acid free BSA and 2% FBS) for a total period of 14 days. Differentiation was carried out under normoxia (21% O_2_) with the standard high glucose (17.5 mM) media described above and following exposure to hypoxia (4% O_2_), free fatty acids (0.1 mM oleic and linoleic acid in 10% fatty acid free BSA) or low glucose media (5 mM in a 1:1 mixture of glucose-free DMEM and Ham’s F-12) throughout the entire differentiation time course. In addition, a subcutaneous immortalized AMSC (iAMSC) cell line was added to every village as an internal control to assess batch correction efficiency (see Supplementary Notes). iAMSCs were maintained in growth media (DMEM supplemented with 1% Penicillin-Streptomycin and 10% FBS). After pooling into AMSC villages, cells were induced and differentiated as described above.

Four individual AMSC villages were generated using cells from a total of 118 donors that were randomly selected to ensure overall balanced representation across sex, age, BMI, and predicted individual polygenic risk scores^128^ (**Supplemental Table S1** for the full PRS set we used for selection). Cells from each donor were cultured in PM independently in a T-75 flask until around 80% confluent, at which point AMSCs from 20 to 40 donors were pooled at equal proportions and seeded at a density of 400,000 cells per well of a 6 well plate. This seeding density allowed to decrease the proliferation period and thus minimize donor dropouts by the timely differentiation of fully confluent AMSC villages.

### Nuclei isolation, snMultiome library generation and sequencing

Nuclei were isolated at day 0 and 14 of differentiation under the corresponding experimental conditions. First, AMSCs were washed once with pre-warmed PBS and incubated with 2 ml per well of serum-free DMEM/F12 supplemented with 50 μg/ml DNase I (Stem Cell Tech, #07900) for 10 min at 37°C in a cell culture incubator. Next, cells were placed on ice, washed twice with ice-cold PBS and collected in 500 µl of ice-cold Nuclei Lysis Buffer (5 mM CaCl2, 3 mM Mg(Ac)2, 10 mM Tris pH 7.5, 320 mM Sucrose, 0.1 mM mM EDTA, 0.1% IGEPAL CA-630, 1 mM DTT and 1 U/μl RNase inhibitor) into a pre-chilled 1.5 ml low protein binding tube (Eppendorf, #022431081). Samples were mechanically disrupted by pipetting 15 times, followed by 3 min incubation on ice and subsequent 15 strokes with a tight pestle in a glass Dounce homogenizer on ice. The resulting cell lysate was transferred to a 5 ml tube on ice, ice-cold Wash Buffer (1X PBS, 2% BSA and 1 U/μl RNase inhibitor) was added for a 4 ml final volume and mixed gently by inverting the tube five times. Samples were next centrifuged for 5 min at 500 rcf at 4°C in a swinging bucket centrifuge with the acceleration and brake set at 50% of the maximal setting. The supernatant was discarded; the nuclei pellet was resuspended in 2 ml of Freezing Buffer (Wash Buffer supplemented with 10% DMSO) and 1 ml aliquots transferred to a cryovial for overnight freezing in a freezing container at −80°C. For each AMSC village, the full set of samples were thawed in a room temperature water bath and filtered through a 30 μm cell strainer (Sysmex America, #04-004-2326) into a pre-chilled 1.5 ml low protein binding tube on ice. Nuclei were permeabilized with 0.5% Tween-20 and 0.4% Digitonin, followed by pipette mixing five times and incubation on ice for 10 min. Next, nuclei were washed by centrifugation for 5 min at 500 rcf at 4°C in a swinging bucket centrifuge with the acceleration and brake set at 50% of the maximal setting. The supernatant was discarded, the nuclei pellet was resuspended in 150 μl of 1X Nuclei Buffer (10x Genomics, PN-2000207), visually inspected and counted in an hemocytometer using trypan blue staining. Samples were then centrifuged for 5 min at 500 rcf at 4°C in a swinging bucket centrifuge with the acceleration and brake set at 50% of the maximal setting, supernatant was discarded and nuclei pellet was resuspended in appropriate volume of 1X Nuclei Buffer to immediately proceed with the snMultiome protocol.

snMultiome was performed with the Chromium Next GEM Single Cell Multiome ATAC + Gene Expression kit (10x Genomics, PN-1000283; protocol version CG000338 Rev F) according to the manufacturer’s instructions, targeting a recovery of 20,000 to 30,000 nuclei per sample. Both snRNA-seq and snATAC-seq libraries were quantified by KAPA qPCR (Roche. #07960140001), while fragment size distribution was assessed using a TapeStation (Agilent). Sequencing was performed on an Illumina NovaSeq X targeting a minimum of 20,000 and 25,000 read pairs per nucleus for snRNA-seq and snATAC-seq, respectively. Base calling was conducted using Illumina’s RTA3 software, and the BCL Convert (v4.1.7, Illumina, USA) was employed for generating FASTQ files.

### snMultiome data genetic demultiplexing, quality control, processing and annotation

Raw FASTQ data of each sample were then processed with Cell Ranger ARC (v2.0.2) function “cellranger-arc count” with default parameters and mapped to the human GRCh38 genome (cellranger pre-build GRCh38-2020-A) to generate unique molecular identifier (UMI) expression matrices and fragment files. Donor genotype demultiplexing was performed using Dropulation^30^ on the Cell Ranger filtered expression matrices, donor genotype was filtered for common SNPs (minor allele frequency > 1%) and SNPs overlapping genes. The expression matrix of each sample was subsetted by removing the empty droplets or the droplets that contain multiple donorIDs. Nuclei with fewer than 200 detected genes or more than 15% of reads mapping to mitochondrial genes were also removed for RNA modality. Nuclei with TSS <10 or ATAC counts <5000 and >2,000,000 were removed from the ATAC modality. Individual samples after quality control were first processed to inspect data structure and distribution, with RNA and ATAC modalities separately [RNA processing: Seurat^129^ (v5.0.0), ATAC processing: SnapATAC2^130^ (v2.6.1). Subsequently, SCTransform normalization (v2) in Seurat was performed separately for each sample on high-quality nuclei from RNA modality, with the number of UMIs per nuclei regressed out. Principal components (PCs) were identified using the ‘RunPCA’ function for each sample. RPCA integration was then performed using the one-liner integration function “IntegrateLayers”, correcting for the village batch effect. Next, “FindNeighbors” was performed and followed by “FindClusters” with different resolutions to generate cluster coordinates. UMAP were generated by “RunUMAP’ for visualization, with the first 30 dimensions from the integrated rpca. We annotated the major cell states using previously published marker genes^22,24,25^.

For ATAC modality after quality control, a 500-bp genome-wide tile matrix was constructed for each library using snap.pp.add_tile_matrix(), and the top 250,000 most accessible bins were selected as features with snap.pp.select_features(). High-quality ATAC nuclei were then intersected with the annotated high-quality RNA nuclei to allow the major cell state labels to be transferred to the ATAC modality for the paired nuclei barcodes. Cell-type level peak calling for each major cell state under each stimulation was performed using the “snap.tl.macs3” function in SnapATAC2. After peak calling, tl.merge_peaks() function was used to generate unified, non-overlapping, and fixed-width peaks for downstream analysis. Genomic annotation was performed using the TxDb.Hsapiens.UCSC.hg38.knownGene transcript database and org.Hs.eg.db for gene symbol mapping. Peaks were classified relative to genomic features, including promoters (±3 kb from transcription start sites), exons, introns, and intergenic regions.

### Reference mapping

Symphony^48^ is used for reference mapping between our dataset and published in vivo adipose tissue and *in vitro* adipocyte single-nucleus RNA-seq datasets^22,24^. We extracted the expression matrix and metadata for building Symphony references for reference datasets. We first subsetted the data to include only the subcutaneous depot to match our dataset. For reference building, we selected the 2000 most variable genes and subsetted the dataset using these variable genes, performed PCA with 20 dimensions, and calculated the UMAP. For query mapping, we extracted the expression matrix and metadata from the query dataset, then mapped the query dataset onto the reference dataset using the mapQuery() function, with village batch included as a variable to harmonize over. In addition, we transferred the annotated labels from the reference datasets to our data using the knnPredict() function.

### Trajectory analysis

Trajectory analysis was performed using Monocle 3. We first extracted the metadata and expression count matrix from the integrated Seurat object to construct a Monocle3 cds object. The UMAP embeddings and cluster annotations derived from the integrated Seurat object were then transferred to the cds object to ensure consistency. Trajectory analysis was performed using the learn_graph() function, and the resulting trajectories were visualized using plot_cells() function.

### Consensus non-negative matrix factorization (cNMF) analysis

Gene expression programs (GEPs) were identified by consensus non-negative matrix factorization^131^ (cNMF) as implemented in OmicVerse (v1.6.9) with Scanpy as the underlying single-cell framework. Starting from the integrated single-nucleus RNA-seq dataset (337,666 nuclei across the four combined villages), gene counts were normalized with a shifted-logarithm transformation and the 2,000 most variable genes were selected using analytic Pearson residuals. Expression values were then scaled to unit variance and principal component analysis was performed.

cNMF was run across a range of factorization ranks (K = 5-10) using the 2,000 high-variance genes as input. For each value of K, 20 independent NMF replicates were computed. The number of programs was chosen by examining the trade-off between solution stability and reconstruction error across K, from which K = 6 was selected. Consensus GEPs were then derived at K = 6 by clustering the combined replicate spectra and removing outlier components using a local-density threshold of 0.1 prior to consensus estimation.

From the consensus solution, we extracted for each of the six programs the gene spectra (GEP gene scores), the corresponding TPM-scaled spectra, and the top-ranked program genes, together with the per-cell normalized program usages. Each cell was assigned to the program with the highest normalized usage to define cNMF-based clusters. Program usages were visualized on the UMAP embeddings.

### Differential expressed genes (DEGs) analysis

Cell state-specific transcriptomic marker signatures were identified using the FindAllMarkers() function in Seurat, with min.pct = 0.5 and logfc.threshold = 0.75.

Pseudobulk raw counts were generated using the AggregateExpression() function by each donor, stratified by cell state and stimulation condition, with normalization.method = “RC” specified. Differentially expressed genes (DEGs) between pairwise stimulation groups in adipocytes and SWAT cells were then calculated using DESeq2^132^ (R package v1.36.0), including top 5 genetic PCs, age, sex, village batch, and cohort as covariates. Significant DEGs were defined by FDR < 0.05 and |log2 FC| > 0.2.

### Functional pathway enrichment analysis

Enriched pathways for cell state-specific transcriptomic marker signatures (using only genes with adjusted P < 0.05) were determined using enrichR (v3.2) with the following databases: GO Biological Process 2023, MSigDB Hallmark 2020, Reactome 2022, KEGG 2021 Human, and WikiPathways 2023 Human. Pathways with adjusted P < 0.05 were considered significantly enriched.

Gene set enrichment analysis (GSEA) was performed using ClusterProfiler^133^ (R package v4.9.1) and ReactomePA^134^ (v1.40.0). All DEGs between groups (pairwise comparison between stimulatory groups or high vs low scDRS SWAT cell populations) were extracted. After pre-ranking by log2 fold change, DEGs were then fed to gseGO(), gseKEGG() and gsePathway() functions for GO (Ashburner et al., 2000), and KEGG (Kanehisa et al., 2000) and REACTOME (Fabregat et al., 2018) pathways for enrichment analysis. Significantly enriched pathways were defined by Benjamini-Hochberg adj. p-value < 0.05 and |NES| > 1).

### DAP and motif analysis

Cell state specific DAPs were identified using the SnapATAC2 function snap.tl.diff_test() (one-sided test, direction=“positive”, min_log_fc = 0.01), against a background of cells from the other cell types. Transcription factor motif enrichment was then performed on the per-cell-type marker peak sets using snap.tl.motif_enrichment() with the CIS-BP motif database (snap.datasets.cis_bp(unique=True)) against the hg38 genome, tested jointly across all four cell types. Significantly enriched cell-state– specific TF motifs were defined as those with FDR ≤ 0.05 and |log2(fold change)| ≥ 0.5.

To identify transcription factor motifs associated with stimulatory condition-induced chromatin remodeling in Adipogenic and SWAT cells, differentially accessible peaks (DAPs) between paired stimulatory conditions were called with snap.tl.diff_test() in SnapATAC2. Up- and down-regulated DAPs were defined as peaks with adjusted p-value < 0.05 and log2(fold change) ≥ 0.2 or ≤ −0.2, respectively. Motif enrichment was performed on the up- and down-regulated DAP sets using snap.tl.motif_enrichment() with the CIS-BP motif database (snap.datasets.cis_bp(unique=True)). Significantly enriched TF motifs were defined as those with adjusted P < 0.05 and |log2(fold change)|≥ 0.2. Motifs were further filtered based on minimal detectable expression of the corresponding TF genes in matched pseudobulk snRNA-seq data for each experimental condition (CPM > 1), thereby excluding TFs with low or undetectable expression that are unlikely to drive regulatory activity.

### scE2G enhancer-gene linkage analysis

To link candidate enhancers to their target genes in specific cell types, we applied scE2G^52^ (v1.3, https://github.com/EngreitzLab/scE2G) to our dataset. For each cell state under stimuli, we prepared the RNA count matrix and ATAC fragments as input, and ran the scE2G^multiome^ pipeline with default settings. As part of the pipeline, candidate elements were defined by calling peaks with MACS2^135^ (v2.2.9.1) on the pseudobulk ATAC fragments for each cell type (using parameters from the ENCODE-rE2G^136^ ATAC-seq pipeline), taking 500-bp elements centered on peak summits, removing ENCODE blacklisted regions^137^, and adding 500-bp promoter elements centered on each gene’s transcription start site; candidate element-gene pairs were then generated for all elements within 5 Mb of each promoter. Regulatory enhancer-gene interactions were defined as enhancer-gene pairs with an E2G.Score.qnorm greater than 0.177.

Briefly, scE2G^multiome^ is a supervised logistic regression classifier adapted for single-cell multiome data from the ENCODE-rE2G model, and trained on gold-standard CRISPR perturbation data in K562 cells. It integrates six features: (i) the Activity, Responsiveness, and Contact (ARC)-E2G score; quantitative measures of chromatin accessibility at or near (ii) the element and (iii) the promoter; (iv) genomic distance and (v) gene density between the element and promoter; and (vi) promoter class, indicating whether the gene is ubiquitously expressed across cell types. The ARC-E2G score integrates the ABC score with the Kendall correlation between element accessibility and gene expression across single cells, where ABC activity is computed from pseudobulk scATAC-seq data and contact is estimated as an inverse function of genomic distance; this integration improves calibration across different sequencing depths. The score threshold of 0.177 was determined as the value yielding 70% recall when evaluating predictions in K562 cells against CRISPRi-validated enhancer-gene pairs.

### Protein-protein interaction (PPI) analysis

PPI analysis of transcription factors associated with the enriched motifs was performed using STRING^138^ v12.0 (https://string-db.org). The analysis was restricted to the physical subnetwork, with edge thickness indicating interaction confidence, a minimum interaction score of 0.4, and active interaction sources limited to experiments and databases. Clustering was conducted using the Markov Cluster Algorithm (MCL) with an inflation parameter of 3.

### Single-cell disease heritability enrichment analysis

scDRS (single cell disease-relevant score) analysis^59^ (v1.0.2, using default settings) was used to determine which cell state and stimulation was enriched for specific GWAS traits, by quantifying the aggregate expression of putative disease relevant genes prioritized by MAGMA (v1.10) in each cell to generate per cell disease relevant score. The transcriptomic dataset is inspected for deviations from expected expression levels based on 1000 control gene sets with matching mean and variance to the disease gene sets. The “compute-score” function was used to generate the per cell scDRS. To further determine the statistical significance of the scDRS results, we performed group-level analysis using the “perform-downstream” function. We examined the overall significant level of both cell state-disease association and the associations under stimuli (P <0.05), we also computed the fraction of cells with significant scores (FDR <0.2). We selected 17 metabolic disease relevant GWASes including fat distribution traits (BMI, GFATadjBMI, ASATadjBMI, VAT/ASAT ratio, WHRadjBMI), insulin sensitivity and resistance traits (FI, IFCadjBMI, ISIadjBMI, HOMAIRadjBMI), glucose/lipid homeostasis traits (LDL, HDL, Triglyceride, HbA1c, Glucose), as well as complex disease traits (T2D, T1D, CAD) for scDRS analysis. Primary GWAS sources for all traits are listed in **Supplementary Notes**. Furthermore, Pearson correlation between PC1 or PC2 and per cell scDRS of a given GWAS was performed to evaluate whether the adipogenic or AMSC/SWAT differentiation trajectories are significantly associated with specific disease traits.

### Stratified LD-score regression (sLDSC) analysis

s-LDSC analysis was performed using ldsc (v1.0.1) as described in Finucane et al. to partition the heritability of the top 100,000 cell state-representative ATAC peaks. ATAC peaks are first pseudobulked by cell state under their relevant stimulations. GWAS summary statistics were obtained from meta-analyses of a subset of metabolic traits (as detailed in the scDRS section). Summary statistics were preprocessed using the “munge_sumstats.py” script. Annotation and LD score files were generated using the make_annot.py and ldsc.py scripts included in the LDSC package, applying default parameters. Heritability (h2g) was estimated by solving the LD score regression equation. We evaluated whether GWAS signal was significantly enriched in loci defined by cell state- and stimulus-specific ATAC peaks, while controlling for the 96 categories of the baseline model (v2.2). The baseline model measures non-specific enrichment for gene proximity, histone marks, open chromatin and enhancer regions. The ldsc.py script was run with the --h2 flag and default settings.Enrichment between the cell state- and stimuli-specific peaks and the GWAS traits was considered significant at both P < 0.05 and Bonferroni-adjusted FDR < 0.05.

s-LDSC (v3.0.1) was also used to partition the heritability of static and context-dependent eQTLs. We constructed separate S-LDSC annotations for each selected context and for static effects. Each context-specific annotation included all cis-variants linked to significant eGenes in that context. Analyses were restricted to context–cell-state combinations with the largest eGene sets, FFA and low glucose responses in SWAT cells, and FFA, low glucose and hypoxia responses in adipocytes, to preserve power and avoid unstable estimates from contexts with few eGenes, such as donor age and sex. For comparison, we generated a static annotation for each cell state (AMSCs, SWAT cells and adipocytes) using all cis-variants of static eGenes and the same *cis*-window definition. Each annotation was intersected with the 1000 Genomes Phase 3 European reference panel (GRCh38), and LD scores were calculated in 1-cM windows using HapMap3 variants. GWAS summary statistics for 13 metabolic traits were harmonized to the HapMap3 variant set and processed with ‘munge_sumstats.py’. Partitioned heritability was then estimated for each annotation separately, conditional on the baseline-LD model v2.2, using ‘--overlap-annot’ and ‘--print-coefficients’. For each annotation-trait pair, we recorded heritability enrichment and the standardized coefficient (τ), which quantifies the annotation-specific contribution conditional on the baseline model. *P* values for enrichment and τ were corrected across traits within each annotation using the Benjamini–Hochberg procedure. Interpretation was restricted to traits with well-estimated total heritability (total h² z-score > 4), retaining 11 of 13 traits.

### Pseudobulk genome-wide cis-eQTL and caQTL mapping

Pseudobulk cis-eQTL and cis-caQTL mapping was performed using TensorQTL^139^, stratified by cell state and stimulus. Single-nucleus profiles were first aggregated into pseudobulk samples by each donor × cell state × stimulus, and samples derived from fewer than 100 cells were excluded. For expression, gene counts were CPM-normalized per cell and averaged across cells within each pseudobulk sample. For chromatin accessibility, fragment counts were summed across cells within each pseudobulk sample, the top 100,000 most accessible peaks were retained for analysis, and peak counts were normalized to the sum of reads in peaks per sample. Expression and accessibility counts were inverse-normal transformed (INT) within each village batch to normalize their distributions, and the transformed values were merged across batches for QTL mapping. Proliferating AMSCs and hypoxia-exposed adipocytes were excluded due to insufficient sample sizes (43 and 20 after pseudobulking, respectively).

For eQTLs, the cis-window was defined as ±1 Mb around the gene transcription start site; for caQTL discovery, a ±10 kb cis-window around the peak was used. For downstream colocalization and SMR analyses, caQTL summary statistics were re-computed using a ±1 Mb cis-window to capture the full regulatory neighborhood and to match the cis-window used for eQTL and GWAS summary statistics. For eQTL mapping, models included the top five PEER factors, the top five genotype principal components, and donor age, sex, village batch, and cohort; for caQTL mapping, the same covariate set was used, with the top five chromatin accessibility principal components in place of PEER factors. Genome-wide significant cis-QTLs were identified using the permutation pass of tensorQTL (cis mode), which yields an empirical beta-approximated P value for the top variant per gene or peak. For eGenes, P values were corrected across all tested genes using Storey’s q-value, and significance was defined at q < 0.05. For caPeaks, P values were corrected across all tested peaks using the Benjamini-Hochberg procedure, and significance was defined at BH-adjusted FDR < 0.05.

### Multivariate analysis (mashr) of e/caQTL effects

To characterize sharing and condition specificity of caQTL effects across cell states and stimuli, we applied the multivariate adaptive shrinkage method mashr^66^ (v0.2.79). Because the majority of caPeaks were detected in only a subset of conditions (with ∼37% shared across all eight conditions), restricting the analysis to the intersection would have discarded most of the data; we therefore defined the analysis universe as the union of caPeaks detected in any condition (n = 153,146). For each caPeak, we identified its lead variant as the SNP with the lowest nominal P value across all eight conditions, and constructed matrices of effect sizes (B^) and standard errors (Ŝ) by extracting the slope and standard error of this lead variant in every condition. When a peak was not tested in a given condition, the corresponding B^ entry was set to 0 and the Ŝ entry to 10.

To estimate the null correlation structure across conditions, we constructed a random-test set by sampling 100,000 SNP-peak pairs uniformly from the full set of tested cis pairs. The null correlation matrix V^ was estimated from this random set using estimate_null_correlation_simple. Data-driven covariance matrices were learned from the strong set using principal-components analysis (cov_pca, top five components) followed by extreme deconvolution (cov_ed); canonical covariance matrices were generated using cov_canonical. The mashr model was fitted on the random set with the union of canonical and data-driven covariances, and posterior summaries (posterior mean, posterior standard deviation, local false sign rate [LFSR]) were computed for each peak-condition pair in the strong set using the fitted model with fixed mixture proportions (fixg = TRUE). caQTL effects were considered significant in a given condition at LFSR < 0.05. This setup is conservative with respect to detecting condition-specific signals, as caQTLs measured in only a subset of conditions are penalized toward the null in untested conditions.

### Single-cell context cis-eQTL mapping

Context specific cis eQTL mapping was performed using CASTIE (Context Aware Single Cell eQTL Tool; v0.2.5), which fits a Poisson mixed effects model with a log link. Models were fitted separately within each cell state: AMSCs, adipocytes and SWAT cells. For cell *i* from donor *d*, the observed expression count was modeled y_i_ ∼ Poisson(μ_i_), with *μ*_i_ = *exp*(*η*_i_), and log(nCount_RNA_i_) included as an offset. For adipocytes and SWAT cells, the linear predictor was:

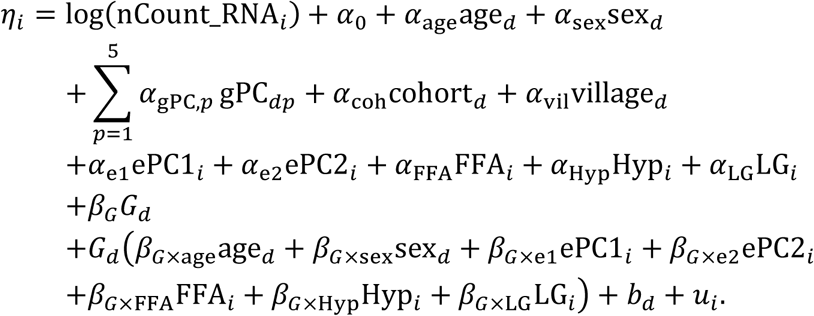

Here, *G_d_* denotes genotype dosage for donor d, *gPC_dp_* denotes the *pth* genetic principal component, ePC1 and ePC2 denote expression principal components, and FFA, Hypoxia and LowGlucose are binary exposure indicators, with basal cells used as the reference condition. bd and ui are random effects. The AMSC model used the same specification but omitted the main effects and genotype interaction terms for FFA, hypoxia and low glucose. Sample level covariates therefore included age, sex, five genetic principal components, cohort and village. Cell-level covariates included two expression principal components and, for adipocytes and SWAT cells, binary indicators for FFA, hypoxia and low glucose. Genotype-by-context terms included G × age, G × sex, G × ePC1 and G × ePC2 in all cell states, with additional G × FFA, G × hypoxia and G × low glucose terms in adipocytes and SWAT cells. Within each cell state, genes expressed in at least 10% of donors were tested against cis variants with minor allele frequency greater than 5%. eGenes were identified at a false discovery rate below 5%. For each cell, the context specific allelic effect was calculated as^80^ βG + Σc βG×Cc C_i_c.

### Single-cell cis-eQTL mapping

As a non-context specific comparison, we performed standard single cell cis eQTL mapping with SAIGE-QTL^112^ (v0.3.5), analyzing each cell type and condition separately. Adipocytes and SWAT cells were analyzed under basal, free fatty acid, low glucose and hypoxia conditions, whereas AMSCs were analyzed without stimulation, yielding nine strata in total. Within each stratum, expression counts were modeled using a Poisson mixed effects model with a log link and a genetic static effect, without genotype by context interaction terms. Covariates included age, sex, the first five genetic principal components, the first two expression principal components, cohort and village, with log transformed total RNA counts included as an offset. Age, sex, genetic principal components, cohort and village were specified at the donor level. Consistent with the context aware analysis, genes expressed in at least 10% of donors were tested against variants with minor allele frequency greater than 5%. Gene level P values for the static genetic effect were calculated using ACAT-V, and eGenes were defined at a false discovery rate below 5% using the qvalue package in R.

### Cellular program QTL (cpQTL) mapping

Cellular programs with well-established roles in adipocyte biology and metabolic disease development were selected from Gene Ontology, Reactome, KEGG, and WikiPathways. These included glucose metabolism (GO0006006 and R-HSA-70326), insulin signaling (hsa04910 and WP481), oxidative phosphorylation (GO0006119 and hsa00190), lipid biosynthesis (GO0006633, GO0046460, and WP357), lipid catabolism (GO0006635 and R-HSA-77289), inflammation (GO0002526 and GO0002544), cellular senescence (GO0090398 and R-HSA-2559583), thermogenesis (GO0120162 and WP4321), adipogenesis (R-HSA-9843745 and WP236), and Wnt signaling (GO0016055, hsa04310, and R-HSA-195721). These cellular programs also met a second selection criterion of being responsive to the stimuli at the transcriptional level in adipocytes and SWAT cells in our experimental system. Each gene within these cellular programs was then verified to be expressed in our cellular model. Prior downstream analysis, each cellular program was manually curated to retain only core genes, thereby excluding upstream and secondary regulators.

Next, for each cellular program, per-donor gene expression values were log-transformed and PCA was used to extract the first two PCs of each program for each donor. The resulting PCA-derived, cellular program-specific pseudogene signatures were used to map genome-wide program trans-QTLs (cpQTLs) (p < 5 × 10^-6^) with TensorQTL, including age, sex, village, cohort and five genetic principal components as covariates. For each identified cpQTL, gene prioritization was performed using cS2G data^96^ using a score cutoff of 0.5, while a rapid distance-based (window-size: +/- 10 kbp) clumping was achieved by an in-house script to identify the most significant variant within the window. Additionally, PheWAS data were obtained from the Common Metabolic Diseases Knowledge Portal (https://md.hugeamp.org) to identify overlaps between cpQTLs and metabolic disease-associated traits (p < 5 × 10^-8^).

### Colocalization analyses

Colocalization was assessed with the coloc R package (v5.2.3) using the Bayesian single-causal-variant approximation^140^ (coloc.abf). At each locus, effect alleles were harmonized between the two datasets, QTL effect sizes were sign-flipped where the effect allele was reversed, and strand-ambiguous palindromic variants (A/T and C/G) were removed. Loci were retained only if at least 50 variants overlapped after harmonization. Signal pairs with a posterior probability of a shared region PP.H4 > 0.7 were considered colocalized.

eQTL-GWAS colocalization. eQTLs were tested against 13 cardiometabolic GWAS spanning adiposity and fat-distribution traits (BMI; WHR, abdominal, and gluteofemoral fat adjusted for BMI), glycemic and insulin traits (fasting insulin, HbA1c, and HOMA-IR, ISI, and IFC each adjusted for BMI), type 2 diabetes, and lipid traits (HDL, LDL, triglycerides). We applied a GWAS-centric analysis: for each GWAS lead variant, candidate eGenes were defined as those whose cis window (±1 Mb around the gene midpoint) spanned the lead variant. We considered the GWAS lead variant if the individual study reported so; otherwise, we identified genome-wide significant (P < 5e-8) signals in 1Mb windows. GWAS and eQTL summary statistics were then extracted within ±500 kb of the GWAS lead variant and intersected by genomic position. coloc.abf was run per candidate eGene. Both basal and context-dependent eQTLs were evaluated. For basal eQTLs, the eGenes identified under basal conditions in each cell state were used directly. For context-dependent eQTLs, the static effect and each context-specific genotype-by-context (G×C) interaction effect (free fatty acid [FFA], low glucose, hypoxia) were tested. The AMSC quiescent state was tested for the static effect only.

caQTL-GWAS colocalization. Using the same GWAS-centric ±500 kb window, cis-caQTLs were tested against each GWAS. Peak significance was determined from the TensorQTL cis-permutation pass (10 kb window; permutation FDR < 0.05); for colocalization, the nominal cis summary statistics of these peaks (mapped with a ±1 Mb cis window) were used, restricted to the ±500 kb region around the GWAS lead variant. Each significant peak overlapping a locus was tested individually against the GWAS with coloc.abf.

eQTL-caQTL colocalization. To link genetically regulated expression with chromatin accessibility within each cell type, eQTLs and caQTLs were colocalized in an eGene-centric design. For each eGene, the eQTL lead variant was defined as the most significant cis-variant, a ±1 Mb window was taken around it, and all significant caQTL peaks (permutation FDR < 0.05) overlapping the window were tested. coloc.abf was run separately for each overlapping peak.

### Summary-data-based Mendelian randomization

Summary-data-based Mendelian randomization^141^ (SMR) with the accompanying heterogeneity in dependent instruments (HEIDI) test (SMR v1.4.0) was used to identify pleiotropic associations, taking the top associated cis-QTL as the instrumental variable and 1000 Genomes European-ancestry samples (GRCh38) as the LD reference. Within each cell state we tested three types of association: variant-eGene-trait (cis-eQTL against GWAS), using the CASTIE static (main genotype) effect for the basal/quiescent analysis and the genotype-by-context (G×C) interaction effect for each stimulatory context (FFA, low glucose, hypoxia), tested against the 13 cardiometabolic GWAS described above (Colocalization analyses); variant-caPeak-trait (cis-caQTL against the same GWAS), using the nominal cis-caQTL summary statistics (±1 Mb window) of chromatin peaks significant in the TensorQTL cis-permutation analysis (10 kb window; permutation FDR < 0.05); and variant-caPeak-eGene, linking chromatin accessibility to gene expression with the cis-caQTL as the exposure and the cis-eQTL of each eGene as the outcome. QTL summary statistics were formatted as BESD files, and the instrument-selection threshold was P_QTL < 1 × 10⁻⁵ (relaxed to 1 × 10⁻⁴ for the perturbation conditions in the variant-caPeak-eGene analysis). Within each cell state, P_SMR was corrected by the Bonferroni method, associations with P_SMR < 0.05/(number of tests) and P_HEIDI ≥ 0.01 were considered significant, and for the QTL-GWAS tests significant associations with P_GWAS > 1 × 10⁻⁵ were additionally excluded. We did not use the HEIDI test for G × C eQTLs because genotype LD alone does not capture uncertainty and covariance in G × C statistics.

### Transcription factor binding analysis

To investigate whether sequence-level mechanisms could account for the low-glucose-specific eQTL effect at the *NCOR2* locus, we performed transcription factor (TF) binding analysis using motifbreakR^142^ (v2.10.2) against the HOCOMOCO v11 human position-weight-matrix (PWM) collection^143^, retrieved via the MotifDb Bioconductor package. Four candidate variants at the *NCOR2* locus colocalized between the ISIadjBMI GWAS, eQTL, and caQTL signals were tested: the GWAS lead SNP rs1906937 (chr12:124,054,990, C>A), the eQTL lead SNP (chr12:123,991,393, A>G; adipocyte under low glucose), and two caQTL lead SNPs (chr12:123,955,563, C>A and chr12:123,949,358, C>T). Motif matches were scored using motifbreakR’s information-content method (method = “ic”) with a p-value threshold of 1 × 10⁻³, a uniform nucleotide background (A = C = G = T = 0.25), and filterp = TRUE. For each hit, the change in predicted binding score between alleles (alleleDiff = scoreAlt − scoreRef) was used to classify the variant as motif-creating (alleleDiff > 0) or motif-disrupting (alleleDiff < 0), and motifbreakR’s built-in effect classification was used to separate “strong” from “weak” hits. Sequence logos for the heterodimer components were generated from the corresponding HOCOMOCO PWMs using the ggseqlogo R package (method = “bits”).

### snATAC-seq peaks and variant annotation

Annotation of the genomic location of snATAC-seq peaks, eQTLs, caQTLs and cpQTLs was performed using the ChIPseeker package^144^ (v1.45.0).

### CRISPR interference (CRISPRi) validation of effector genes

The targeted CRISPRi experiments used a human white adipose tissue (hWAT) preadipocyte line expressing an AAVS1 safe harbor-integrated doxycycline-inducible dCas9-KRAB (idCas9-KRAB). The parental hWAT preadipocytes were isolated and characterized previously^145^ and the idCas9-KRAB line was generated from these cells as described^146^. Cells were maintained in maintenance medium (DMEM-High glucose, 10% FBS) at 37 °C and 5% CO₂ and passaged before reaching confluence.

For differentiation, cells were plated at 2 × 10⁵ cells per well in 6-well plates, and doxycycline (0.1 µg/mL) was added the following day to induce dCas9-KRAB for 2 days before the start of differentiation and maintained throughout. Differentiation was then induced by switching to differentiation medium, maintenance medium supplemented with 33 µM biotin, 17 µM pantothenate, 0.5 µM insulin, 0.1 µM dexamethasone, 2 nM triiodothyronine (T3), 500 µM IBMX, 30 µM indomethacin, and 1 µM rosiglitazone. Media with doxycycline was refreshed every 2-3 days for 14 days.

sgRNAs targeting the transcription start sites of the target genes *CITED2* and *TCF7L2*, together with non-targeting controls (guide sequences see **Supplementary Notes**), were designed using CRISPick (Broad Institute; https://portals.broadinstitute.org/gppx/crispick/public). The sgRNAs were synthesized as an oligonucleotide pool, amplified by PCR, and cloned as a pool into the lentiviral CROP-seq-Opti vector (Addgene Plasmid #106280), which was cut with BsmBI, using Gibson assembly. The assembled pool was introduced into electrocompetent bacteria by electroporation, and plasmid DNA was purified with an endotoxin-free prep. Lentivirus was then produced in HEK293T cells by transfection of the pooled sgRNA library plasmid with the packaging plasmid psPAX2 and a VSVG envelope plasmid. Viral supernatant was collected ∼72 h post-transfection, passed through a 0.45-µm filter, and stored at −80 °C. idCas9-KRAB hWAT cells were transduced with the pooled sgRNA lentivirus in the presence of 4ug/mL polybrene at low MOI, selected with puromycin for 72 hours, expanded, and then induced with doxycycline and differentiated as described above.

At day 14 of differentiation, cells were lifted to a single-cell suspension, filtered, and counted, and viability was assessed with trypan blue. Single-cell suspensions were loaded on the 10x Genomics Chromium platform using the Chromium Next GEM Single Cell 3′ v4 kit according to the user’s guide. In addition to the gene-expression library, sgRNA identities were recovered from the amplified cDNA by targeted (“dial-out”) PCR to generate a paired guide library. Both libraries were sequenced on an Illumina NovaSeq X platform, and reads were processed with Cell Ranger multi (v9.0.1) to generate single-cell gene-expression matrices and per-cell sgRNA assignments.

Differential expression analysis was performed using the improved negative binomial test implemented in SCEPTRE^147^ (v0.9.1). Count and guide matrices were imported directly from the CellRanger “sample_filtered_feature_bc_matrix” outputs with “import_data_from_cellranger”, treating the experiment as low multiplicity of infection (MOI). Guides were annotated as either TSS-targeting or non-targeting controls. gRNAs were assigned to cells using the thresholding method. Cell-wise quality control removed cells with a mitochondrial UMI fraction above 0.15 and cells whose number of expressed genes fell outside the 10th-99th percentile range, in addition to SCEPTRE’s default low-MOI removal of cells containing zero or more than one gRNA (MOI after guide assignment in SCEPTRE = 0.81). Pairwise quality control retained target-gene pairs with at least seven cells expressing the response gene in both the treatment and control groups. Tests were adjusted for the default cell-level covariates computed by SCEPTRE together with sample lane information. Prior to the discovery analysis, a calibration check was run in which negative-control pairs were constructed from non-targeting gRNAs to confirm that the test produced approximately uniform p-values and controlled false discoveries; the discovery analysis was then run with the same parameters. Significance was determined by Benjamini-Hochberg FDR correction at FDR < 0.05, and effect sizes are reported as log2 fold changes in expression.

For the *cis* analysis, candidate target-gene pairs were constructed with “construct_cis_pairs” using a distance threshold of 5 Mb, and a left-tailed test was used. For the *trans* analysis, all trans target-gene pairs were constructed with “construct_trans_pairs”, excluding positive-control pairs, and a two-sided test was used. To reduce computational burden in the trans setting, the response set was restricted to genes with nonzero expression in at least 50 cells across all libraries prior to testing. Genes identified as differentially expressed by SCEPTRE were subjected to gene set enrichment analysis (GSEA) as described above.

## Data availability

Paired single nucleus ATAC-seq and RNA-seq raw sequencing data as well as the processed seurat object (RNA modality) and snapATAC2 Anndata object (ATAC modality) will be available and deposited at GEO upon publication.

## Code availability

GitHub for CASTIE: https://github.com/ZhouLabGenetics/CASTIE. Processed R or Python objects and Code will be available from GitHub upon publication. Any additional information required to reanalyse the data reported in this paper is available from the lead contact upon request.

## Acknowledgements

This work was supported by the Novo Nordisk Foundation (NNF21SA0072102, NNF20OC0059796), and NIH grants RC2DK116691, RC2 DK144819-01, UM1DK126185, and the Boston Area Nutrition Obesity Research Center sponsored by NIH P30 DK040561. M.C. is further supported by the Weissman Family MGH Research Scholar Award. We thank our surgical colleagues at BIDMC for their help procuring samples.

## Author contributions

Conceptualization, M.C, Y.H, J.P.S., T.R.J.; Funding acquisition, M.C., B.N., R.A., W.Z.; Methodology, Y.H., J.P.S., Y.C.L., T.R.J., W.Z. and M.C.; Investigation, Y.H., J.P.S., S.D., B.M., N.N., P.K., B.S., S.S.P. and M.C.; Data curation, Y.H., J.P.S., Y.C.L., M.M., T.R.J., H.D. and W.Z.; Formal analysis, Y.H., J.P.S., Y.C.L., M.M., W.L.Q., T.R.J., H.D., A.G.M. and K.H.; Validation, R.M.G., T.M.B.; Visualization, Y.H., J.P.S., Y.C.L., A.G.M., T.R.J., H.D., and K.H.; Writing - Original draft, Y.H., J.P.S., Y.C.L., and M.C.; Writing - review & editing, Y.H, J.P.S., S.D., Y.C.L., B.M., M.M., N.N., P.K., B.S., S.V.S.P., R.M.G., T.M.B., A.G.M., K.H., W.L.Q., R.A., T.R.J., H.D., C.G., B.N., W.Z. and M.C.; Software, Y.H., Y.C.L., W.Z., T.R.J. and H.D.; Supervision, M.C., W.Z., T.R.J., H.D., C.G., and B.N.

## Supplementary information

|  |
| --- |
| Supplementary Table 1. Donor characteristics |
| Supplementary Table 2. snMultiome sequencing QC metrics |
| Supplementary Table 3. Transcriptomic signatures of cNMF clusters |
| Supplementary Table 4. Transcriptomic signature enrichment pathway analysis of cNMF clusters |
| Supplementary Table 5. Cell state marker genes |
| Supplementary Table 6. Cell state marker snATAC-seq peaks |
| Supplementary Table 7. Cell state marker TF motifs |
| Supplementary Table 8. Cell state transcriptomic signature enrichment pathway analysis |
| Supplementary Table 9. DEGs between stimulatory groups |
| Supplementary Table 10. DAPs between stimulatory groups |
| Supplementary Table 11. Enrichment pathway analysis of DEGs |
| Supplementary Table 12. scE2G peak-gene links |
| Supplementary Table 13. DAPs TF motifs enrichment analysis |
| Supplementary Table 14. DAPs TF motifs PPI clusters |
| Supplementary Table 15. Statistics of the scDRS group-level analysis across cell state |
| Supplementary Table 16. Enrichment pathway analysis of DEGs between high-scDRS and low-scDRS SWAT cells |
| Supplementary Table 17. Statistics of the scDRS group-level analysis across cell state and stimuli |
| Supplementary Table 18. Statistics of the scDRS vs expression PC correlation |
| Supplementary Table 19. Pseudobulk eQTLs |
| Supplementary Table 20. Pseudobulk caQTLs |
| Supplementary Table 21. Single-cell context eQTLs |
| Supplementary Table 22. Genomic distributions of single-cell context eQTLs |
| Supplementary Table 23. Colocalization between eQTLs and caQTLs |
| Supplementary Table 24. Colocalization between single-cell context eQTLs and GWAS traits |
| Supplementary Table 25. Colocalization between caQTLs and GWAS traits |
| Supplementary Table 26. Binding TFs at NCOR2 locus |
| Supplementary Table 27. SMR between single-cell context eQTLs and GWAS traits |
| Supplementary Table 28. SMR between caQTLs and GWAS traits |
| Supplementary Table 29. SMR between caQTLs and eQTLs |
| Supplementary Table 30. SMR 3-way regulatory cascade |
| Supplementary Table 31. Adipose biological processes after CITED2 knockdown |
| Supplementary Table 32. cpQTLs |
| Supplementary Table 33. Adipose biological processes after TCF7L2 knockdown |

## Extended data Figures

**Extended Data Fig. 1 (related to Figure 1):**
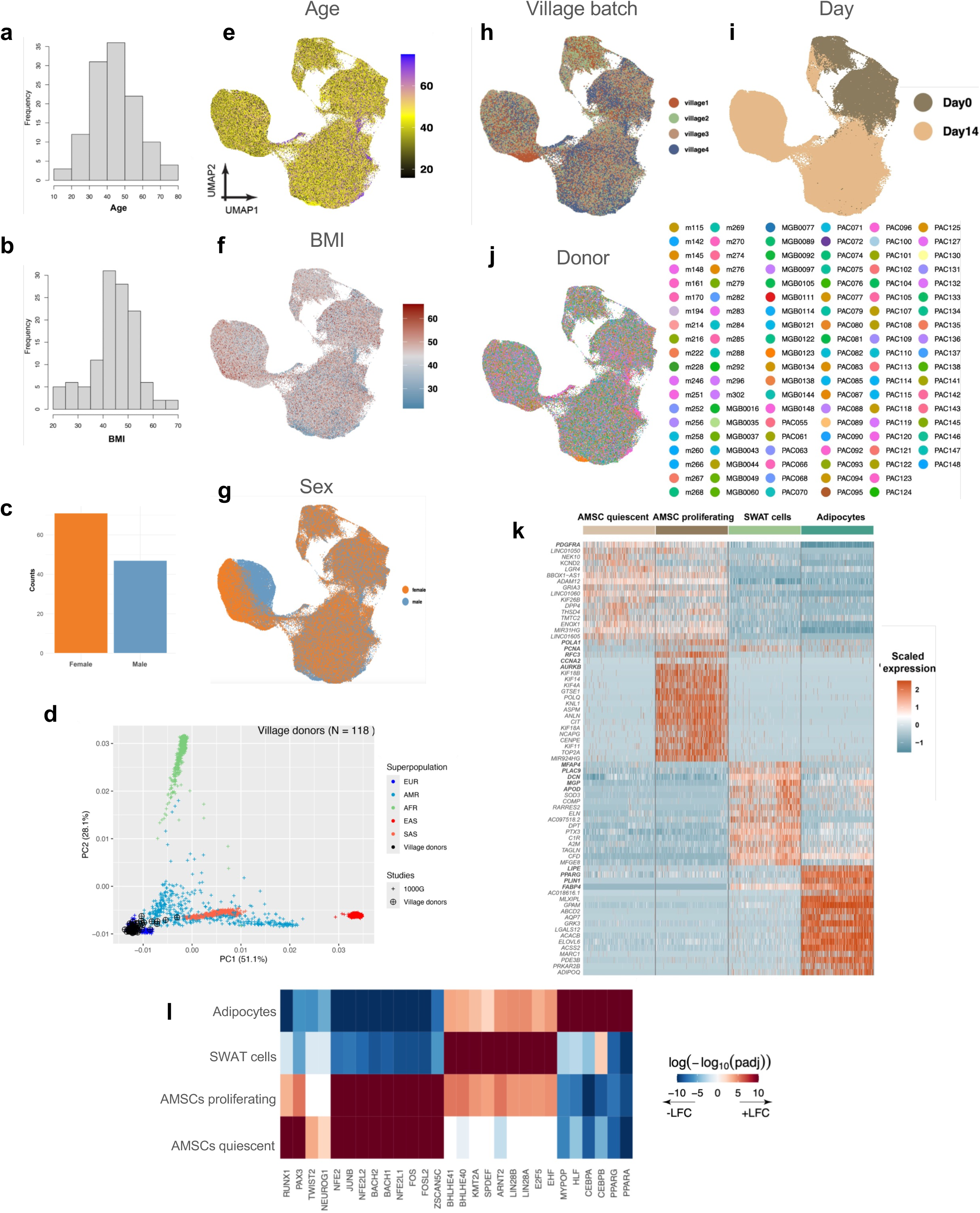
A human adipocyte snMultiome map under metabolic disease-relevant stimuli. **a-c,** Histograms of age, BMI and sex across all donors. **d,** Principal-component analysis of genotypes for the 118 village donors (black) projected onto 1000 Genomes reference populations (EUR, AMR, AFR, EAS, SAS), showing the ancestry composition of the cohort. **e-j,** UAMPs colored by age, BMI, sex, village batch, day and donor ID after batch correction. **k,** Heatmap of transcriptomic signatures of each cell state under basal condition (min.pct =0.5, log2FC >0.75; canonical marker genes are highlighted in bold). **l,** Heatmap of transcription factor (TF) motifs enriched in cell state marker peaks. Color indicates the signed enrichment, −log₁₀(adj. P) × sign of log fold change (+LFC / −LFC); enriched motifs are shown at adj. P < 0.05.

**Extended Data Fig. 2 (related to Figure 1):**
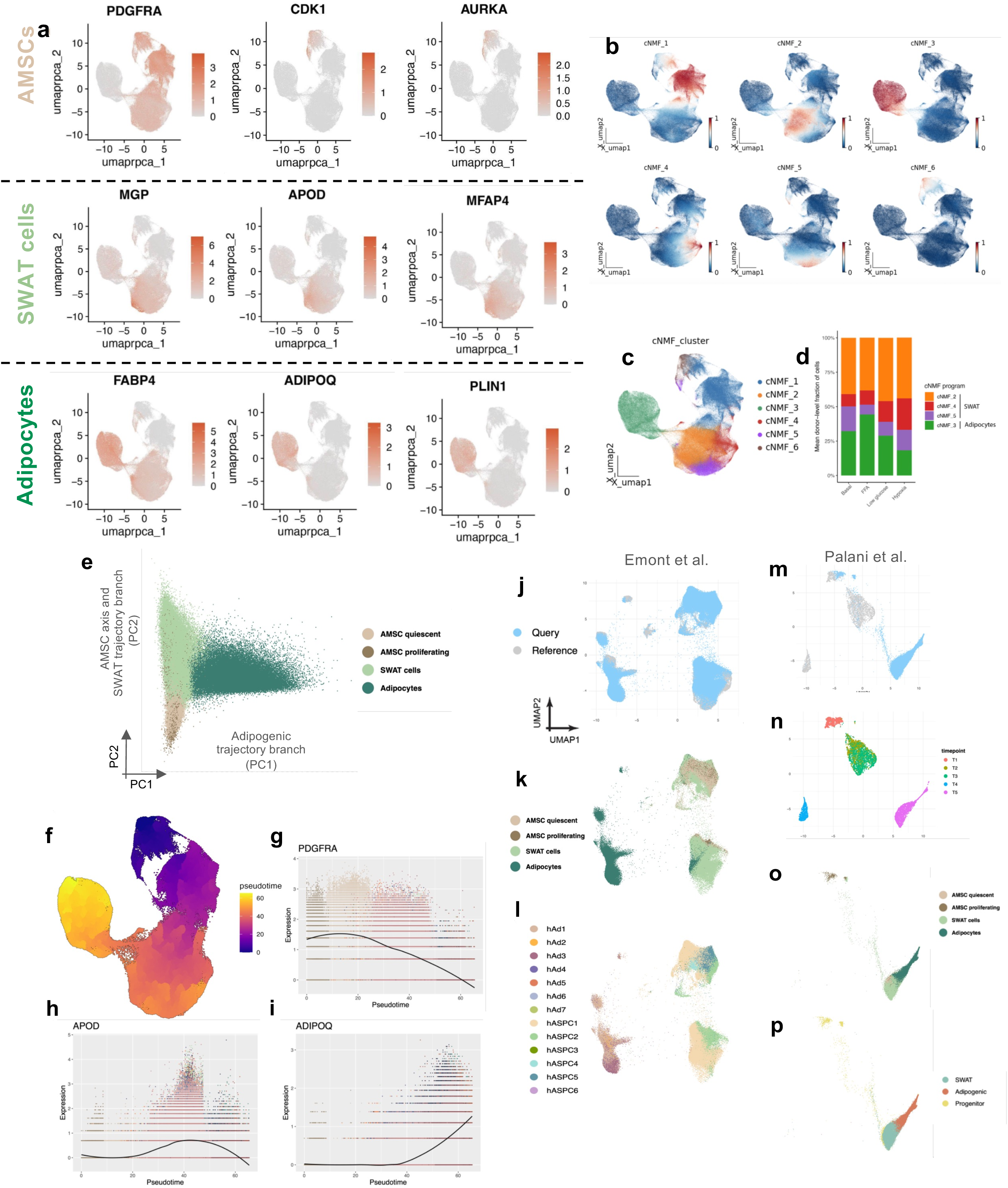
A human adipocyte snMultiome map under metabolic disease-relevant stimuli. **a,** UMAPs of canonical cell-state marker gene expression: AMSC (*PDGFRA*, *CDK1*, *AURKA*), SWAT cells (*MGP*, *APOD*, *MFAP4*), and adipocytes (*FABP4*, *ADIPOQ*, *PLIN1*). Color indicates normalized expression. **b,** UMAPs of the six consensus non-negative matrix factorization (cNMF) gene-expression programs. Color indicates the scaled program scores. **c,** UMAP colored by the dominant cNMF program assigned to each cell. **d,** Mean per-donor fraction of cells assigned to each cNMF program across stimulus conditions (basal, FFA, low glucose, hypoxia) for SWAT cells and adipocytes. **e,** PCA analysis shows that PC1 captures the adipogenic trajectory, while PC2 reflects the AMSC axis and SWAT cell trajectory. **f,** UMAP colored by pseudotime along the differentiation trajectory. **g-i,** Expression of marker genes along pseudotime: **(g)** *PDGFRA* (AMSCs), **(h)** *APOD* (SWAT cells), and **(i)** *ADIPOQ* (adipocytes), illustrating the progression from progenitor to differentiated states. **j-p,** Co-embedding of our dataset with *in vivo* adipose tissue and *in vitro* adipocyte differentiation sn/scRNA-seq reference datasets. **(j)** and **(m)** co-embedded UMAP embedding of the query dataset (our data in blue) and *in vivo/in vitro* dataset (Emont et al. and Palani et al. in gray). **(k)** and **(o)** UMAP colored by cell state annotation from our dataset. **n,** UMAP colored by five experimental time points from the *in vitro* reference data. **(l)** and **(p)** UMAP colored by transferred annotations from the *in vivo and in vitro* reference dataset.

**Extended Data Fig. 3 (related to Figure 2):**
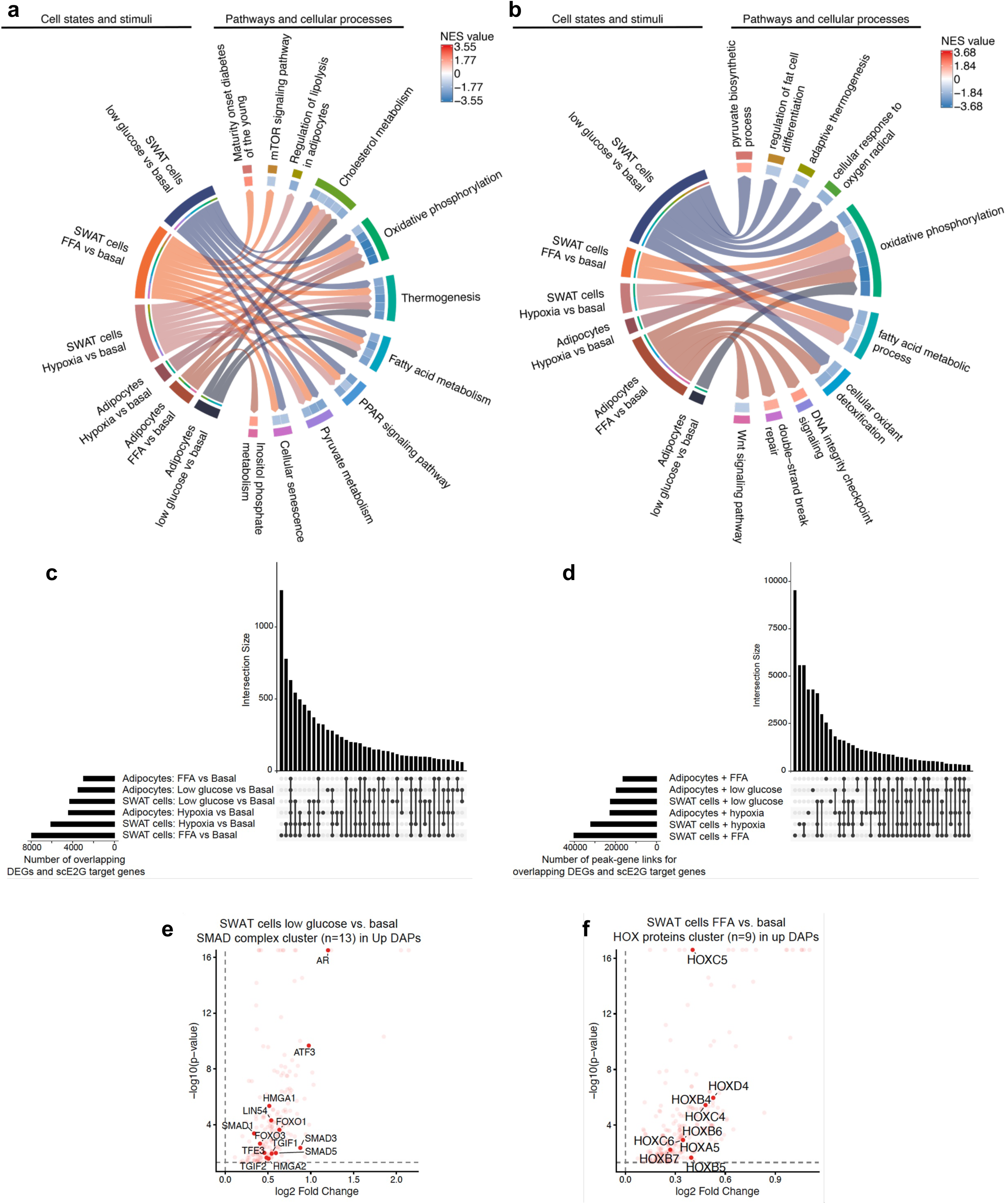
Functional enrichment analysis of DEGs. **a,b**, Representative metabolic disease-associated cellular programs from functional enrichment analysis from (**a**) KEGG and (**b**) GOBP databases (NES: normalized enrichment score) of DEGs after 14 days of exposure to FFA, low glucose, or hypoxia compared to the basal state. **c,** Number of peak-gene links per cell state and stimulus predicted by scE2G analysis. **d,e**, Predicted protein-protein interaction networks of the (**d**) nuclear receptor cluster under low glucose and (**e**) the AP-1 cluster under FFA in adipocyte chromatin regions gaining accessibility (Up DAPs).

**Extended Data Fig. 4 (related to Figure 3):**
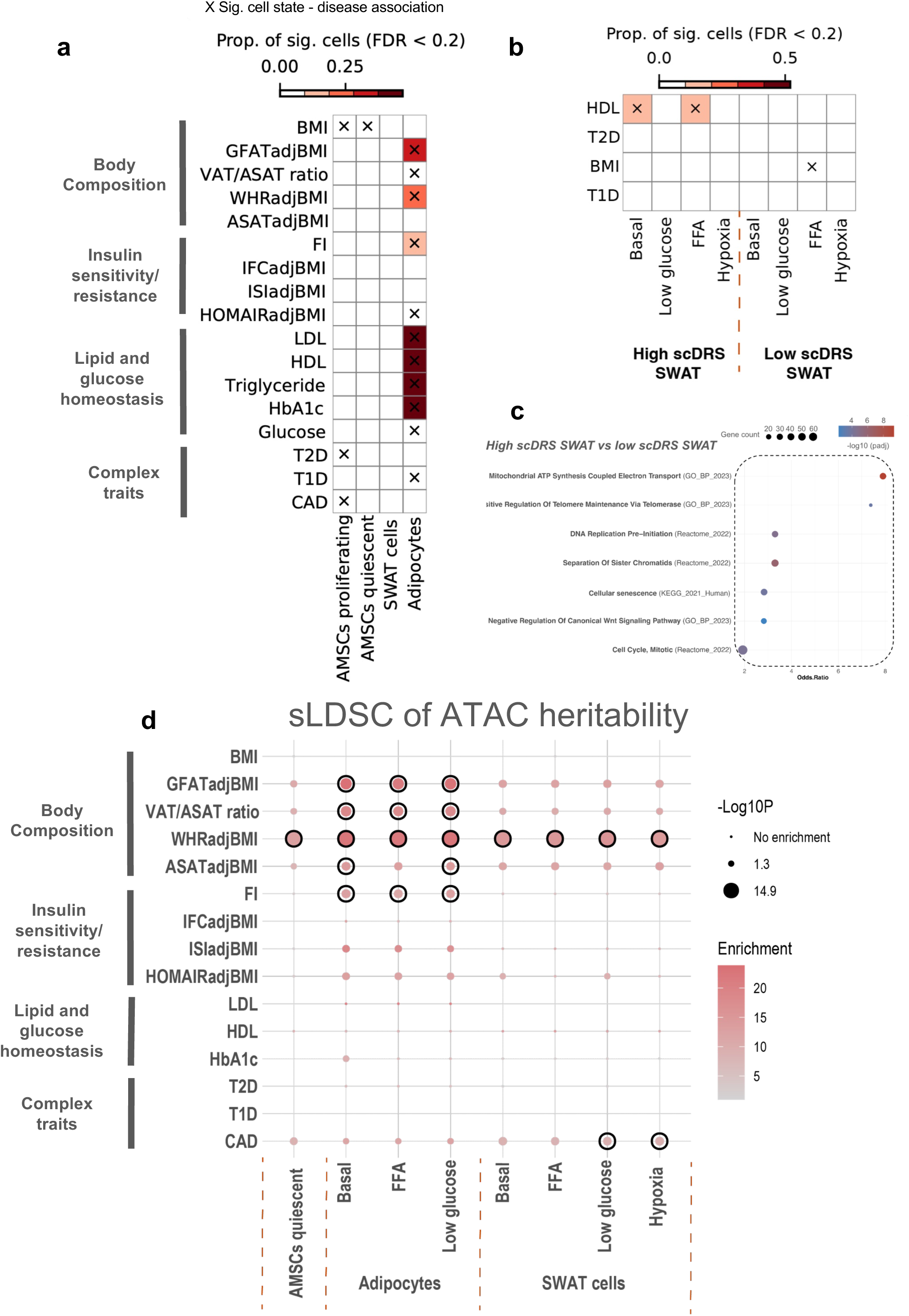
Disease heritability mapping. **a,b** Heatmaps showing heritability enrichment between **(a)** major cell states, **(b)** sub populations of SWAT cells under stimuli (columns) and GWAS traits (rows). Heatmap color represents the proportion of significantly associated nuclei (FDR < 0.2 across all nuclei for a given trait). Cross indicate significant cell-state-disease associations (BH adjusted FDR < 0.05 across all pairs of cell states and traits; MC test). **c,** Dotplot showing the significantly enriched pathways (padj <0.05) for DEGs between high-scDRS and low-scDRS SWAT populations. **d,** sLDSC heritability enrichment analysis based on ATAC modality between a given cell state under stimuli and GWAS trait. Circle with outlines indicates significant enrichment after bonferroni adjustment (q <0.05). The smallest dots represent non-significant enrichment (P >=0.05). Dot color reflects the magnitude of heritability enrichment.

**Extended Data Fig. 5 (related to Figure 4):**
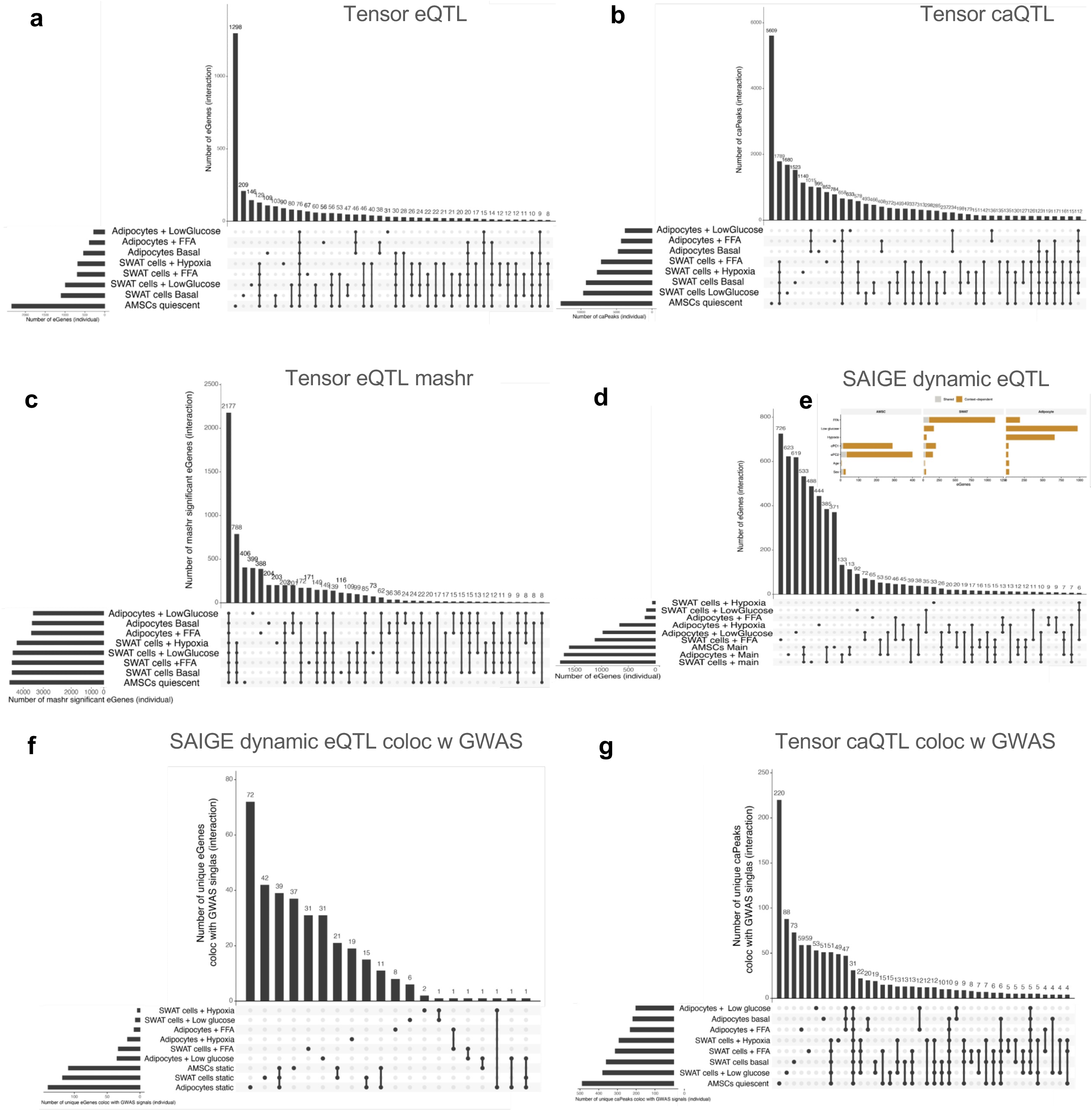
e/caQTL mapping. **a,** UpSet plot showing the number of eGenes identified in each cell state under stimuli using pseudobulk QTL mapping (TensorQTL, FDR <0.05). **b,** UpSet plot showing the number of caPeaks identified in each cell state under stimuli using pseudobulk QTL mapping (TensorQTL, FDR <0.05). **c,** UpSet plot showing the number of eGenes identified in each cell state under stimuli using pseudobulk QTL after multivariate adaptive shrinkage (mashr, lfsr <0.05). **d,** UpSet plot showing the number of eGenes identified in each cell state under stimuli using single-cell context QTL mapping (CASTIE, FDR <0.05). **e,** Number of context-dependent eGenes identified per context (FFA, low glucose, hypoxia, expression PCs ePC1-ePC2, age, sex) in each cell state. Bars are colored by category (gold, context-dependent; grey, shared). **f,g,** UpSet plots showing the number of unique **(e)** eGenes and **(f)** caPeaks colocalize with GWAS trait signals across cell state and stimuli.

**Extended Data Fig. 6 (related to Figure 4):**
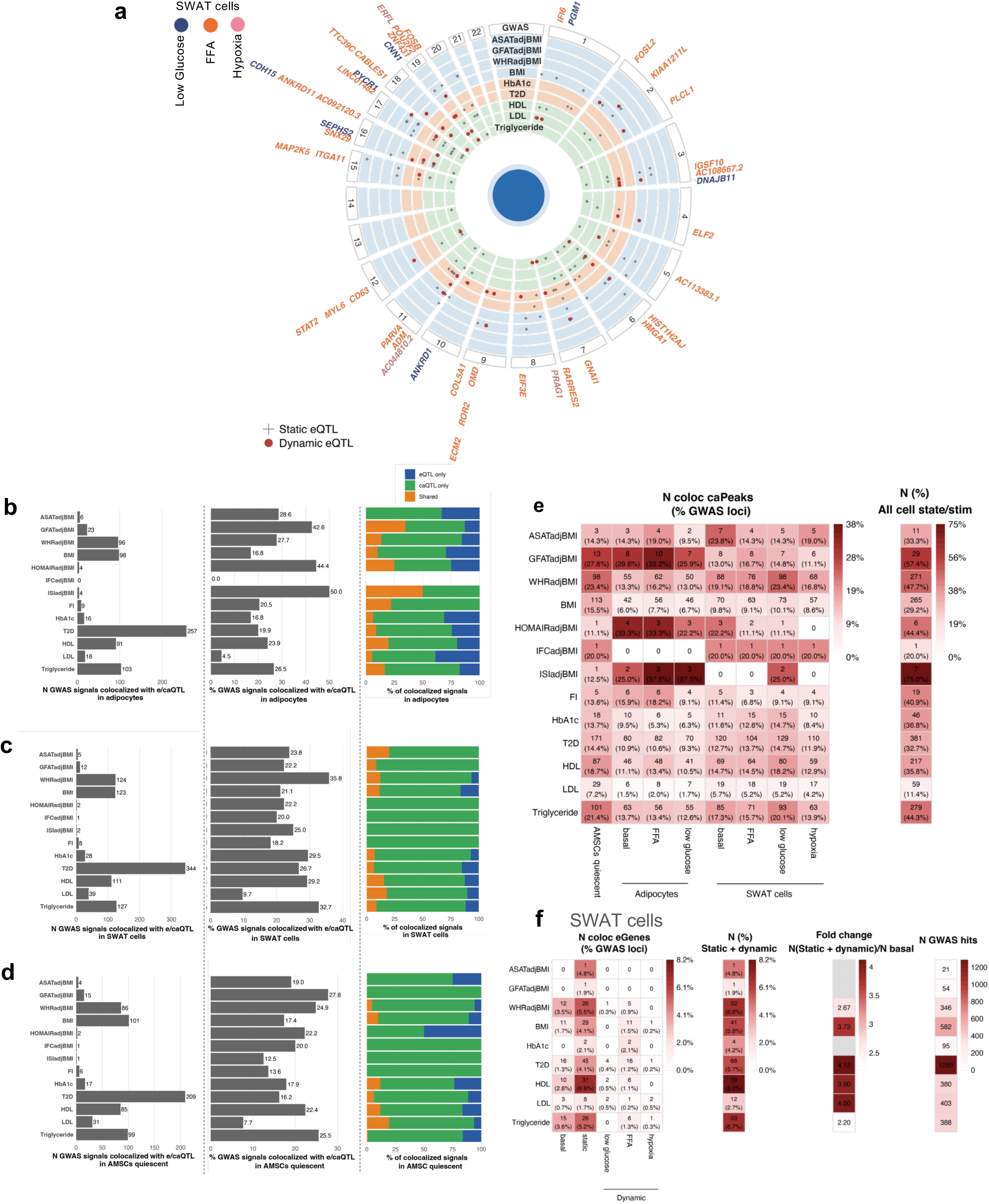
e/caQTL mapping. **a**, Circular plot of eQTL-GWAS colocalizations in SWAT cells across stimuli. Genes are positioned by genomic location and colored by stimulus, concentric tracks correspond to GWAS traits, and points mark significant colocalizations (red dot = dynamic eGenes, grey cross = static eGenes). **b-d,** Number (left), percentage (middle), and colocalization pattern (right) of GWAS signals that colocalize (PP.H4>0.7) with eQTL and/or caQTL signals across stimuli and traits in **(b)** adipocytes, **(c)** SWAT cells and **(d)** AMSCs. The right panel shows the proportion of colocalized GWAS signals explained by caQTL only, eQTL only or both. **e,** Per-trait caQTL colocalization summary across cell states and stimuli. Numbers of colocalized caPeaks (and % of GWAS loci explained) are indicated in the heatmap. Color indicates the % GWAS loci can be explained by caQTL signals. **f,** Per-trait eQTL colocalization summary in SWAT cells. From left to right: number of colocalized eGenes at basal condition, under static and dynamic effects (and % of GWAS loci explained); number (%) of GWAS signals with static + dynamic eQTL explain; fold change in colocalized signals from static + dynamic versus basal-only models; and number of total GWAS hits. Color indicates the % GWAS loci can be explained by eQTL signals.

**Extended Data Fig. 7 (related to Figure 5):**
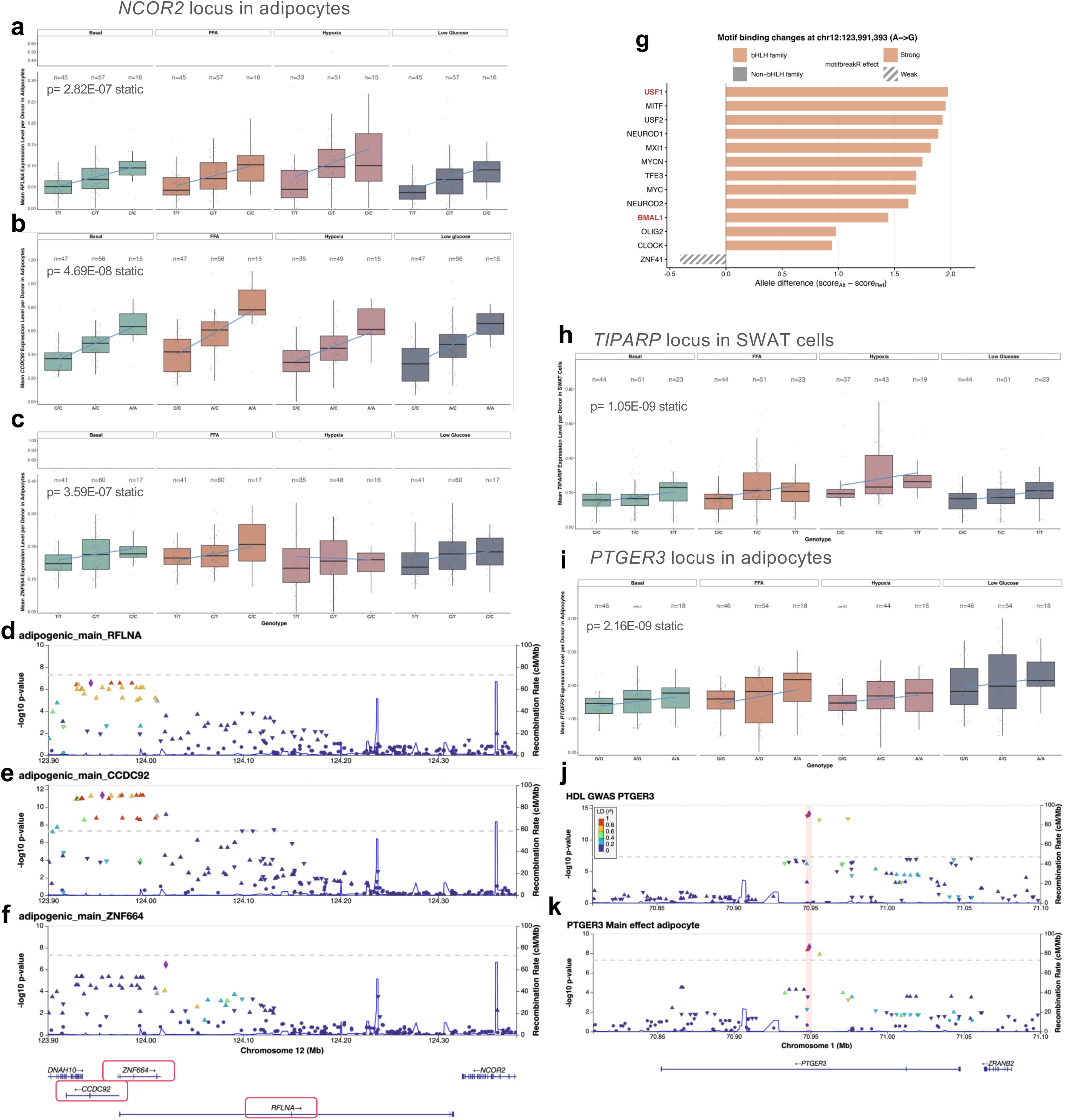
Additional e/caQTL colocalization examples at the *NCOR2*, *TIPARP*, and *PTGER3* loci. **a-c**, Expression by genotype in adipocytes for neighboring eGenes at the *NCOR2* locus: **(a)** *RFLNA*, **(b)** *CCDC92*, and **(c)** *ZNF664*, each showing a static eQTL across stimuli. **d-f,** Locus plots of adipocyte static-effect eQTLs at the *NCOR2* locus for **(d)** *RFLNA*, **(e)** *CCDC92*, and **(f)** *ZNF664*. Each panel shows −log₁₀(P) for the eQTL summary statistics (left y-axis) and the recombination rate (right y-axis); points are colored by LD (r²) with the lead variant. **g**, Motif-binding changes at chr12:123,991,393 (A→G), ranked by allele difference (scoreAlt − scoreRef). Bars indicate transcription factors with altered predicted binding; the alt allele **(G)** strengthens binding of bHLH/bHLH-Zip factors. **h,** *TIPARP* expression by genotype across stimuli in SWAT cells, showing a static eQTL. **i,** *PTGER3* expression by genotype across stimuli in adipocytes, showing a static eQTL. **j,k,** Locus plots showing colocalization of the *PTGER3* locus with **(j)** HDL GWAS and **(k)** the static-effect *PTGER3* eQTL in adipocytes. −log₁₀(P) for the GWAS or eQTL summary statistics (left y-axis) and recombination rate (right y-axis); points are colored by LD (r²) with the lead variant rs500647 (highlighted).

**Extended Data Fig. 8 (related to Figure 6):**
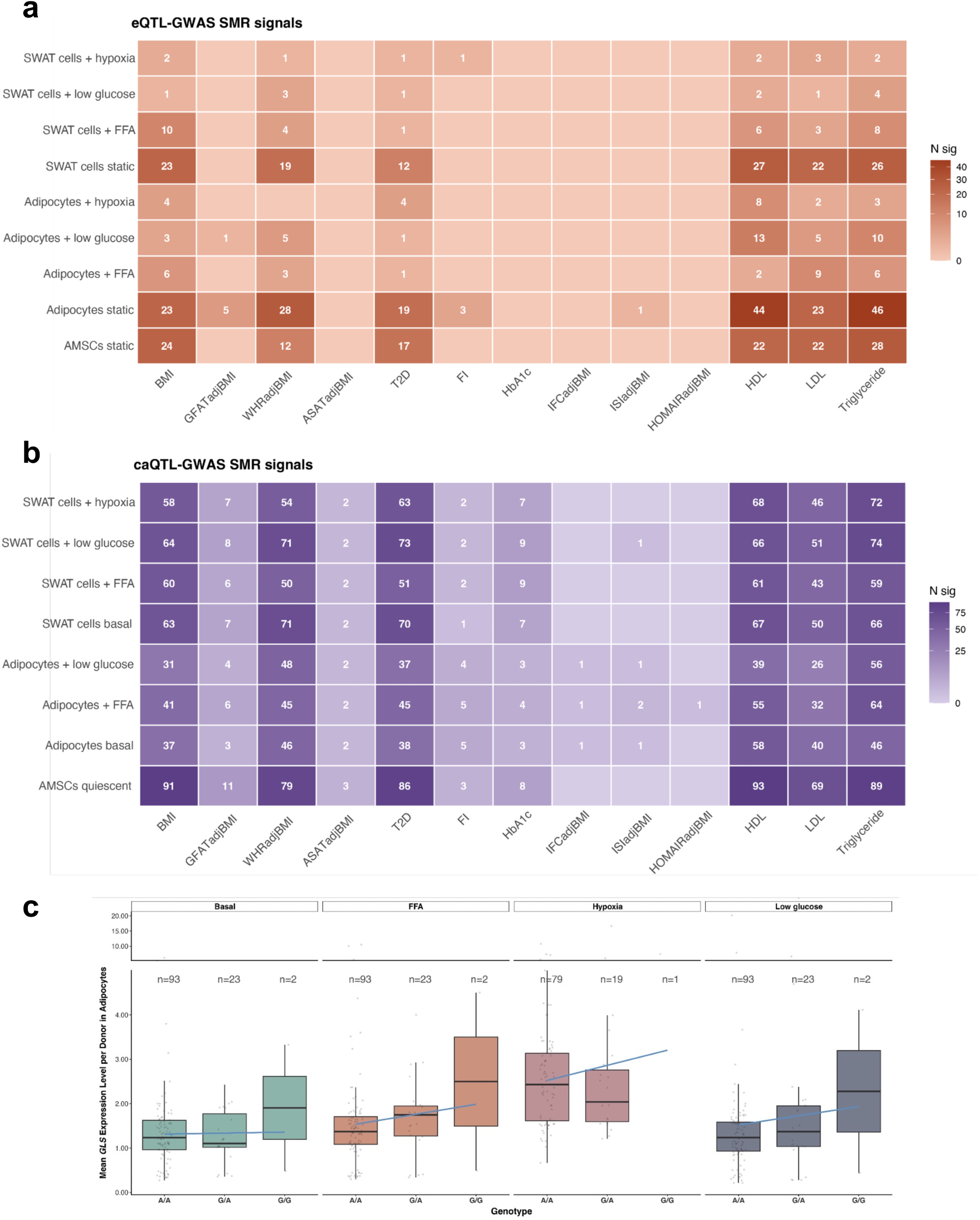
SMR analysis. **a,** Number of significant eQTL-SMR signals per GxC pattern (rows) and GWAS trait (columns). Color intensity indicates the number of significant eGene-trait associations (N sig). **b,** Number of significant caQTL-SMR signals per cell state × stimulus condition (rows) and GWAS trait (columns). Color intensity indicates the number of significant caPeak-trait associations (N sig). **c,** *GLS* expression by genotype across stimuli in adipocytes, showing a FFA-specific eQTL (lead eQTL variant rs11696087).

**Extended Data Fig. 9 (related to Figure 7):**
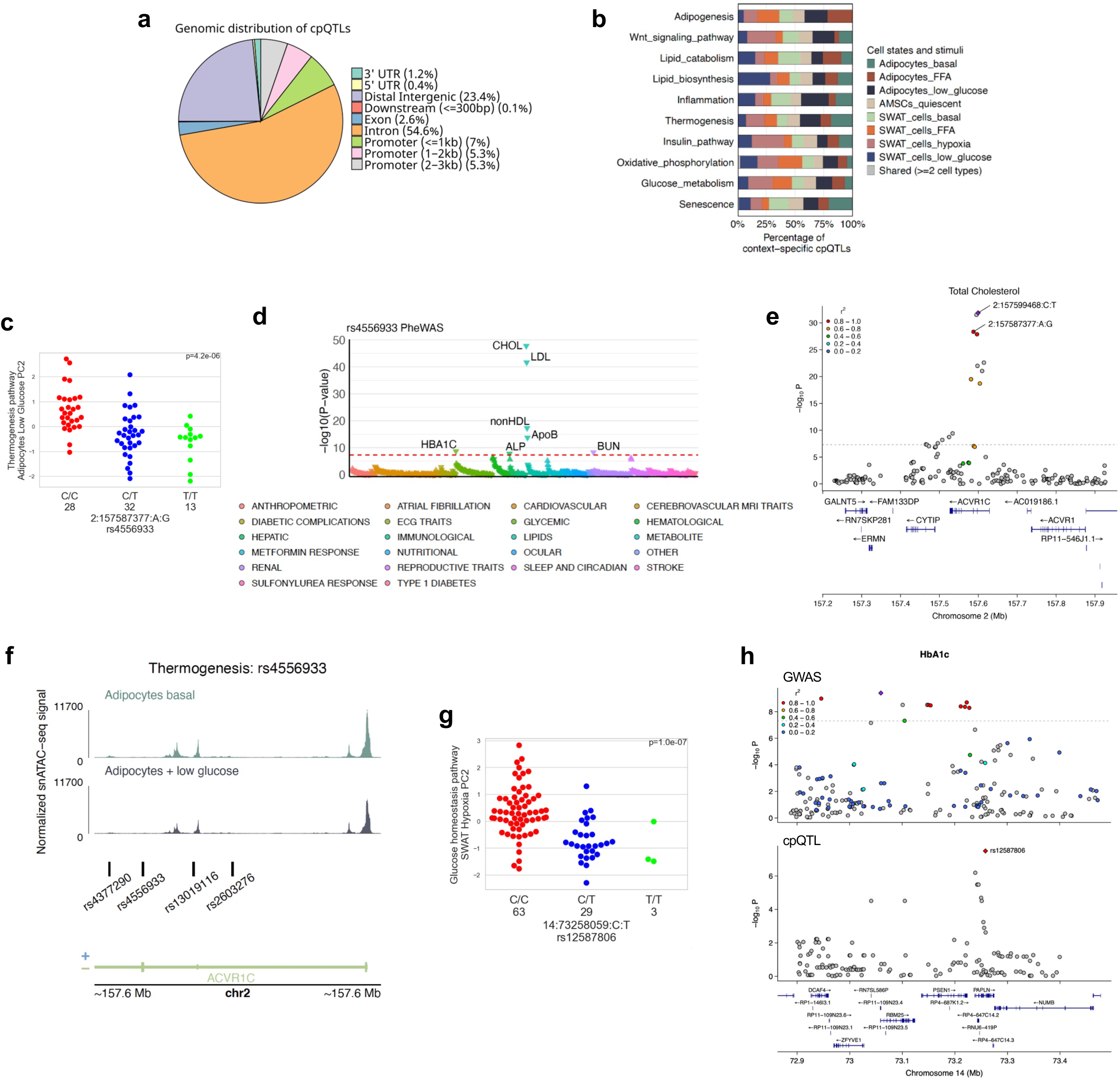
Cellular program QTLs analysis. **a**, Pie chart showing the genomic distribution of all cpQTLs (p-value < 5 × 10^-6^). **b**, Proportion of context-specific cpQTLs across different cell states and stimuli within each cellular program. **c**, Thermogenesis program expression in adipocytes under low glucose across rs4556933 alleles. **d**, PheWAS associations for the cpQTL rs4556933. **e**, **f**, *ACVR1C* locus with (**d**) total cholesterol GWAS association highlighting the lead SNP and cpQTL rs4556933, and (**e**) genomic tracks of normalized snATAC-seq signal in adipocytes under basal or low glucose conditions, with cpQTLs rsIDs highlighted in the bottom panel. **g,** Glucose metabolism program expression in SWAT cells exposed to hypoxia across rs12587806 alleles. **h**, HbA1c GWAS (top panel) and rs12587806 cpQTL (bottom panel) locus plots, highlighting significant colocalization.

