## Supplementary information for "Metabolic disease-relevant stimuli unmask context-dependent genetic regulation of cardiometabolic loci in human adipocytes"

### Description of Supplementary information


| | |
|---|---|
| Supplementary notes | Page 1 - 4 |
| Supplementary figures | Page 5 - 6 |

### Supplementary notes

#### Extended information for snMultiome data processing

We first processed each sample independently to assess data quality for both snRNA-seq and snATAC-seq modalities. Initially, a village batch effect was observed when all samples were merged without batch correction **(Supplementary Fig. S1a)**. After applying RPCA batch correction on RNA modality, we found that the village-specific batch effects were effectively harmonized across the dataset **(Extended Data Fig. 1g)**. We observed the similar batch correction performance using scVI (**Supplementary Fig. S1b**) (corrected for village batch and conditioned by 10x chip loading batch, with 92.5% of proliferating AMSCs, 92.73% of quiescent AMSCs, 90.24% of SWAT cells, and 97.43% of adipocytes overlapping between the two methods). A subcutaneous immortalized adipose tissue derived mesenchymal stem cell (iAMSC; donor m130) was included in each village as an internal control to evaluate the efficiency of batch correction. Notably, iAMSCs from donor m130 consistently clustered together post-correction **(Supplementary Fig. S1c)**, while the overall structure of the dataset was maintained, indicating successful removal of batch effects without overcorrection.

#### Extended information about the AMSC proliferating state

Proliferating AMSCs undergoing cell cycling were predominantly derived from villages 1 and 2 (median 37-42 %), whereas a large fraction (95-97 %) of AMSC populations from villages 3 and 4 were in a quiescent state (**Supplementary Figure S2**). This observation may be attributed to 1) Differences in donor numbers: villages 1 and 2 each included ~20 donors, while villages 3 and 4 each included ~40 donors. 2) Cohort effects across sub-villages: villages 1 and 2 were constructed from a single donor cohort, whereas villages 3 and 4 included donors from 2-3 different cohorts. It remains unclear whether the observed effect is driven by the number of donors, the inclusion of multiple cohorts, or both. Given that this cell state was recovered from only a limited number of donors and preliminary from village 1 and 2, we excluded it from pseudobulk QTL-related analyses, which require sufficient sample sizes per cell state. In future studies, an ideal design would be pooling a consistent number of donors across multiple cohorts. Aside from the variability in AMSC proliferation across villages, we did not observe other notable biases among sub-villages.

**Donor-level cell-state proportions are reproducible across villages and not driven by technical covariates**

Donor-level cell-state compositions differed across villages (Kruskal–Wallis, BH-adjusted $P < 0.01$ for all states, median adipocyte fraction 49.2, 31.1, 32.7 and 21.5% in villages 1–4). This inter-individual variability was reproduced within every village, with each village spanning a similarly wide range of adipocyte fractions (6.6–69.3, 6.3–57.2, 3.6–59.8 and 1.0–68.2% for villages 1–4), villages differed in the median numbers but not in the breadth of this variation, indicating a reproducible biological characteristic rather than a batch effect. We further assessed three technical covariates.

Per-donor sequencing depth varied across villages (median UMI 4,018–7,599; Kruskal–Wallis $P = 2.5\times10^{-15}$) and was negatively correlated with adipocyte proportion across donors (Spearman $\rho = -0.56$, $P < 2.2\times10^{-16}$). This association was also present within villages ($\rho = -0.08$, $-0.66$, $-0.21$ and $-0.63$ for villages 1–4, significant in villages 2 and 4), indicating it is not merely a between-village effect. Two aspects indicate this reflects biology rather than a depth-driven artifact. First, clustering and annotation were performed on SCTransform-normalized expression, in which sequencing depth was modeled as a latent technical covariate and additionally included as a regression variable, so that cluster assignment and cell state annotation are not directly confounded by per-cell depth. Second, among all cell states, adipocyte nuclei had the lowest sequencing depth (median 5,732 UMI and 2,578 detected genes, versus 6,691–7,400 UMI and 3,003–3,440 detected genes for other states), consistent with the lower RNA content of mature adipocytes. Donors with a higher adipocyte fraction therefore yield lower median per-donor depth. The negative depth–composition association is thus a consequence of composition rather than its cause. Nuclei recovery (the number of nuclei passing quality control) did not differ across villages (Kruskal–Wallis P = 0.20) and therefore did not account for the village-level compositional differences. Demultiplexing was performed using genotype-based assignment, and only confidently assigned singlets were retained; because genotype-based singlet assignment is independent of expression depth or content, it is not expected to bias cell-state composition.

**Donor identity is the dominant source of variance in cell-state composition.**

We partitioned the variance in day 14 cell-state composition using a linear mixed model with donor, stimulation condition and village as random effects (adipocyte fraction logit-transformed as the response; donor nested within village; REML). Donor identity accounted for the majority of variance (65.9%), far exceeding village (14.2%), stimulation condition (12.4%) and residual variance (7.5%). Partitioning the SWAT fraction yielded an essentially identical structure (donor 62.3%, village 17.4%, stimulation 11.2%, residual 9.1%), consistent with the complementary nature of these states. The small village component relative to donor indicates that cross-village differences largely reflect donor-level biological variation rather than independent batch effects.

**Primary GWAS sources for the traits used in scDRS analysis**

GFATadjBMI (Gluteofemoral fat adjusted for BMI), ASATadjBMI (Abdominal subcutaneous adipose tissue adjusted for BMI), VAT/ASAT ratio (Visceral to abdominal subcutaneous adipose tissue ratio): *Agrawal, S., et al. (2022). Nat Commun.*

FI (Fasting Insulin)*: Chen, J., et al. (2021). Nat Genet.*

IFCadjBMI (Insulin Resistance based on Insulin Fold Change adjusted for BMI), ISIadjBMI (Stumvoll's Insulin Sensitivity Index adj. BMI): *Williamson, A., et al. (2023). Nat Genet.*

HOMAIRadjBMI (Homeostatic Model Assessment of Insulin Resistance adjusted for BMI): *Manning, A.K., et al. (2012). Nat Genet.*

BMI (body mass index), WHRadjBMI (Waist-Hip Ratio adjusted for BMI), LDL (Low-Density Lipoprotein), HDL (High-Density Lipoprotein), Triglyceride, HbA1c (Hemoglobin A1c), Glucose: *UK Biobank*

T2D (Type 2 Diabetes): *Suzuki, K., et al. (2024). Nat Genet.*

T1D (Type 1 Diabetes): Bradfield et al. 2011 Plos Genet

CAD (Coronary Artery Disease): *Schunkert et al. (2011). Nat Genet.*

For BMI, WHRadjBMI, LDL, HDL, Triglyceride, HbA1c, Glucose, T1D, CAD traits, we used processed summary statistics from the scDRS data resource, the remaining traits were munged in-house from the original summary statistics using scDRS function “munge-gs”.

### Supplementary figures

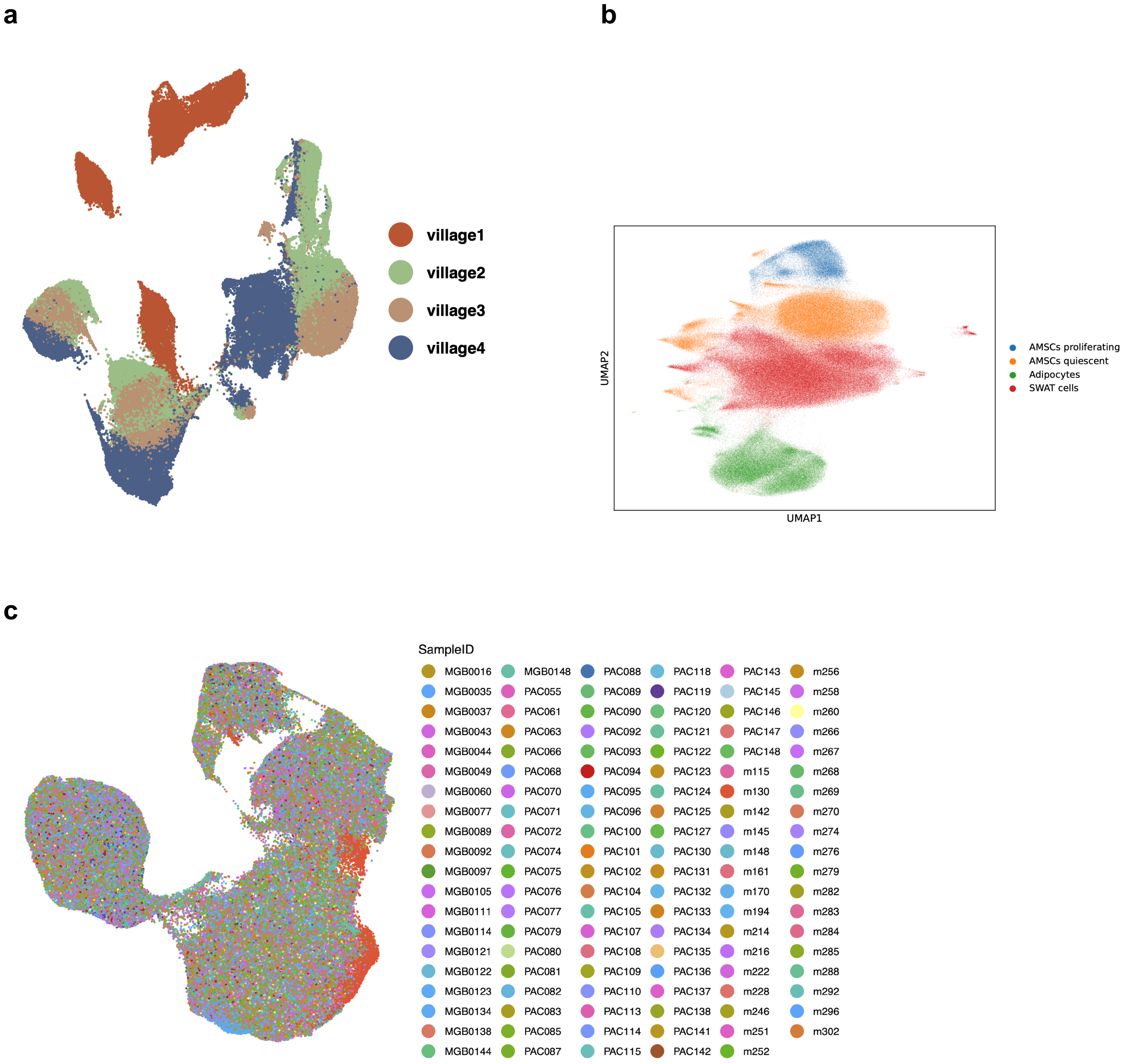


**Supplementary Figure S2. Extended information regarding the batch correction.**

**a,** UMAP colored by village batch before RPCA batch correction. **b,** UMAP colored by donor ID after RPCA batch correction, with iAMSC cell line included.

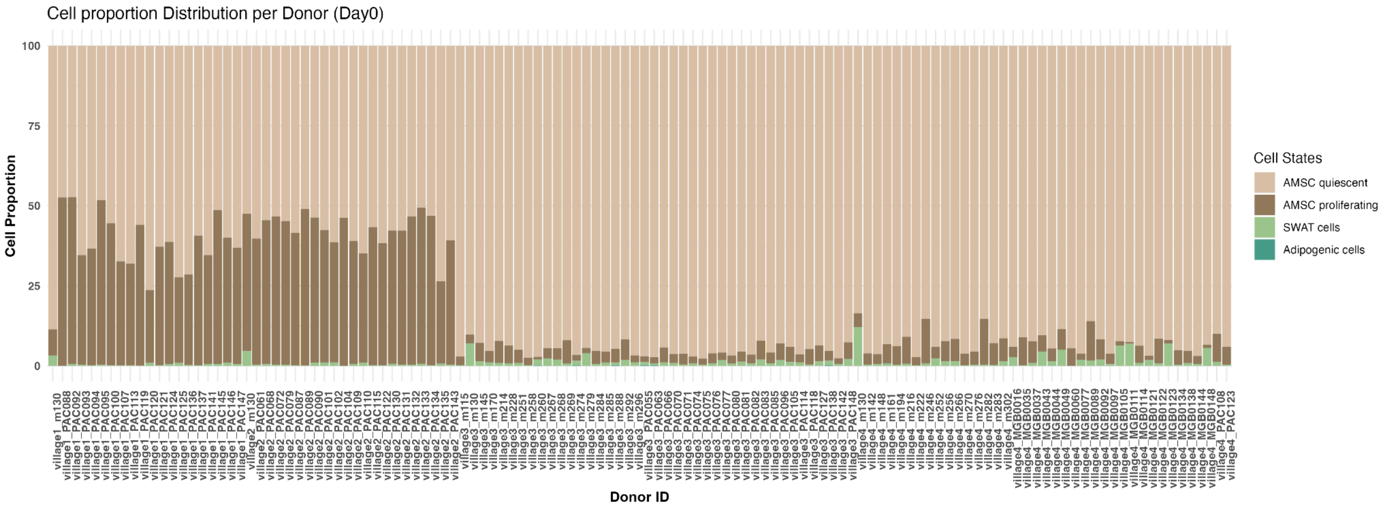


**Supplementary Figure S3. Extended information regarding the AMSC proliferating rate.** Stacked barplot colored by cell states to represent per donor cell state proportion at day 0.
